# Effects of Cellular Memory and Adaptation Cost on Optimal Survival in Fluctuating Environments

**DOI:** 10.1101/2025.05.24.655868

**Authors:** Paras Jain, Mohit Kumar Jolly, Jason T. George

## Abstract

Cells invariably encounter unpredictable changes in their microenvironment and adapt by orchestrating substantial alterations in their molecular states, often resulting in appreciable phenotypic changes. The timescale of molecular adaptation depends on how quickly a cell loses its molecular memory of past environmental encounters through the degradation rate of proteins unfavorable to the current environment. Concurrently, *de novo* synthesis of favorable biomolecules during adaptation imposes an energetic cost that impacts cellular fitness. Here, by developing a phenomenological model of intracellular processing of environmental signals and associated cell-state switching, we study the dynamical implications of cellular memory and adaptation cost on cellular responses to a changing environment. We find that while increasing cellular memory reduces cell fitness in periodic environments, counterintuitively, increasing adaptation cost can improve growth by minimizing mismatches between the environment and the cell state. Similarly, we observed a variable role of memory capacity and adaptation cost for stochastic correlated environments, with increasing memory and cost improving cellular fitness in negatively but not positively correlated environments. Lastly, we show that cellular memory and population heterogeneity in adaptation cost explained reported experimental observations in melanoma: increased population survival during drug treatment when the population was either enriched for rare cells expressing resistance marker genes or primed with low-dose drug treatment before exposure to high-dose treatment. Overall, this work establishes a foundational model for studying how cellular memory dynamics and adaptation cost drive cellular adaptation under different environmental conditions and explain complex cellular behaviors.

## Introduction

Phenotypic adaptation is a defining response and necessity of cellular systems navigating variable environments. Studies across disparate biological contexts, spanning bacterial, yeast, and cancer systems, have extensively characterized cellular phenotypic changes in response to steady and step changes in the environment. Only recently has there been an increasing focus on cellular responses to fluctuating environments (1–5). Environmental fluctuations drive cellular adaptation across many timescales. For example, second-to-minute scale changes in nutrient gradients drive bacterial motility, while hour-to-day scale changes in nutrient source induce distinct cellular growth phenotypes (4–8). It has been shown empirically that fluctuating environments produce cellular responses that can differ significantly from what would be expected from taking a suitable weighted average of responses in static environments (4). Moreover, these responses to dynamic environmental changes result from the orchestration of cellular action based on previous environmental encounters, representing an imprinting that we refer to as ‘cellular memory’ (5, 9, 10). A more complete understanding of cellular behavior therefore necessitates understanding the dynamic cellular responses to fluctuating environmental conditions.

Previous theoretical models have contributed significant insights into the dynamics of cellular responses in changing environments (11, 12). While studying long-term population-level survival under spontaneous versus adaptive phenotypic transitions in stochastic environments, Kussell and Leibler identified a threshold environmental fluctuation rate below which adaptive switching is more beneficial than spontaneous switching (bet-hedging) (13). Subsequent models identified changes in adaptation dynamics depending on whether adaptation was driven exclusively by past environmental encounters (14). Additional studies on cellular adaptation demonstrated that hysteresis in cell-state switching benefits cellular growth in stochastic, but not periodic, fluctuating environments (15). While the above studies have contributed to our understanding of adaptation in dynamic environments, the relationship between intracellular responses to changing environments and their consequent effects on cellular adaptation remains to be elucidated (Figure 1A) (16–18).

**Fig. 1.**
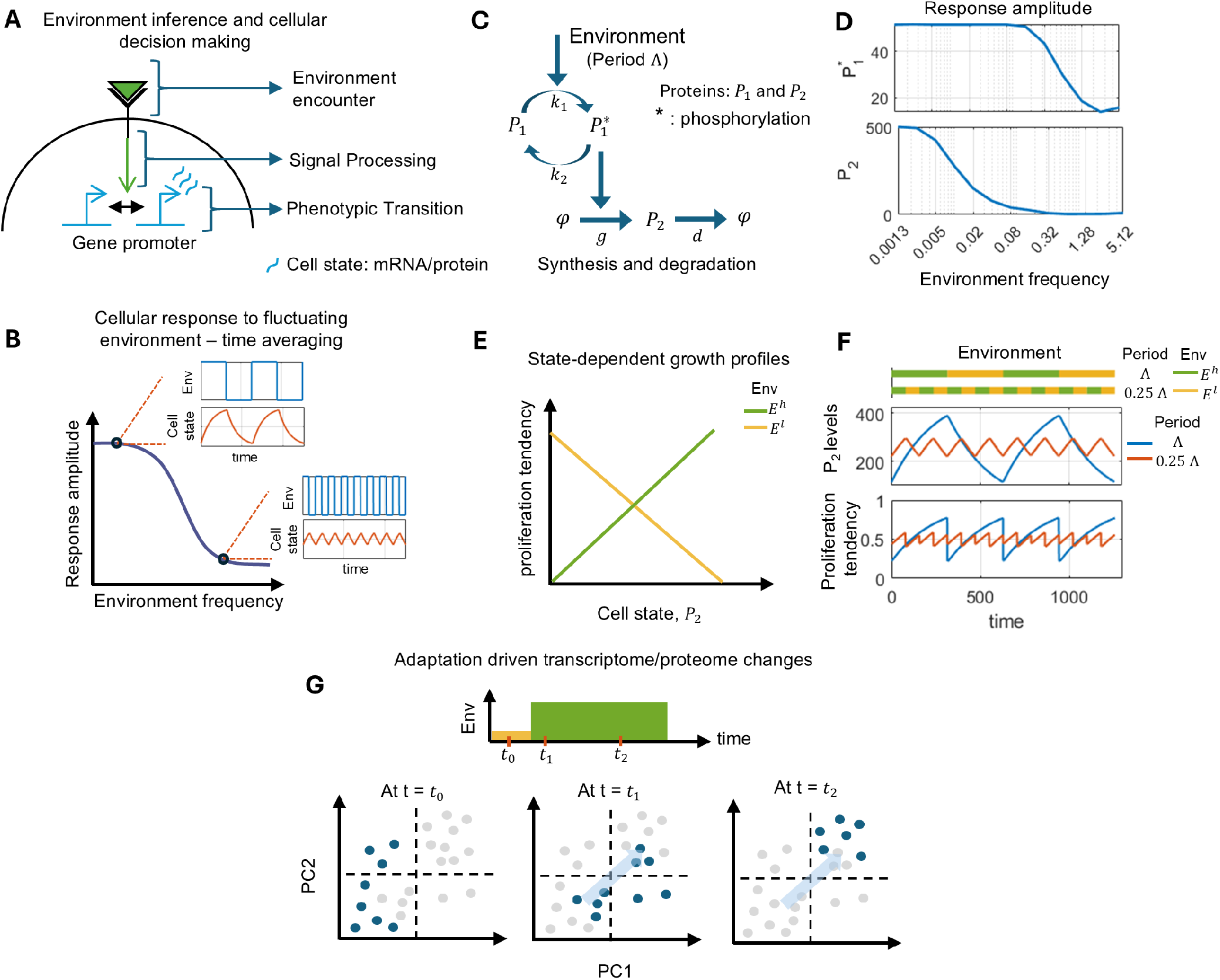
Cellular response in fluctuating environment. **A)** Schematic representation of the intracellular process involved in cellular adaptation in fluctuating environment. **B)** Change in the amplitude of the cellular response with increasing frequency of the oscillatory environment. **C)** Simple intracellular biochemical reaction network mediating cellular response in fluctuating environment. **D)** Frequency response of the intracellular reaction network shown in panel C. It recapitulates the experimentally observed frequency response shown in panel B. **E)** Dependence of cell growth on the abundance of key proteins in a given environment. **F)** Dynamics of cell state and fitness based on cellular response via circuit in panel C and cell state-benefit mapping in panel E. **G)** Schematic representation of a Principal Component Analysis (PCA) plot depicting global changes in gene expression that occur during phenotypic adaptation under the influence of altered environment. Here, each point represents a cell and its (blue) color represent their sampling times before and after the population was subjected to environmental change. Such global changes in the internal state of cells points toward the energetic cost of adaptation due to *de-novo* synthesis of several bio-molecules. The model equations and parameters used to perform quantification in panel C to F are presented in the SI text

Depending on the type of environmental change, one or more conserved intracellular responses are activated. Examples include the expression of Lac operon genes in *E. coli* during nutrient shifts from glucose to lactose and glycerol production in yeast cells upon increase in salinity of the environment (19–22). Importantly, once the environment switches to its prior state, the relaxation time of the biochemical reaction network to its basal levels determines the timescales of cellular memory of the past environmental signals. For example, in *E. coli* cells, dilution of LacY protein levels during subsequent divisions following lactose exposure erases the cellular memory of lactose signals, evident in increased lag and recovery times for cell growth upon lactose re-exposure (Figure 1E, F) (4, 19). Similarly, residual MDR1 levels in cancer cells post-vincristine-pulsed treatment enhanced their survival upon re-exposure to the drug (20). In each case, the precise implications of variability in relaxation times and subsequent memory erasure on cellular adaptation in fluctuating environments remain an open question.

In addition to the timescales of molecular memory affecting adaptation, since cellular maintenance of a particular phenotypic state requires a constant cellular energy influx to maintain persistent synthesis and degradation of favorable biomolecules, there is a trade-off between fitness gains in the current environment and the preparedness for future changes(23–25). Such energetic constraints derived from the production of proteins favorable for unforeseen environments limit cellular adaptation ability, thereby causing them to perform *de-novo* synthesis of essential proteins in response to the inferred environment (Figure 1G) (26, 27). How adaptation costs affect the long-term growth of cells in fluctuating environments is yet to be explored.

Here, we present a phenomenological model of coupling intracellular signaling dynamics with cellular decision-making to study cellular adaptations in fluctuating environments. In our model, intracellular signaling acts to time-average past environmental signals to make an optimal decision of residing in a cell state that maximizes the average fitness of the cell. By developing and applying analytical theory and numerical simulations, we first demonstrate the applicability of the developed framework by recapitulating the cellular adaptation dynamics observed in *E. coli* cells under fluctuating nutrient environments. Next, using the framework, we characterized the effects of relaxation times (cellular memory) and adaptation costs on the cellular response to periodic and stochastic environmental changes. We found that while increasing cellular memory slows adaptation and reduces cellular fitness in periodic environments, increasing adaptation costs can in fact enhance cellular fitness by reducing cellular maladaptation. Similarly, we observed variable roles of cellular memory and adaptation in stochastic correlated environments. A larger cellular memory and adaptation cost result in higher fitness level in negatively correlated stochastic environments than in positively correlated ones. To apply our model, we showed that the presence of cellular memory and population-level heterogeneity in adaptation cost explained the reported experimental observations of enhanced population survival of melanoma cells during cytotoxic drug treatments when 1) the population was enriched for ‘pre-resistant’ cells or 2) the population was primed with a low-dose treatment before getting exposed to high-dose treatment. Overall, our study highlights the non-intuitive roles of cellular memory and adaptation cost in dictating the dynamics of cellular adaptation and consequent fitness in variable environments.

## Model Development

To study how relaxation times of signaling dynamics and adaptation cost influence cellular adaptation, we employed a discrete-time approach to model the environmental changes, their processing by the cell and resulting cellular adaptation.

### Fluctuating environment dynamics

We assumed that at any time instance ‘*n*’ the environment is either a beneficial/high (e.g. nutrient) environment (E-high, *E*^*h*^) or a detrimental/low environment (E-low, *E*^*l*^). In this study we consider the following two dynamics of the discrete *E*^*h*^ and *E*^*l*^ environments:

1. Stochastic environment fluctuations with ℙ (*E*_*n*_ = *E*^*h*^) = *p* and one step autocorrelation of *ρ*_*EE*_(1). The environment dynamics can be captured using the following transition probability matrix:

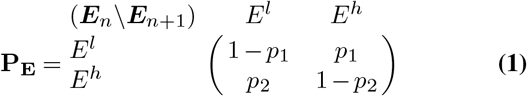

where, *p*_1_ and *p*_2_ represent transition probabilities between environmental states, with *p*_1_ = min(1, *p* (1 − *ρ*_*EE*_(1))) and *p*_2_ = min(1, (1 − *p*) (1 − *ρ*_*EE*_(1))). When *ρ*_*EE*_(1) = 0, we have *p*_1_ = 1 and *p*_2_ = 1 − *p*.
2. Periodic environment fluctuations with period *T* . The environment dynamics at time *n* = *r T* + *l*, where *r* and *l* are integers, is given by:

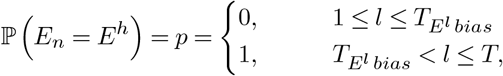

where, 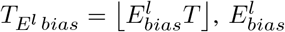 denotes the fraction of one environmental oscillation in the *E*^*l*^ state, and ⌊·⌋ represents the floor function giving the greatest integer less than or equal to its argument.

### Cellular environmental signal processing

One common feature observed across cellular responses to different stressors is that cells follow changes in the environment up to a particular frequency above which the cellular response becomes time-averaged (Figure 1B) (28, 29). Such distinct – periodic and time-averaged – cellular responses to environmental fluctuations can be achieved by a simple biochemical reaction network, based on the relative timescales of signaling (de-)activation or synthesis and degradation of downstream gene products with respect to environment fluctuation timescales (Figure 1C, D). Thus, to capture the above cellular signaling dynamics, while at the same time focusing on studying the influence of its relaxation dynamics on cellular adaptation, we assume that cells have a discrete memory of their past environmental exposures and the memory updates by incorporating more recent environmental signals and losing the previous ones (Figure 2A). Further, as biomolecules commonly exhibit first-order degradation kinetics, we assumed that all previously encountered environmental exposures have an equal probability of being forgotten. Towards this end, at each time step, a random environmental signal encoded in memory is replaced with a new one from the current environment. The degradation rate constants, which determine the responsiveness of the signaling dynamics, are modeled by a finite cell memory capacity ‘*m*’. A smaller memory capacity would then result in more rapid washout of past environmental signals, thereby increasing cellular responsiveness to a new environmental encounter.

**Fig. 2.**
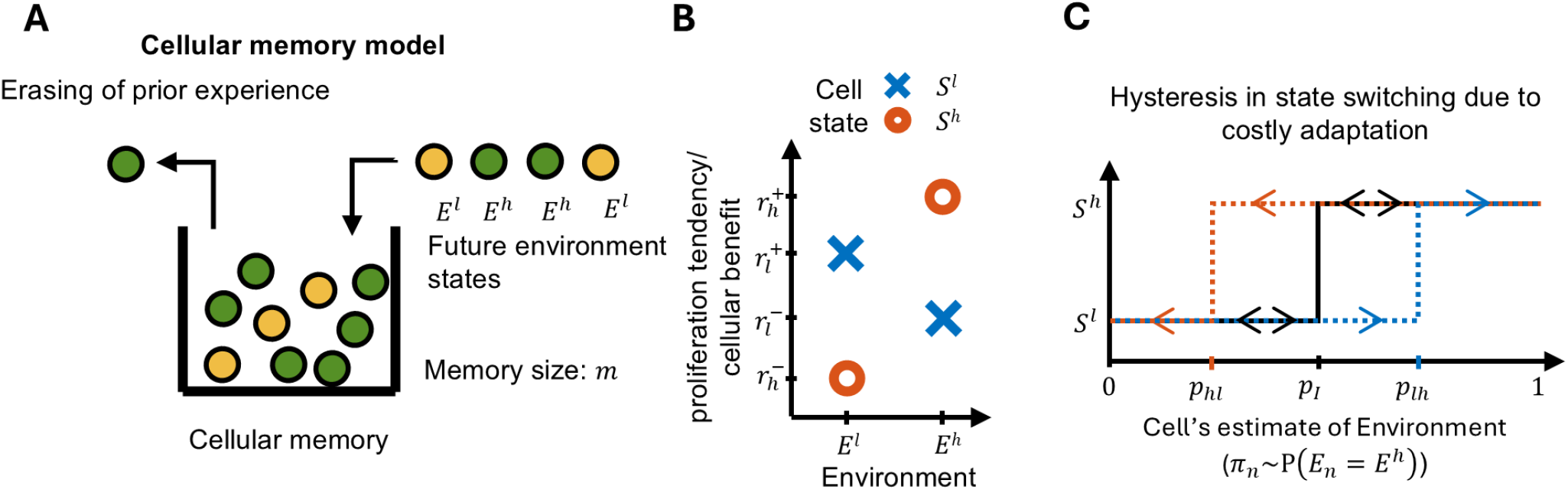
Cell memory model, environment and cell states, and cellular decision making. **A)** Proposed cell memory model to model intracellular signaling dynamics. Consideration that environment can reside in one of the two states *E*^*h*^ and *E*^*l*^. At every discrete time step a random past environmental signal (either *E*^*h*^ or *E*^*l*^) is erased from the cell memory based on its frequency and the current environmental signal is stored. **B)** For the two environments, *E*^*h*^ and *E*^*l*^, we assume there exists optimal cell states *S*^*h*^ and *S*^*l*^ that respectively maximize the fitness. **C)** Points of cellular decision making to reside in either cell state *S*^*h*^ or *S*^*l*^ based on the cell’s estimate of the average environment (*π*_*n*_ ∼ ℙ (*E*_*n*_ = *E*^*h*^)) and maximization of the average fitness (eq. 3, eq. 4). *p*_*I*_ represents indifference environment (ℙ (*E*_*n*_ = *E*^*h*^ ∗)) where residing in either cell state is equally beneficial. If there is a cost associated with phenotypic adaptation, then the above indifference environment, *p*_*I*_, splits into *p*_*lh*_ and *p*_*hl*_ for *S*^*l*^ → *S*^*h*^ and *S*^*h*^ → *S*^*l*^ state transitions, respectively.

Now, for an environment that fluctuates between *E*^*h*^ and *E*^*l*^ states, we use *K*_*n*_ to denote the total number of prior *E*^*h*^ state signals in the cell memory at time *n*. Thus, at each time time, the fractions 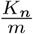 and 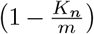 denote the probabilities of losing a *E*^*h*^ or *E*^*l*^ environmental signal from the memory, respectively. Similarly, *p* and (1 − *p*) denote the (independent) respective probabilities of gaining an *E*^*h*^ and *E*^*l*^ environmental signal at each time step, respectively. The signaling dynamics via cell memory updates can be modeled as a time-homogeneous Markov Chain with states *K*_*n*_ ∈ [0, 1, …, *m* − 1, *m*] and state-transition probabilities given by:

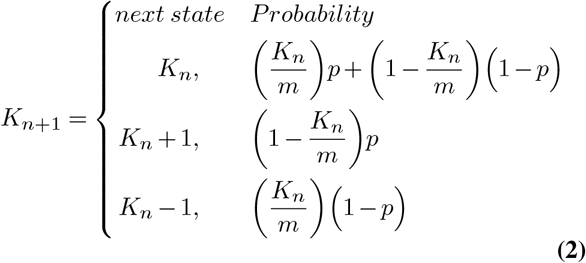

The above state transition probabilities hold for periodic and uncorrelated stochastic environments since in both cases *E*_*n*_ is independent of the previous time step. However, in correlated stochastic environments both *K*_*n*_ and *E*_*n*_ depend on *E*_*n*−1_, and so we define the joint transition probabilities for the environment, *E*_*n*_, and cell state, *K*_*n*_ in *SI Eq. S3* .

### Optimal phenotypic states in fluctuating environments

In this framework the environment is in either *E*^*h*^ or *E*^*l*^ state at any time instance ‘*n*’. We therefore consider two phenotypic states, *S*^*h*^ and *S*^*l*^, which are preferred in environments *E*^*h*^ and *E*^*l*^, respectively, and impart greater fitness to the cell. Fitness in our model is indicated by 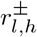, with the subscript indicating the cell state *S* and the superscript denoting whether the cell state *S* matches (+) or mismatches (−) with the environment *E*. For example, a proper match of cell state *S*^*l*^ with environment *E*^*l*^ (resp. *S*^*h*^ with *E*^*h*^), results in an enhanced fitness 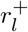 (resp. 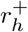). However, a mismatch *S*^*l*^ with *E*^*h*^ (resp. *S*^*h*^ with *E*^*l*^) results in a reduced benefit 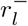 (resp. 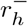) (Figure 2B).

Cells can switch between *S*^*h*^ and *S*^*l*^ states based on the memory of past environmental signals to maximize their average fitness in future environments. Furthermore, as cell state switching may involve substantial changes in multiple biomolecules at the transcriptome/proteome levels (Figure 1G), we also incorporate adaptation costs *c*_*lh*_ and *c*_*hl*_ for transitions *S*^*l*^ → *S*^*h*^ and *S*^*h*^ → *S*^*l*^, respectively.

Letting *S*_*n*_ denote the cell state at time *n*, the fitness of the cell is given by:

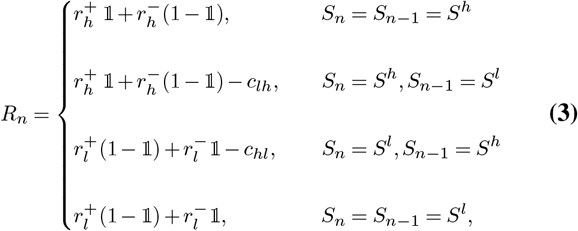

where, 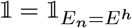 and denote the usual indicator random variables:

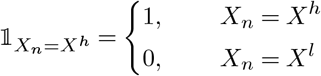

The differential fitness accrued by each environment-phenotype pair impart a phenotypic preference for each environmental state that depends on *p*. This preference partitions the environmental landscapes into two groups separated by a critical indifference environment for which each phenotype is equally beneficial. This indifference environment can be calculated by equating the expectations of each pair of values in Eq. (3) having the same phenotype at the previous time step *S*_*n*−1_ (See SI section S4 for full details). In particular, we have

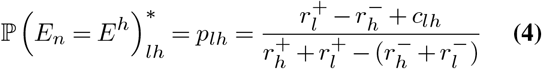

In the event that the previous state of the cell is *S*_*n*−1_ = *S*^*l*^, and ℙ (*E*_*n*_ *E < p*_*lh*_, then *S*_*n*_ *S* is more beneficial than *S*_*n*_ = *S*^*h*^. Otherwise, *S*_*n*_ = *S*^*h*^ is more beneficial (Figure 2C).

Similarly, the indifference environment, *p*_*hl*_, for *S*^*h*^ to *S*^*l*^ transition can be quantified by replacing *c*_*lh*_ with −*c*_*hl*_ in eq. Eq. (4). Therefore, when *S*_*n*−1_ = *S*^*h*^, and ℙ(*E*_*n*_ = *E*^*h*^ *) > p*_*hl*_, then *S*_*n*_ = *S*^*h*^ is more beneficial; otherwise *S*_*n*_ = *S*^*l*^ is more beneficial (Figure 2C).

### Memory-driven cellular inference of the environment

To reside in a optimal state that maximizes average fitness the cell uses its memory of *E*^*h*^ environment signals, *K*_*n*_, over the previous *m* time instances to estimate the environment probability, *p*. The estimate *π*_*n*_ of *p* is determined by the relative frequency of *E*^*h*^ signals in the cell memory:

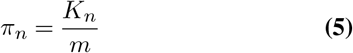

### Cellular decision making in fluctuating environments

Cells use *π*_*n*_ as an estimate of the likelihood of a future *E*^*h*^ state in order to maximize their fitness at the next time step (Figure 2C):

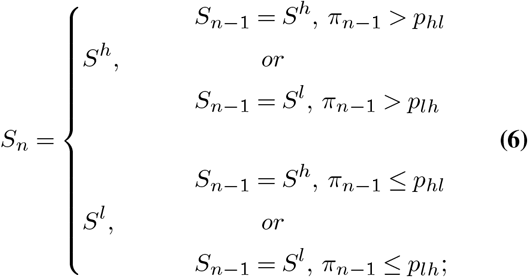

We note that when transitions bear no cost, the decision making points of indifference agree (*p*_*lh*_ = *p*_*hl*_ = *p*_*I*_). Positive costs results in strict inequality *p*_*lh*_ *> p*_*hl*_, causing hysteresis in cell state switching, i.e., environmental conditions for state switching are different for *S*^*l*^ → *S*^*h*^ and *S*^*h*^ → *S*^*l*^ transitions (Figure 2C).

### Quantification of average fitness

From Eq. 3 we can write down the cell fitness conditional on phenotypic state as follows,

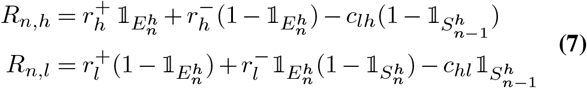

where, 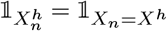.

The conditional expected benefits at time *n* are then given by:

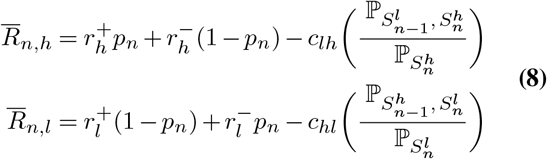

where,

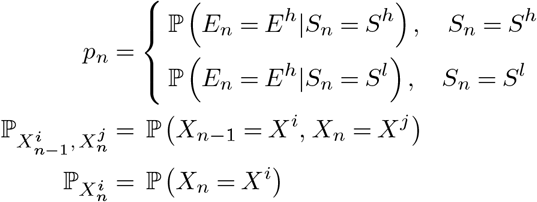

And, the unconditional average fitness is:

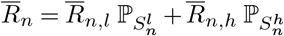

Detailed quantification of *p*_*n*_, 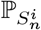, and joint probabilities 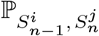 are given in the *SI section S8*.

### Population dynamics

So far, we have described how cell memory updates in time, how this memory is used as an estimate of future environments to chose an optimal cell state, and the relation between environment-state (mis-)match and cell fitness. If we consider a population of cells, with each cell making an independent decision to reside in either state based on its memory, then we could capture the dynamic change in the mean population size using a discrete time Markov chain model by adapting the method developed in Devaraj and Bose (30) as follows:

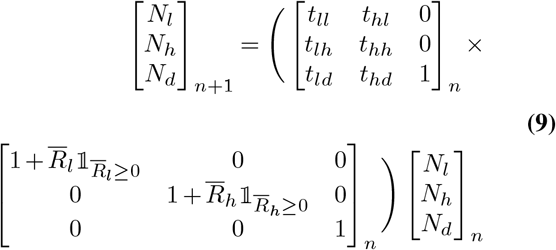

where, *N*_*i*_’s with *i* ∈ {*l, h, d}* are the mean number of cells in states *S*^*l*^, *S*^*h*^, and death, ‘*d*’, cell compartment; *t*_*ij*_’s with *i, j* ∈ { *l, h, d}* are the transition probabilities between states *S*^*l*^, *S*^*h*^ and death cell compartments; 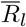 and 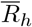 are conditional expected fitness values for residing in cell states *S*^*l*^ and *S*^*h*^, respectively (Eq. 8) and define the fraction of cells in each state that divide in a step time. The elements of the above state transition matrix must follow the below inequalities:

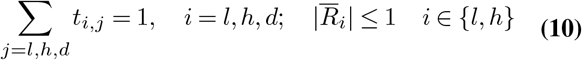

The quantification of the transition probabilities *t*_*ij*_ and conditional fitness 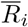 are described in *SI section 10*.

Average per capita growth rate of the population is given as:

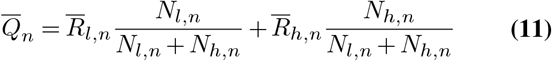

### Choice of parameter values

For the evaluation of impact of memory size and adaptation cost on average fitness in periodic and stochastic environments, we have considered relative (dimensionless) fitness of the cell state on its (mis-)match with the environment, 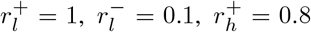, and 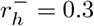. The above parameter choice results in *p*_*I*_ = 0.5, and represents a cytostatic (growth reduction) response to cell state-environment mismatch.

For studying population dynamics under a sudden switch to a lethal environment, we chose dimensional value 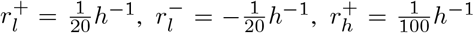, and 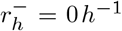, such that they resemble division and death timescales of melanoma cancer cells as we associate the observation made with them with our findings (9, 31). The above parameter choice represents a cytotoxic (cell death) response resulting from mismatch between environment and cell state, and results in *p*_*I*_ = 0.45.

Cases having non-negligible adaptation cost are assumed to be symmetric with respect to navigating low-to-high and high-to-low environments (i.e., *c*_*lh*_ = *c*_*hl*_). Adaptation costs are sampled from the interval 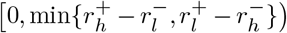 where 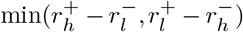 gives the maximum adaptation cost at or beyond which either *p*_*lh*_ ≥ 1 or *p*_*hl*_ ≤ 0. Adaptation costs have the same dimensional units as fitness parameters (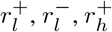, and 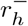).

## Results

### Cell memory dynamics recapitulate observed adaptation in periodic environments

The proposed model and the mapping of the growth benefits to the cell state are intended to capture the time-averaged response of intracellular biochemical networks and the associated changes in the dynamics of cell growth in fluctuating environments (Figure 1B, C, F). To test the model’s validity, we first checked whether it was able to qualitatively match the time-averaged environmental response of *E. coli* cells experiencing nutrient shifts between glucose and lactose carbon sources (4, 19). For this, we considered environmental oscillations between *E*^*h*^ and *E*^*l*^ states with periods of 10 and 40. We observed changes in the cell’s estimate of the environment (*π*_*n*_), the cell state (*S*_*n*_), and the fitness over time. We found that oscillating environments with shorter periods led to lower amplitude fluctuations in *π*_*n*_ and *S*_*n*_ when compared to those longer periods. These reduced fluctuations in *π*_*n*_ resulted in faster recovery of fitness whenever there was a change of environment (Figure 1A). However, such responsiveness in shorter periods also came with a tradeoff in reduced average fitness over one complete environmental cycle when compared to environment osculations having longer periods (dashed red lines in Figure 1 Aiii). These cell state and growth dynamics recapitulate those of the LacY protein and its associated changes in growth rate in *E. coli* cells across different environmental conditions (19). In the analysis above, we also observe an excellent agreement between the numerical simulations and our theory.

Given that the time-averaged cellular responses in periodic environments agreed with empirical observations, we next sought to understand how changes in memory capacity and adaptation cost influence cellular adaptation.

### Increasing memory delays adaptation and reduces fitness in periodic environments

Increasing memory capacity represents, for example, a reduction in the rate of post-transcriptional modifications of signaling proteins or the degradation of proteins that determine a phenotypic state. Thus, larger memory capacities reduce cellular responsiveness to high-frequency oscillations, thereby limiting phenotypic adaptation (Figure 1B, D). Indeed, on increasing memory size for an oscillating environment with fixed period, we observed reductions in the mean fitness over each period. These fitness reductions arise as a consequence of increased cellular inertia to state changes on environmental change, which result in environment-phenotype mismatch (Figure 3B, C, Figure S1A) (see *Model Development section Quantification of average fitness* for the theoretical calculation). Further, cellular responses to shorter periods were time-averaged more than those for longer periods for the same memory size (Figure 2A), which explained why longer periods conferred a greater fitness benefit than shorter periods across all memory sizes (Figure 3B). Next, we evaluated the influence of cellular memory on fitness in biased periodic environment, with 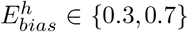 and denote the fraction of one environmental oscillation in *E*^*h*^ state. We found a non-monotonic change in average benefit over an cycle with increasing memory size, especially for smaller periods. The benefit first decreases for small-to-intermediate memory sizes (*m* ∼ [2, 20]) and then increases for large memory sizes (Figure S2A). However, the benefits saturate at large memory sizes, with reducing differences between environmental periods.

**Fig. 3.**
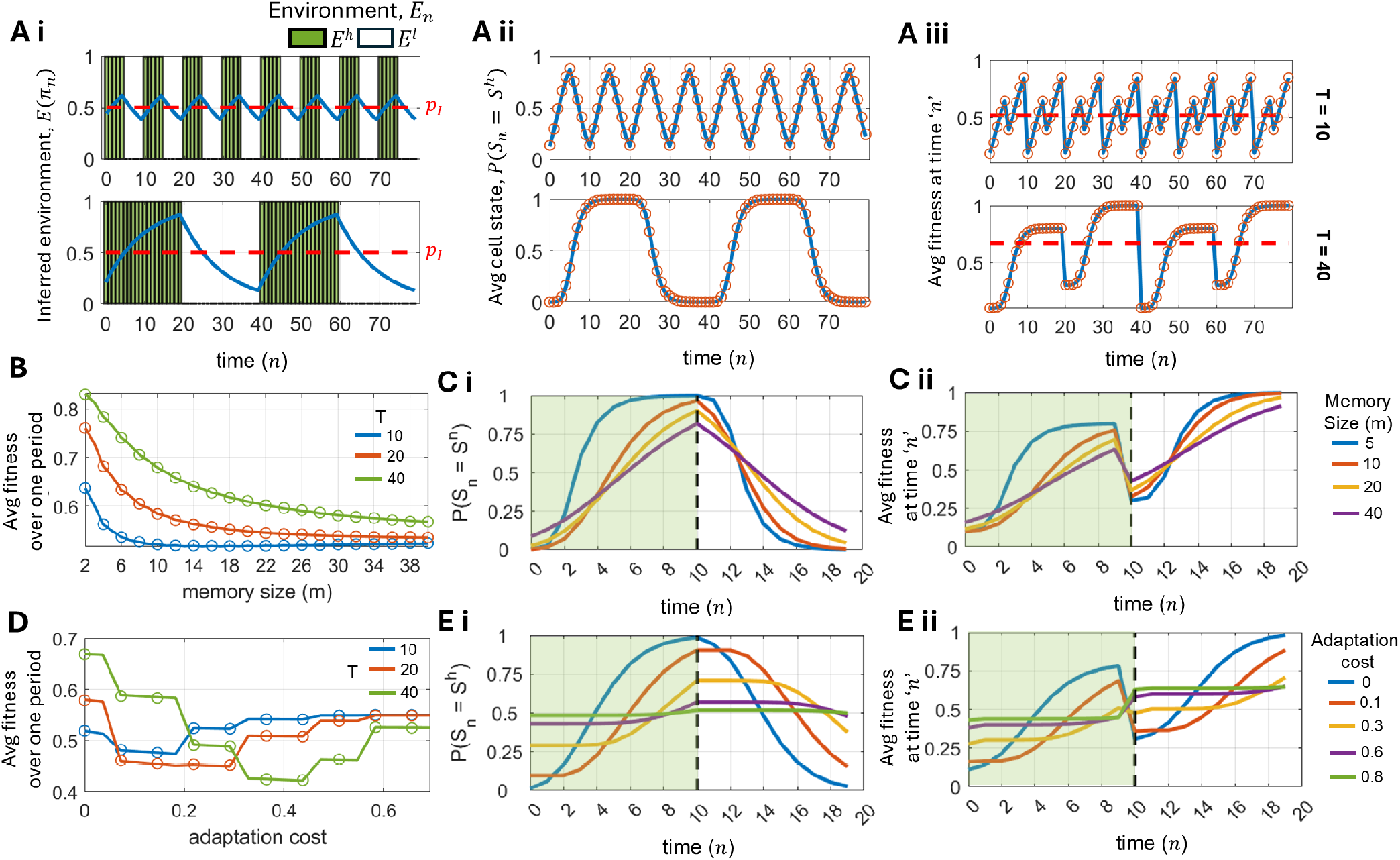
Cellular adaptation to periodic changes in the environment. **A)** Dynamics of (i) cell’s inference average environment, *π*_*n*_), (ii) probability of *S*^*h*^ cell state, and (iii) average fitness in periodic environments of period 10 and 40. In panel Ai, red dashed line denote the indifference probability, *p*_*I*_ ; and in panel Aiii red dashed line denote the average fitness over one period. **B)** Changes in the average fitness over one complete cycle with increasing memory size. **C)** Dynamics of cell state and fitness during an environmental cycle of period 20 for increasing memory size. Here, the environment is in the *E*^*h*^ at times left of the dashed line and in the *E*^*l*^ state subsequently. **D)** Changes in the average fitness over one complete cycle with increasing adaptation cost. **E)** Dynamics of cell state and benefit during an environmental cycle of period 20 for increasing adaptation cost. Here, the environment is in the *E*^*h*^ at times left of the dashed line and in the *E*^*l*^ state subsequently. In all the analysis above, solid lines represent analytical results with their corresponding numerical simulation shown by circles, ‘o’; memory size, *m* = 11 and adaptation cost, *c*_*lh*_= *c*_*hl*_ = 0, unless stated otherwise; 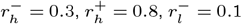, and 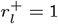..

### Large adaptation costs prevent maladaptation by minimizing phenotypic switching

Increasing adaptation costs are associated with greater molecular-level differences between the two optimal phenotypes (*S*^*h*^ and *S*^*l*^), which impose an energetic burden on the *de novo* synthesis of key proteins during cell-state transitions in response to environmental changes. Therefore, increasing adaptation costs slows down or even restricts cellular adaptation, as reflected in the altered indifference points for transitioning between cell states Eq. (4). Upon switching from low-to-intermediate costs, we observed a reduction in average fitness for all environmental cycles (Figure 3D) (see *Model Development section Quantification of average fitness* for the theoretical calculation). Such a reduction in fitness is due to delayed transition to the optimal cell state in the current environment cycle (e.g. ℙ (*S*_*n*_ = *S*^*h*^*) <* 0.5 for most of *E*^*h*^ environment cycle with cost 0.3 in Figure 3Ei, Figure S1B), leading to maladaptation (environment-phenotype mismatch). However, as state transitions become increasingly restricted with increasing adaptation costs, there are fewer maladaptation events than with intermediate costs, which, in fact, results in an increase in average fitness (Figure 3D, Figure S1B). Further, the minimal fitness values occur at different adaptation costs, depending on the period of the environmental oscillations. Similarly, in biased periodic environments, we observe period-dependent non-monotonic changes in fitness with adaptation costs, with higher costs always being more beneficial in fast-oscillating environments (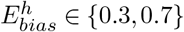 in Figure S2B). This finding suggests that for heterogeneous cell populations with variable costs, subpopulations with lower costs have a relative fitness advantage in slowly oscillating environments. However, this feature becomes disadvantageous in rapidly oscillating environments. Thus, the environmental period can shape the phenotypic distribution along adaptation costs within the cell population over time.

Overall, by studying periodic shifts in the environments, we observe that having smaller cellular memory and lower adaptations costs enable faster cell state switching and improve cellular adaptability.

### Correlation of stochastic environments determines optimal memory size and adaptation cost

The previous subsections detailed cellular responses to controlled periodic oscillations in the environment. However, naturally occurring environments change stochastically, broadly defined by underlying distributional or correlation characteristics. Thus, correlated stochastic signals represent an important class of environments that cellular systems encounter and must effectively navigate.

In an uncorrelated stochastic environment, the probability distribution of cell inference of the environment, *π*_*n*_, follows a binomial distribution with mean *p* and variance *p*(1 − *p*)*/m* (Figure 4A *ρ*_*EE*_(1) = 0). Correlated environments result in a distinct *π*_*n*_ distribution. While negatively correlated environments constrain the *π*_*n*_ distribution around its mean *p*, positively correlated environments widen *π*_*n*_ distribution, with higher probabilities around *π*_*n*_ = 0 and = 1. These distributional changes arise from reduced temporal variation in time-averaged *π*_*n*_ values in fast-fluctuating environments and large temporal variation in *π*_*n*_ values in slowly fluctuating environments.

**Fig. 4.**
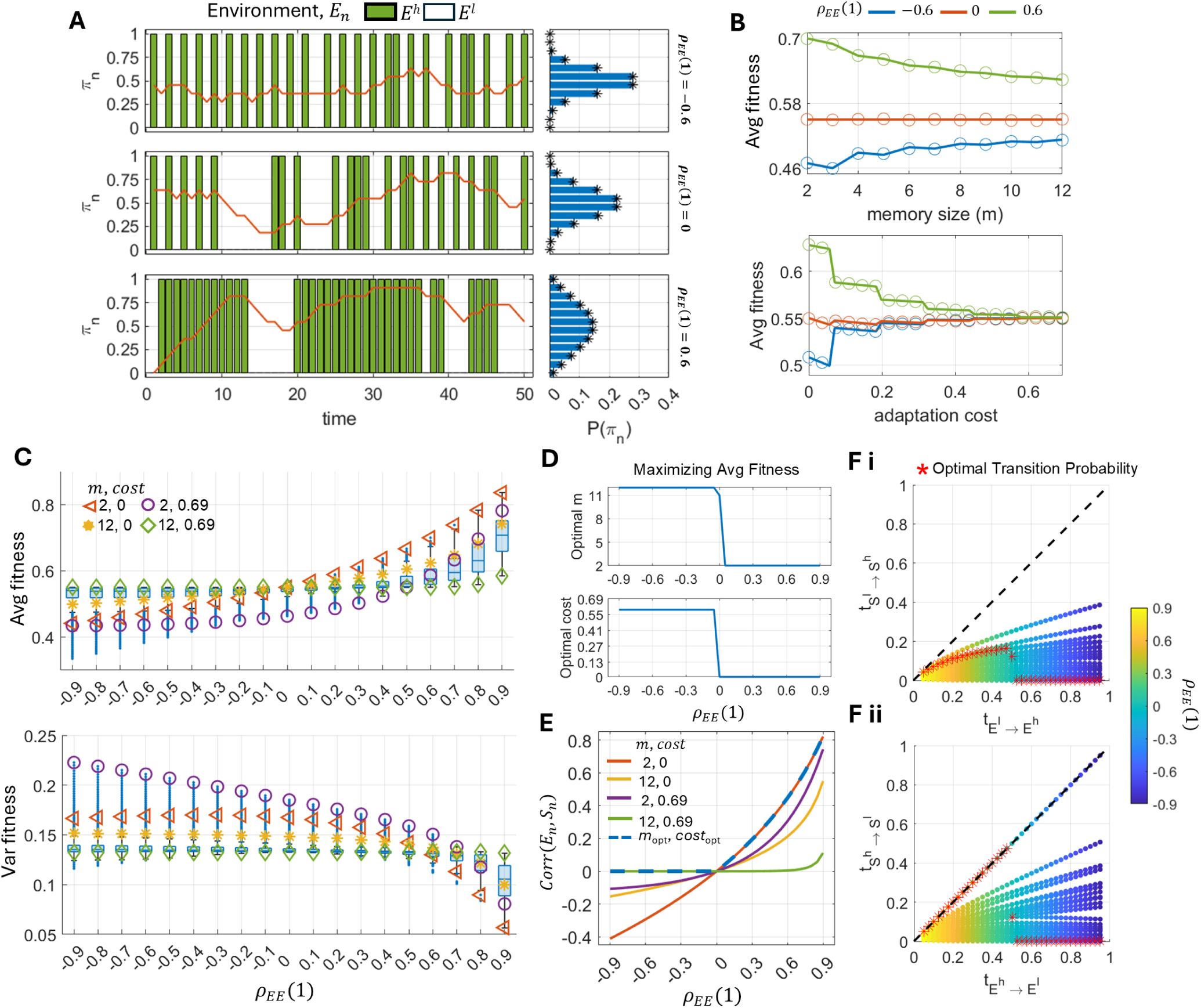
Cellular adaptation to correlated stochastic environmental fluctuations. **A)** Sample trajectories of the environment, and its cellular estimate *π*_*n*_ are drawn for increasing environmental correlation. The stationary distribution of environment estimate, P (*π*_*n*_), for different environment correlations are shown using bars. Numerical validation of the analytical distribution is shown by astericks, ‘∗’. **B)** Changes in average fitness for increasing memory capacity and adaptation cost for different environmental correlation. Numerical validation of the analytical quantification is shown by circles, ‘*o*’. **C)** Changes in the mean and variance of fitness with environmental correlation. The distributions for each correlation value are obtained by all possible combinations of *m* ∈ [2, 12] and 50 values of adaptation cost linearly sampled from the range *c*_*lh*_ = *c*_*hl*_ ∈ [0, 0.69]. Average fitness trajectories from the extreme combination values (*m, c*) ∈ {{2, 0}, {2, 0.69}, {12, 0}, {2, 0.69}} are highlighted with specific markers. **D)** Changes in the optimal memory size and cost that maximize average fitness for increasing environmental correlation. **E)** Cross-correlation between the current environment (*E*_*n*_) and cell state (*S*_*n*_) with increasing autocorrelation in the environment, *ρ*_*EE*_ (1). The dashed blue curve highlights the correlation obtained from optimal (*m, c*) obtained from panel B. **F)** Changes in the cell state transition probabilities (i) 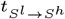 and (ii) 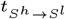 with environmental transition probabilities, 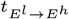 and 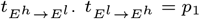 and 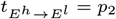 are obtained based on the fixed environmental correlation value (Eq. 1). Scatter points are combinations of transition probabilities obtained from all possible combinations of *m* ∈ [2, 12] and 50 values of adaptation cost linearly sampled from the range *c*_*lh*_ = *c*_*hl*_ ∈ [0, 0.69] with environmental correlation, *p*_*EE*_ (1) ∈ [−0.9, 0.9] having a step size of 0.1. Optimal combinations of environment and cell state transition probabilities for increasing environmental correlation are denoted by astericks, ‘∗’. In the above quantification, 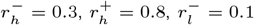 and 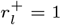; and ℙ (*E*_*n*_ = *E*^*h*^ )= *p* = 0.5.; adaptation cost (*c*_*lh*_ = *c*_*hl*_ = 0) and memory capacity *m* = 11, unless stated otherwise.

Next, we analyzed the effect of memory capacity and adaptation costs in cellular adaptation for cells navigating environments with distinct correlations, using both analytical and numerical approaches (*Model Development section Quantification of average fitness*). We found that the fitness, given a memory capacity and an adaptation cost, depended strongly on environmental correlation. With an excellent agreement between the theory and simulations, we observed that while increasing memory size or adaptation cost conferred higher average fitness values in negatively correlated (highly fluctuating) environments, they reduced fitness for positively correlated (slowly fluctuating) ones (Figure 4B). Further, increasing environmental correlation (from negative to positive) provide a higher average fitness with a lower variance (Figure S4A). Thus, on comparing theoretical estimates of average fitness resulting from a range of memory sizes *m* ∈ [2, 12] and adaptation costs *c*_*lh*_ = *c*_*hl*_ ∈ [0, 0.69], we observed – 1) a general trend of increasing average fitness levels with concomitant reductions in its variance with environmental correlation, and 2) decrease in the optimal memory size and adaptation cost that maximized average fitness with increasing environmental correlation (Figure 4C, D). Such changes in optimality occur because in slowly fluctuating (positively correlated) environments a smaller memory size with lower adaptation cost results in more rapid adaptation and a higher positive correlation between environment and cell state correlation (Figure S4B – quantification of residence times in cell states (32), see *SI section S9* for calculation of *Corr*(*E*_*n*_*S*_*n*_)). In contrast, rapid state switching proves detrimental in highly fluctuating (negatively correlated) environments as the adapted state may not be optimal for the upcoming environment, leading to environment states mismatches and higher negative correlation between environment and cell state and greater cell (Figure 4E, Figure S4B – quantification of residence time in cell states). We found that a larger memory capacity and a higher adaptation cost reduce cell-environment state mismatches in a negatively correlated environment and thereby confer a higher fitness. The above changes in cross-correlation between the environment and cell state – driven by environmental fluctuations – stem from differences in the transition probabilities of the environment and cell state (see *SI section S7* for calculation of cell state transition probabilities). For optimal memory size *m* and adaptation cost-maximizing fitness, we observe almost no cell-state transitions in a negatively correlated environment, whereas cell-state and environment transitions align in a positively correlated environment (Figure 4F).

While analyzing fitness changes in biased stochastic correlated environments (*P* ∈ (*S*_*n*_ = *S*^*h*^) = *p ∈ {*0.3, 0.7}), we find similar roles of memory size and adaptation cost fitness in cellular adaptation as observed above for the unbiased environment *p* = 0.5 (Figure S5). We again observed a general trend of increasing average fitness and decreasing variance with increasing environment correlation, along with optimality of large memory and high cost to maximize average fitness in negatively, but not positively, correlated environments (Figure S6 and Figure S7).

Overall, in a stochastic environment, the nature of environmental correlation determines the effects of memory capacity and cost on cellular adaptation. In the analyses to follow, we present reported experimental observations on cellular adaptation to cytotoxic drug treatments in melanoma cancer cells. We examine these observations in the context of our model and show how the presence of cellular memory and adaptation costs modulates the overall population survival in cytotoxic environments.

### Pre-resistant cells with low adaptation costs exhibit improved adaptation in lethal environments

Recent single-cell characterization of drug-naïve BRAF mutant melanoma cells found a small subpopulation of cells that expressed a few resistance marker genes (e.g., EGFR and AXL) above the population average value. These drug-naïve resistance marker-expressing cells, or ‘pre-resistant’ cells, had a greater likelihood to develop a fully resistant state on treatment with vemurafenib relative to the remaining ‘sensitive’ cells of the population, although they had a similar proliferation capacity under normal growth conditions. One can consider the above existence of two (or more) subpopulations among the drug naïve cells with varying resistance propensities as subpopulations with distinct adaptation costs since pre-resistant cells are phenotypically closer to the resistant cell state than the majority of sensitive cells in the population, and thus, have lower relative adaptation cost. Further, the experimentally observed similarity in fitness between pre-resistant and sensitive cells in constant environments persists with distinct adaptation costs. Thus, we next examined how heterogeneity in adaptation costs affects the dynamics of the overall population under sudden and sustained exposure to the lethal drug. We assume that cells possess a non-negative growth rate when mismatching with the *E*^*l*^ environment 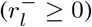, but have a high negative growth rate when mismatching with the *E*^*h*^ environment. This latter condition represents the response of sensitive and resistant cells to a lethal drug. Moreover, to account for heterogeneity in adaptation cost, we consider cells in a population having low (*c*_*lh*_ = *c*_*hl*_ = 0.025) and high (0.05) adaptation costs with a distribution of 0.04% and 99.96%, respectively, capturing the experimentally observed rarity of pre-resistant cells in the parental population.

We analyzed dynamic changes in growth rate and the mean cell counts of individual low- and high-cost subpopulations resulting from step changes in the environment with the addition of drug (environment *E*^*l*^ for time *<* 0; environment *E*^*h*^ for time ≥ 0) using the theoretical results as described in *Model Development section Population Dynamics* and *SI section S10*. Both subpopulations have the same per-capita growth rate before environmental shifts, and consequently they maintain their phenotypic composition under net growth in this condition (Figure 5A, Bi). Upon a sudden switch to the cytotoxic environment, cells in both subpopulations die proportionally until around 40 *h*. Low-cost subpopulation adapt first to the lethal environment and begin to grow; however, because the most abundant cells high-cost cells continue to die due to their delayed adaptation, total cell counts decline persistently. Eventually, the high-cost cells also adapt, ultimately leading to a positive growth rate by the end time point. Consistent with the empirical observation of increased resistant colonies in populations enriched for pre-resistant cells (9), we also observed that population enrichment with cells having low adaptation costs (100% of cells with 0.025 cost) led to notable increase in total cell counts relative to heterogeneous population (0.04% and 99.96% distribution in low and high costs, respectively) after 7 days of drug exposure (Figure 5Bii). Moreover, the enhanced survival of enriched population relative to heterogeneous population persisted across variable memory sizes, with larger memory sizes leading to larger difference in their survivability (Figure S9).

**Fig. 5.**
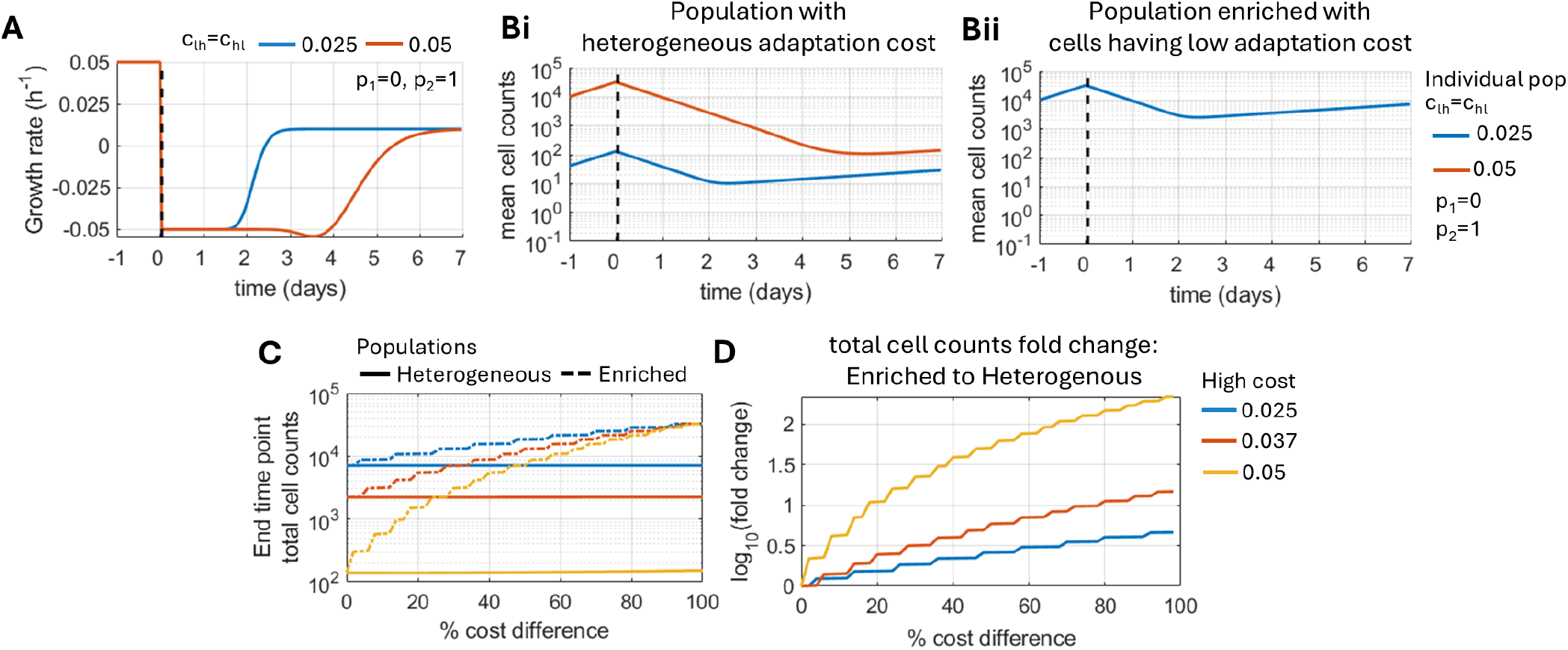
Pre-resistant cells with lower adaptation cost enhance population survival to cytotoxic environments. **A)** Dynamic changes in average fitness of the two subpopulations with distinct adaptation costs (0.025, 0.05) upon encountering a sudden change in the environment (*E*_*n*_ = *E*^*l*^ for time *<* 0 to *E*_*n*_ = *E*^*h*^ for time ≥ 0). During time *<* 0, the two subpopulations had same per-capita growth rates **B)** Dynamic of mean cell counts of subpopulations with distinct adaptation costs to sudden environmental change. The initial proportion of two subpopulations with costs 0.025 and 0.05 were (i) 0.04% and 99.96%, respectively, for the heterogeneous population, and (ii) 100% and 0%, respectively, for the low cost enriched population. **C)** Influence of variation in adaptation cost of low-cost cells relative to high-cost cells on the size of enriched and heterogeneous populations by day 7 of drug exposure. Here, the blue, red, and yellow curves represent the results for the three fixed and distinct adaptation costs of the high-cost subpopulation; the heterogeneous population is composed of 0.04% and 99.96% low- and high-cost cells, respectively. **D)** Fold increase in total cell count of enriched population relative to heterogeneous population at day 7 of drug exposure for increasing difference between adaptation cost of low- and high-cost subpopulations. In the above, 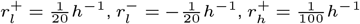, and 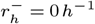; and m = 40. The initial population size was 10^4^ cells, and the population grew in an *E*^*l*^ environment (*p*_1_ = 0) for 24h before switching environments at time 0.

Since the relative reduction in cost for between pre-resistant and sensitive cells is unknown, we also explored the model’s behavior for multiple values of high adaptation cost ({0.025, 0.037, 0.05}) while varying the relative cost of the low-cost cells, in order to quantify their influence on the total cell counts and fold increase in population size with enriched population relative to a heterogeneous population by day 7 of drug exposure. We found that while reducing the absolute adaptation cost of the high-cost subpopulation led to enhanced survival (larger total cell counts) of the heterogeneous population, their survival was largely insensitive to variation in the adaptation cost of the low-cost cells relative to high-cost cells, an outcome of a very low percentage of low-cost cells in the population (0.04%; Figure 5C). However, we found a significant increase in the survival of the enriched population, with a relative reduction in the adaptation cost of low-cost cells, especially more when the reduction is from larger values (high cost: 0.05; Figure 5C). Thus, while increasing the relative differences in costs, we observed an increase in the fold change in total cell counts in the enriched population relative to the heterogeneous population, and this effect was more prominent for larger adaptation costs in highcost cells (Figure 5D).

Overall, we can attribute the extent of molecular similarity between the drug-naïve and drug-resistant cellular phenotypes to heterogeneity in adaptation costs, with the preresistant cells (having higher adaptability) exhibiting lower adaptation costs than the rest of the drug-naïve population.

### Cellular memory enhances survival in step-wise changing environments

In the previous section, we analyzed the population dynamics resulting from the sudden and continued presence of a cytotoxic environment (ℙ (*E*_*n*_ = *E*^*h*^) = *p*_2_ = 1 for time ≥ 0), which was used to represent constant treatment with a high drug concentration. However, multiple studies have observed that cell populations often exhibit enhanced survival when faced with stepwise escalation of drug concentration relative to those with direct exposure to high drug concentrations (9, 33, 34). For example, priming melanoma cells with a low dose (1nM) of trametinib for 2 weeks prior to treatment with a high dose (5nM) increased cell survival and led to more resistant colony formation. Although a multitude of complex processes could regulate the above history-dependent behavior of cells, we tested whether the proposed phenomenological model of cellular memory is sufficient to explain the effects of cellular priming with a low drug concentration on the population’s long-term response to high-concentration treatments. To model less lethal environmental exposures, we considered *E*^*h*^ environments to occur with ℙ (*E*_*n*_ = *E*^*h*^) = *p*_2_ *<* 1 with zero autocorrelation, *ρ*_*EE*_(1) = 0. Here, an uncorrelated stochastic environment is used as it avoids environmental fluctuations associated with negative autocorrelation and long periods of drug presence and absence due to positive autocorrelation (Figure 4A).

We first analyzed the dynamic changes in growth rates and population size as the environment switched from ideal conditions (*p*_1_ = 0) to increasing average lethality (larger *p*_2_ values) (using theoretical results from *Model Development section Population Dynamics* and *SI section S10*). We found that for *p*_2_ ≤ *p*_*I*_, cells have a positive average growth rate at all times and the population continued to expand, albeit with a slower rate, even after the environmental switch. However, for *p*_2_ *> p*_*I*_, a sudden change in the environment led to negative average growth rates, which then recovered to higher levels as cells in the population adapted to the new environmental condition (Figure 6Ai). The extent of recovery of growth rates on cellular adaptation depended on the magnitude of difference between the new average environment and the cellular decision-making point (*p*_2_ − *p*_*I*_), with larger values leading to asymptotically positive growth rates. Thus, we found that the adapted growth rate was larger in the most lethal environment, *p*_2_ = 1 relative to intermediate environments with *p*_*I*_ *< p*_2_ *<* 1, which ultimately lead to a larger population at the end time point (Figure 6Aii). Such adverse effects of continued treatment on population size control as compared to intermittent treatment has been previously observed in melanoma cells (35).

**Fig. 6.**
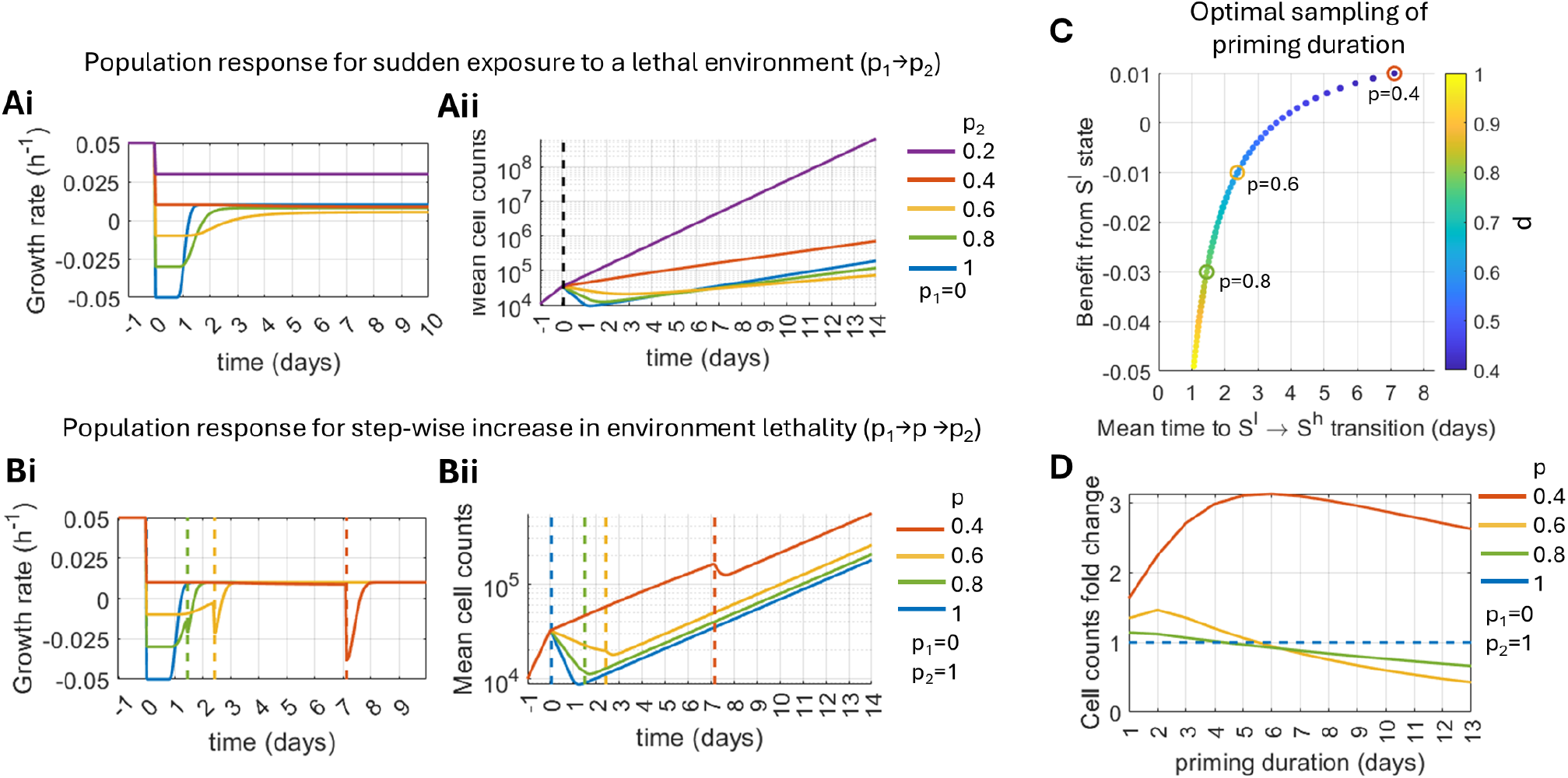
Cellular memory of sublethal environments prime for a quick adaptation to future lethal environment exposures. **A)** Dynamic changes in the average fitness and mean population size to sudden exposure to cytotoxic environments. Here, the stochastic environment has zero autocorrelation, *ρ*_*EE*_ (1) = 0, and the mean environment, *P (E*_*n*_ = *E*^*h*^) = *p*_1_ = 0 for time *<* 0, and *P (E*_*n*_ = *E*^*h*^) = *p*_2_ = {0.2, 0.4, 0.6, 0.8, 1} for time ≥ 0. **B)** Dynamic changes in the fitness and mean population size when the cell population was primed with less lethal environments (0 *< p <* 1) for a defined duration before switching to the most lethal environment (*p*_2_ = 1). The duration of the priming environment, *P (E*_*n*_ = *E*^*h*^) = *p* = {0.4, 0.6, 0.8}, was sampled from the mean time the cell took to switch its state from *S*^*l*^ → *S*^*h*^ for the chosen priming environment as highlighted in panel D. **C)** Optimal sampling of the priming duration with less lethal environment based on the mean adaptation to *S*^*h*^ state. The y-axis represents the average fitness the cell gains from residing in the *S*^*l*^ state until it switches to the *S*^*h*^ state. The red, yellow, and green circles highlight mean adaptation time vs fitness for intermediate environments *p* = {0.4, 0.6, 0.8}, respectively. **D)** Fold change in mean cell counts at the end time point (day 14) for primed populations relative to unprimed populations for increasing priming duration and variable priming environment (*p*). In all the above analyses, 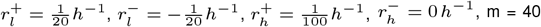, and *c*_*lh*_ = *c*_*hl*_ = 0. The initial population size was 10^4^ cells, and the population grew in an *E*^*l*^ environment (*p*_1_ = 0) for 24h before switching environments at time 0.

Having characterized the cellular and population-level responses to step changes in the environment with increasing levels of toxicity, we next analyzed how priming with less lethal environments (ℙ (*E*_*n*_ = *E*^*h*^) = *p*) affects the response to future lethal exposures. To choose priming durations for less lethal environments, we leveraged our understanding of cellular memory dynamics to quantify the mean time a cell took to attain the *S*^*h*^ state upon switching to intermediate environments, knowing that *S*^*h*^ is the optimal state when the environment eventually becomes highly lethal. Although there was a significant decrease in mean transition time to the *S*^*h*^ state with increasing lethality of the intermediate environment, *p*, we observed that cells exhibited reduced growth rates during the transition period (Figure 6C). On priming the cell population for a duration appropriate to the given intermediate environment lethality, followed by switching to the most lethal environment, we observed an enhanced population size compared to the cell population directly exposed to the highly lethal environment (Figure 6B). The above observation holds across all intermediate environments, with the priming benefit dependent on the intermediate environment and its priming duration (Figure 6D). However, prolonged priming could also result in a negative outcome with overall reduced population size as compared to a direct switch to a highly lethal environment (Figure 6D). Such a favorable window of priming durations depends on cellular memory and adaptation costs, with larger memory size further expanding the favorable window, while larger adaptation costs shrink it (Figure S10).

Overall, the proposed cellular memory and decision-making model recapitulated the observed enhancement in population survival in lethal environments with higher drug concentrations when primed with lower drug concentrations for an appropriate duration (9).

## Discussion

Cells must continually adapt to fluctuations in their environments by making phenotypic choices that enhance growth or survival or result in desirable actions (such as migration toward a nutrient source). In this study we developed a phenomenological approach to relate time-averaged past environmental signals and cellular decisions to switch its state to optimize for cell’s average fitness in fluctuating environments; using this framework, we have elucidated how the decay timescales of past environment signals from the memory and adaptation cost together influence cell state switching and growth in fluctuating environments.

In periodic environments, we observed that increasing memory size resulted in reduced fitness, irrespective of the environment period (Figure 3B). Such a reduction in fitness arose from slower adaptation to fluctuating environments with large memory sizes (Figure 3C). Since increasing memory size reduces the rate of degradation of key biomolecules that determine the cell state, diminished cell state transitions upon environmental switches were observed with reduced mRNA degradation rates in a simple bistable gene regulatory circuit (17). Another way that memory size could influence cellular adaptation is by controlling variability in cell states, with smaller memory sizes leading to larger variability and promoting instantaneous population-level adaptation via bet-hedging (36, 37). Further, studies on *E. coli* and *S. cerevisiae* adapting to nutrient shifts and oscillating osmotic stress, respectively, have reported suboptimal growth at specific intermediate environmental periods because of hyperactivation of the involved response pathways when environmental oscillation timescales matched with the pathway’s activation and deactivation timescales (4, 38). By analyzing changes in fitness in oscillating environments with variable period and cell memory capacity, we were able to recapitulate the above fitness reduction at intermediate frequencies. However, we found that the suboptimal period also depends on the cell’s adaptation cost (Figure S3). Thus, there exists a relationship between the relaxation timescales of the signaling response, the adaptation cost, and the efficiency of cellular adaptation in oscillating environments.

In periodic environments, we also observed non-monotone changes in fitness with increasing adaptation cost, with distinct costs conferring optimal fitness depending on the period of an oscillating environment (Figure 3D). The presence of environment-dependent optimal costs would suggest that subpopulations having extreme adaptation costs – resulting either from, for instance, genetic mutation or from population-level heterogeneity – may be most effectively selected for in an evolutionary experiment in periodic environments. Our model would predict that such ‘fluctuation-advantaged’ subpopulations exhibit improved adaptation and fitness dynamics relative to wild-type only in fluctuating, but not constant, environments (5). In correlated stochastic environments, we observed that large memory sizes or high adaptation costs reduce fitness for increasing intertemporal environmental correlation (Figure 4B). In this setting, fitness reduction occurred because large memory or adaptation costs increase a cell’s inertia to switch its state, which is beneficial in uncertain environments but not otherwise. Such an inertia to cell state changes has been empirically observed with cells growing under fluctuating osmotic stress with distinct environment correlations, wherein cells exhibited minimal changes in their internal state when grown in fluctuating environments having zero autocorrelation. These cells exhibited major transcriptomic and epigenomic changes when grown in fluctuating environments having positive autocorrelation (39).

While studying correlated stochastic environments, we also observed a general trend of increasing mean and decreasing variance of fitness with increasing correlation (Figure 4C). The above trends in the mean and variance of fitness were also empirically observed in *S. cerevisiae* cells exposed to fluctuating osmotic stresses with varying correlations (29). Although the precise cellular response of *S. cerevisiae* to such fluctuating environments is yet to be understood (Figure 4E), our result suggests a potential role of time-averaging environment fluctuations and its influence on cell state transitions to understand the dynamics of *S. cerevisiae* phenotypic response, as has been observed previously for periodic fluctuations. Further, we observed that memory size and adaptation cost modulate cell state transition probabilities, such that optimal fitness may be achieved by either restricting cell state transitions in negatively correlated environments or having cell state closely follow environmental transitions in positively correlated environments (Figure 4D, F). Thus, our modeling approach predicts that to optimally gain from distinct environmental conditions would require either 1) active changes in memory size and adaptation cost during the span of few cell generations, 2) environment-dependent changes in biomolecule degradation rates or 3) cell population heterogeneity, with examples of all three reported in the literature (8, 9, 28, 31, 40–43).

We showed that differences among cells in their adaptation costs to transition to a resistant cell state in our modeling framework can represent experimentally observed distinct subpopulations of melanoma cells with varying adaptation potentials to lethal drug treatment and can capture their population dynamics (Figure 5). A similar existence of distinct subpopulations with varied adaptive potential, associated with their adaptation costs, was observed with *S. cerevisiae* cells (8). In the experiment, *S. cerevisiae* cells were modified to place their *HIS3* gene of the histidine biosynthesis system under the control of the GAL system such that histidine is synthesized in a galactose-rich, but not glucose-rich, environment. Thus, upon switching cells from galactose to glucose in a medium lacking histidine, a major fraction of the population became growth-arrested, with outright metabolic inactivity or low metabolic activity, and only a small fraction (about 4%) showed metabolic recovery and concomitant resumption of growth. Intriguingly, within the metabolically recovering population, there was a distribution of recovery times associated with the amount of energy a cell consumed before resuming growth in the stressed environment. Moreover, the cells that showed early metabolic recovery and adaptation ultimately comprised a higher fraction of the population than the remaining recovering cells. Although the developed model and the example presented above highlight the role of one-time costs that cells bear while adapting to new environments, there is also a continued influx of energy required to maintain a phenotypic state due to the presence of negative feedback loops and active turnover of biomolecules (23, 24). This cost of maintaining a phenotype can shape the phenotypic structure of a population and influence the population’s overall response to environmental shifts (18, 26, 27, 40, 44). Our future efforts will focus on understanding the impact of phenotype-maintenance costs on cellular adaptation by updating the current framework to account for a cost-dependent reduction in fitness during environment-phenotype matches.

Lastly, we observed that priming cells with sub-lethal environments enhances their survival in subsequent lethal environments (Figure 6). Although we motivated the above analysis based on empirical observations of melanoma cells, there is an increasing number of studies demonstrating enhanced survival when priming cells with sublethal doses across organisms – bacteria, yeast, and cancer – and different stressors – cytotoxic drug, salinity, pH, temperature, and oxidative stresses (9, 33, 34). Furthermore, preexposure to one stressor has also been shown to enhance the adaptive potential to another stressor, a phenomenon referred to as ‘cross-protection’ (34, 45, 46). We note that cellular machinery that encodes a memory of past environment exposure is likely to be organism dependent and more complex than what has been considered in this study, our analysis highlights how the residual signal in cellular memory that follows probabilistic erasure (according to first-order decay) is sufficient to explain observed biological phenomena (9, 47–49). Taking into account the role of cellular memory in adaptation can improve our dynamical understanding of the evolving state of resistance in the population to different drug exposures, which can ultimately inform the design of improved therapeutic strategies (Figure 6C, D).

Overall, the developed framework and applications highlight the influence of cellular memory timescales and adaptation costs on cell-state switching and fitness in distinct fluctuating environments. We show that, although fast turnover of cellular memory with small memory sizes, along with low adaptation cost, enable faster cell state switching and improve adaptation, their fitness advantage depends on the characteristics of fluctuating environments. While we have substantiated the observed functional roles of cellular memory and adaptation costs in cellular adaptation by comparing them closely with recent experimental observations, this work has a few key limitations. First, the dynamics of cellular memory, which drive adaptation, are based solely on the encoding of new signals and the decay of old ones. However, more complex intracellular signaling dynamics, which can perform intricate signal processing such as temporal gradient and fold-change detection, have been observed to drive cellular adaptation in fluctuating environments (50–52). Subsequent studies that consider the precise intracellular signaling dynamics to study cellular adaptation in specific contexts will be an important follow-up task (53). Further, this work considered a discrete-time description of cellular adaptation. Future modeling efforts in a continuous-time setting will offer an additional conceptual tool for the application (53, 54). Lastly, we assumed ‘discrete’ cell states, with transitions between them determined solely by past environmental signals. However, cell states are far from discrete due to stochasticity in intracellular biochemical reactions (55, 56). Such subtle differences in discernible cell states may contribute to the survival of a fraction of cells in the event of environmental change (31, 37, 43, 57). Although population-level variability in adaptation costs, as discussed above, can capture this cell-cell variability, an explicit account of stochastic state switching along with active sensing in a single model may in the future be more informative for their combined effects on cellular adaptation and long-term growth benefits in fluctuating environments (58, 59). Nonetheless, our phenomenological framework offers key insights into the effects of cell molecular memory and adaptation costs on cellular adaptations in fluctuating environments. These results contribute to our understanding of how intracellular processes shape cellular responses to changing environments.

## AUTHOR CONTRIBUTIONS

P.J., M.K.J, and J.T.G. conceived of and designed the research. P.J. performed the research. P.J., M.K.J., and J.T.G contributed new analytic equations, analyzed the results, and wrote the paper. J.T.G. and M.K.J. supervised the research.

## COMPETING FINANCIAL INTERESTS

The authors declare no competing interests.

### ACKNOWLEDGEMENTS

M.K.J. and P.J. were supported by Param Hansa Philanthropies. J.T.G. was supported by the Cancer Prevention Research Institute of Texas (RR210080) and the National Institute of General Medical Sciences of the NIH (R35GM155458). J.T.G. is a CPRIT Scholar in Cancer Research.

## Data and Code Availability

The code files for numerical evaluations of the formulations developed in this study can be found at: https://github.com/Paras-Jain20/Memory-and-cost-in-cellular-adaptation

## Notes

### Competing Interest Statement

The authors have declared no competing interest.

### Summary of Updates

Fixed typo in Figure 4 and SI Figures S6 and S7.

## Bibliography

1. Jacques Monod. The growth of bacterial cultures. Selected Papers in Molecular Biology by Jacques Monod, 139:606, 2012.

2. David W Erickson, Severin J Schink, Vadim Patsalo, James R Williamson, Ulrich Gerland, and Terence Hwa. A global resource allocation strategy governs growth transition kinetics of escherichia coli. Nature, 551(7678):119–123, 2017.

3. Sreenath V Sharma, Diana Y Lee, Bihua Li, Margaret P Quinlan, Fumiyuki Takahashi, Shyamala Maheswaran, Ultan McDermott, Nancy Azizian, Lee Zou, Michael A Fischbach, et al. A chromatin-mediated reversible drug-tolerant state in cancer cell subpopulations. Cell, 141 (1):69–80, 2010.

4. Jen Nguyen, Vicente Fernandez, Sammy Pontrelli, Uwe Sauer, Martin Ackermann, and Roman Stocker. A distinct growth physiology enhances bacterial growth under rapid nutrient fluctuations. Nature Communications, 12(1):3662, 2021.

5. Clare I Abreu, Shaili Mathur, and Dmitri A Petrov. Environmental memory alters the fitness effects of adaptive mutations in fluctuating environments. Nature Ecology & Evolution, 8(9): 1760–1775, 2024.

6. Roman Stocker. Marine microbes see a sea of gradients. science, 338(6107):628–633, 2012.

7. Tom Fenchel. Microbial behavior in a heterogeneous world. Science, 296(5570):1068–1071, 2002.

8. Gabrielle Woronoff, Philippe Nghe, Jean Baudry, Laurent Boitard, Erez Braun, Andrew D Griffiths, and Jérôme Bibette. Metabolic cost of rapid adaptation of single yeast cells. Proceedings of the National Academy of Sciences, 117(20):10660–10666, 2020.

9. Jingxin Li, Pavithran T Ravindran, Aoife O’Farrell, Gianna T Busch, Ryan H Boe, Zijian Niu, Sean Woo, Margaret C Dunagin, Naveen Jain, Yogesh Goyal, et al. Ap-1 mediates cellular adaptation and memory formation during therapy resistance. bioRxiv, 2024.

10. Jason T George. Optimal phenotypic adaptation in fluctuating environments. Biophysical Journal, 122(22):4414–4424, 2023.

11. Nico Geisel, Jose MG Vilar, and J Miguel Rubi. Optimal resting-growth strategies of microbial populations in fluctuating environments. PLoS One, 6(4):e18622, 2011.

12. Edo Kussell, Roy Kishony, Nathalie Q Balaban, and Stanislas Leibler. Bacterial persistence: a model of survival in changing environments. Genetics, 169(4):1807–1814, 2005.

13. Edo Kussell and Stanislas Leibler. Phenotypic diversity, population growth, and information in fluctuating environments. Science, 309(5743):2075–2078, 2005.

14. Gerardo Aquino, Luke Tweedy, Doris Heinrich, and Robert G Endres. Memory improves precision of cell sensing in fluctuating environments. Scientific reports, 4(1):5688, 2014.

15. Gary Friedman, Stephen McCarthy, and Dmitrii Rachinskii. Hysteresis can grant fitness in stochastically varying environment. PLoS One, 9(7):e103241, 2014.

16. Stefan Legewie, Dennis Dienst, Annegret Wilde, Hanspeter Herzel, and Ilka M Axmann. Small rnas establish delays and temporal thresholds in gene expression. Biophysical journal, 95(7):3232–3238, 2008.

17. Elizabeth R Westbrook, Hugh Z Ford, Vlatka Antolović, and Jonathan R Chubb. Clearing the slate: Rna turnover to enable cell-state switching? Development, 150(19), 2023.

18. Eitan Rotem, Adiel Loinger, Irine Ronin, Irit Levin-Reisman, Chana Gabay, Noam Shoresh, Ofer Biham, and Nathalie Q Balaban. Regulation of phenotypic variability by a threshold-based mechanism underlies bacterial persistence. Proceedings of the National Academy of Sciences, 107(28):12541–12546, 2010.

19. Guillaume Lambert and Edo Kussell. Memory and fitness optimization of bacteria under fluctuating environments. PLoS genetics, 10(9):e1004556, 2014.

20. Angela Oliveira Pisco, Amy Brock, Joseph Zhou, Andreas Moor, Mitra Mojtahedi, Dean Jackson, and Sui Huang. Non-darwinian dynamics in therapy-induced cancer drug resistance. Nature communications, 4(1):2467, 2013.

21. Stefan Hohmann. Osmotic stress signaling and osmoadaptation in yeasts. Microbiology and molecular biology reviews, 66(2):300–372, 2002.

22. Maor Knafo, Shahar Rezenman, Reinat Nevo, Ivgeni Tsigalnitski, Ziv Reich, and Ruti Kapon. Variability in protein expression and fitness under stress: Distinct modes of early adaptation of populations. PRX Life, 3(1):013006, 2025.

23. Ganhui Lan, Pablo Sartori, Silke Neumann, Victor Sourjik, and Yuhai Tu. The energy– speed–accuracy trade-off in sensory adaptation. Nature physics, 8(5):422–428, 2012.

24. Wenzhe Ma, Ala Trusina, Hana El-Samad, Wendell A Lim, and Chao Tang. Defining network topologies that can achieve biochemical adaptation. Cell, 138(4):760–773, 2009.

25. Pankaj Mehta and David J Schwab. Energetic costs of cellular computation. Proceedings of the National Academy of Sciences, 109(44):17978–17982, 2012.

26. Santiago Sandoval Motta, Philippe Cluzel, and Maximino Aldana. Adaptive resistance in bacteria requires epigenetic inheritance, genetic noise, and cost of efflux pumps. PloS one, 10(3):e0118464, 2015.

27. Daniel M Stoebel, Antony M Dean, and Daniel E Dykhuizen. The cost of expression of escherichia coli lac operon proteins is in the process, not in the products. Genetics, 178(3): 1653–1660, 2008.

28. Matthew R Bennett, Wyming Lee Pang, Natalie A Ostroff, Bridget L Baumgartner, Sujata Nayak, Lev S Tsimring, and Jeff Hasty. Metabolic gene regulation in a dynamically changing environment. Nature, 454(7208):1119–1122, 2008.

29. Pascal Hersen, Megan N McClean, Lakshminarayanan Mahadevan, and Sharad Ramanathan. Signal processing by the hog map kinase pathway. Proceedings of the National Academy of Sciences, 105(20):7165–7170, 2008.

30. Vimalathithan Devaraj and Biplab Bose. Morphological state transition dynamics in egf-induced epithelial to mesenchymal transition. Journal of clinical medicine, 8(7):911, 2019.

31. Sydney M Shaffer, Margaret C Dunagin, Stefan R Torborg, Eduardo A Torre, Benjamin Emert, Clemens Krepler, Marilda Beqiri, Katrin Sproesser, Patricia A Brafford, Min Xiao, et al. Rare cell variability and drug-induced reprogramming as a mode of cancer drug resistance. Nature, 546(7658):431–435, 2017.

32. Gerardo Rubino and Bruno Sericola. Markov chains and dependability theory. Cambridge University Press, 2014.

33. Alexandro Rodríguez-Rojas, Desiree Y Baeder, Paul Johnston, Roland R Regoes, and Jens Rolff. Bacteria primed by antimicrobial peptides develop tolerance and persist. PLoS Pathogens, 17(3):e1009443, 2021.

34. Diana R Andrade-Linares, Anika Lehmann, and Matthias C Rillig. Microbial stress priming: a meta-analysis. Environmental Microbiology, 18(4):1277–1288, 2016.

35. Andrew J Kavran, Scott A Stuart, Kristyn R Hayashi, Joel M Basken, Barbara J Brandhuber, and Natalie G Ahn. Intermittent treatment of brafv600e melanoma cells delays resistance by adaptive resensitization to drug rechallenge. Proceedings of the National Academy of Sciences, 119(12):e2113535119, 2022.

36. Juan M Pedraza and Johan Paulsson. Effects of molecular memory and bursting on fluctuations in gene expression. Science, 319(5861):339–343, 2008.

37. Sabrina L Spencer, Suzanne Gaudet, John G Albeck, John M Burke, and Peter K Sorger. Non-genetic origins of cell-to-cell variability in trail-induced apoptosis. Nature, 459(7245): 428–432, 2009.

38. Amir Mitchell, Ping Wei, and Wendell A Lim. Oscillatory stress stimulation uncovers an achilles’ heel of the yeast mapk signaling network. Science, 350(6266):1379–1383, 2015.

39. Christelle Leung, Daphné Grulois, Leandro Quadrana, and Luis-Miguel Chevin. Phenotypic plasticity evolves at multiple biological levels in response to environmental predictability in a long-term experiment with a halotolerant microalga. PLoS Biology, 21(3):e3001895, 2023.

40. Jue Wang, Esha Atolia, Bo Hua, Yonatan Savir, Renan Escalante-Chong, and Michael Springer. Natural variation in preparation for nutrient depletion reveals a cost–benefit trade-off. PLoS biology, 13(1):e1002041, 2015.

41. Lydia Freddolino, Jamie Yang, Amir Momen-Roknabadi, and Saeed Tavazoie. Stochastic tuning of gene expression enables cellular adaptation in the absence of pre-existing regulatory circuitry. Elife, 7:e31867, 2018.

42. Mariona Nadal-Ribelles, Guillaume Lieb, Carme Solé, Yaima Matas, Ugo Szachnowski, Sara Andjus, Maria Quintana, Mònica Romo, Aitor Gonzalez Herrero, Antonin Morillon, et al. Transcriptional heterogeneity shapes stress-adaptive responses in yeast. Nature communications, 16(1):2631, 2025.

43. Yogesh Goyal, Gianna T Busch, Maalavika Pillai, Jingxin Li, Ryan H Boe, Emanuelle I Grody, Manoj Chelvanambi, Ian P Dardani, Benjamin Emert, Nicholas Bodkin, et al. Diverse clonal fates emerge upon drug treatment of homogeneous cancer cells. Nature, 620(7974):651–659, 2023.

44. Lucas Henrion, Juan Andres Martinez, Vincent Vandenbroucke, Mathéo Delvenne, Samuel Telek, Andrew Zicler, Alexander Grünberger, and Frank Delvigne. Fitness cost associated with cell phenotypic switching drives population diversification dynamics and controllability. Nature Communications, 14(1):6128, 2023.

45. Amir Mitchell, Gal H Romano, Bella Groisman, Avihu Yona, Erez Dekel, Martin Kupiec, Orna Dahan, and Yitzhak Pilpel. Adaptive prediction of environmental changes by microorganisms. Nature, 460(7252):220–224, 2009.

46. Qiaoning Guan, Suraiya Haroon, Diego González Bravo, Jessica L Will, and Audrey P Gasch. Cellular memory of acquired stress resistance in saccharomyces cerevisiae. Genetics, 192(2):495–505, 2012.

47. Sharmistha Kundu and Craig L Peterson. Dominant role for signal transduction in the transcriptional memory of yeast gal genes. Molecular and cellular biology, 30(10):2330–2340, 2010.

48. Eugenia Lyashenko, Mario Niepel, Purushottam D Dixit, Sang Kyun Lim, Peter K Sorger, and Dennis Vitkup. Receptor-based mechanism of relative sensing and cell memory in mammalian signaling networks. Elife, 9:e50342, 2020.

49. Akhilesh Nandan, Abhishek Das, Robert Lott, and Aneta Koseska. Cells use molecular working memory to navigate in changing chemoattractant fields. Elife, 11:e76825, 2022.

50. Pulin Li, Joseph S Markson, Sheng Wang, Siheng Chen, Vipul Vachharajani, and Michael B Elowitz. Morphogen gradient reconstitution reveals hedgehog pathway design principles. Science, 360(6388):543–548, 2018.

51. Benoit Sorre, Aryeh Warmflash, Ali H Brivanlou, and Eric D Siggia. Encoding of temporal signals by the tgf-*β* pathway and implications for embryonic patterning. Developmental cell, 30(3):334–342, 2014.

52. Yaron E Antebi, Nagarajan Nandagopal, and Michael B Elowitz. An operational view of intercellular signaling pathways. Current opinion in systems biology, 1:16–24, 2017.

53. Josiah Kratz, Huijing Wang, Fangwei Si, and Shiladitya Banerjee. Power-law memory governs bacterial adaptation and learning in fluctuating environments. bioRxiv, pages 2025–04, 2025.

54. Alexander P Browning, Rebecca M Crossley, Chiara Villa, Philip K Maini, Adrianne L Jenner, Tyler Cassidy, and Sara Hamis. Identifiability of phenotypic adaptation from low-cell-count experiments and a stochastic model. PLoS Computational Biology, 21(6):e1013202, 2025.

55. Arjun Raj, Charles S Peskin, Daniel Tranchina, Diana Y Vargas, and Sanjay Tyagi. Stochastic mrna synthesis in mammalian cells. PLoS biology, 4(10):e309, 2006.

56. Arjun Raj and Alexander Van Oudenaarden. Nature, nurture, or chance: stochastic gene expression and its consequences. Cell, 135(2):216–226, 2008.

57. S Hamis et al. Phenotypic heterogeneity in temporally fluctuating environments. Physical Biology, 22(5), 2025.

58. Dino Osmanović, Yitzhak Rabin, and Yoav Soen. A model of epigenetic inheritance accounts for unexpected adaptation to unforeseen challenges. Advanced Science, page 2414297, 2025.

59. Sara Hamis, Alexander P Browning, Adrianne L Jenner, Chiara Villa, Philip K Maini, and Tyler Cassidy. Growth rate-driven modelling suggests that phenotypic adaptation drives drug resistance in brafv600e-mutant melanoma. Communications Biology, 2026.

