## Supplementary material for "Effects of Cellular Memory and Adaptation Cost on Optimal Survival in Fluctuating Environments": SI Text and Figures

April 22, 2026

#### S1 Introduction

Adaptation in response to changing environmental conditions is a hallmark cellular behavior. Cellular adaptation to changing environments is seldom spontaneous, involves a lag phase, and can be informed by previous environmental exposures. For example, bacterial cells experiencing environmental shifts in glucose and lactose concentrations exhibit a decreasing lag in their growth phase with shorter time period of fluctuations, suggestive of the presence of a 'cellular memory' of past glucose/lactose environments [2]. Such cellular memory can be orchestrated by variable levels of mRNA and protein or by chromatin modifications [3, 4]. Thus, the erasure of the past memory (degradation of mRNA and protein or removal of chromatin-modifications) is a necessary consequence of cellular adaptation to new external conditions. Apart from the erasure of the memory, cellular adaptation also depends on the *de-novo* production of mRNA and protein or chromatin remodeling, which are necessary for cells to grow (or persist) in a new environment. Such actions occur at the expense of diverting energy to encode molecular memory that would otherwise be available for division. This *de-novo* production can thus be viewed as an effect cost resulting from adaptation or cell state transition.

Here, we study how cell memory dynamics, along with a non-negligible cost of adaptation, together influence cellular decision-making to maximize the average fitness in a fluctuating environment.

#### S2 Dynamics of fluctuating environment

**Stochastic environments**
Here we consider stochastic environment fluctuations with $\mathbb{P}\left(E_n = E^h\right) = p$ and one step autocorrelation of $\rho_{XX}(1)$. The dynamics of the environment can be captured using the following transition probability matrix:

$$\mathbf{P_E} = \begin{matrix} (\boldsymbol{E}_n \backslash \boldsymbol{E}_{n+1}) \\ E^l \\ E^h \end{matrix} \begin{matrix} \begin{matrix} E^l & \quad E^h \end{matrix} \\ \begin{pmatrix} 1-p_1 & p_1 \\ p_2 & 1-p_2 \end{pmatrix} \end{matrix}$$

where $p_1$ and $p_2$ are the transition probabilities between environmental states with $p_1 = \min(1, p\,(1-\rho_{XX}(1)))$ and $p_2 = \min(1, (1-p)\,(1-\rho_{XX}(1)\,)\,)$. When $\rho_{XX}(1) = 0$, we have $p_1 = p$ and $p_2 = 1 - p$.

**Periodic environments**
In contrast with a stochastic environment, the probability $p_n$ for an environment with period $T$ at a time instance $n = r\,T + l$, where $r$ and $l$ being integers, is given by:

$$\mathbb{P}\left(E_n = E^h\right) = p_n = \begin{cases} 0, & 1 \le l \le T_{E^l\,bias} \\ 1, & T_{E^l\,bias} < l \le T, \end{cases}$$

where, $T_{E^l\,bias} = \lfloor E^l_{bias} T \rfloor$ with $E^l_{bias}$ denoting the fraction of environmental period in $E^l$ state and floor function $\lfloor\cdot\rfloor$ giving greatest integer less than or equal to its argument.

**Deterministic environments**
Deterministic environments are a special case of stochastic environments with $p = 0$ or $p = 1$.

### S3 Dynamics of cellular memory

In the cellular memory model, all environmental states experienced by the cell over a period of $m$ time steps have equal probability to be forgotten (for example, selecting and replacing colored balls in the Pólya urn model). At each time step a random past experience is sampled from the cellular memory and replaced with a new environmental encounter.

Letting $K_n$ represent the number of $E^h$ environmental states retained in the memory window of the cell at time $n$, we note that the fractions $\frac{K_n}{m}$ and $\left(1 - \frac{K_n}{m}\right)$ denote the probabilities of losing a $E^h$ or $E^l$ environmental experience from the memory window, respectively.

#### S3.1 Cell memory dynamics in stochastic environments

Since, we consider stochastic environment having $\mathbb{E}\left[E_n\right] = p$ and one step autocorrelation of $p_{EE}(1)$, the dynamics of environment can be captured using the following transition probability matrix:

$$E_{n+1} = \begin{cases} & \text{next state} \qquad\qquad \text{probability} \\ E^h, & \mathbb{1}_{E_n=E^h}\,(1-p_2) + (1-\mathbb{1}_{E_n=E^h})\,p_1 \\ E^l, & \mathbb{1}_{E_n=E^h}\,p_2 + (1-\mathbb{1}_{E_n=E^h})\,(1-p_1) \end{cases} \tag{S1}$$

where, $p_1 = \min(1, p\,(1 - p_{EE}(1)))$ and $p_2 = \min(1, (1-p)\,(1 - p_{EE}(1)\,)\,)$. The above makes use of the usual indicator random variables:

$$\mathbb{1}_{X_n=X^h} = \begin{cases} 1, & X_n = X^h \\ 0, & X_n = X^l \end{cases} \tag{S2}$$

The joint dynamics of environment and past environment encounters is thus given by:

$$\left[K_{n+1}, E_{n+1}\right] = \begin{cases} \text{next state} & \text{probability} \\ \left[K_n, E^h\right], & \mathbb{1}_{E_n=E^h}(1-p_2)\left(\frac{K_n}{m}\right) + (1-\mathbb{1}_{E_n=E^h})p_1\left(\frac{K_n}{m}\right) \\ \left[K_n+1, E^h\right], & \mathbb{1}_{E_n=E^h}(1-p_2)\left(1-\frac{K_n}{m}\right) + (1-\mathbb{1}_{E_n=E^h})p_1\left(1-\frac{K_n}{m}\right) \\ \left[K_n-1, E^h\right], & 0 \\ \left[K_n, E^l\right], & \mathbb{1}_{E_n=E^h}p_2\left(1-\frac{K_n}{m}\right) + (1-\mathbb{1}_{E_n=E^h})(1-p_1)\left(1-\frac{K_n}{m}\right) \\ \left[K_n+1, E^l\right], & 0 \\ \left[K_n-1, E^l\right], & \mathbb{1}_{E_n=E^h}p_2\left(\frac{K_n}{m}\right) + (1-\mathbb{1}_{E_n=E^h})(1-p_1)\left(\frac{K_n}{m}\right) \end{cases} \tag{S3}$$

The above transition probabilities can be represented in the matrix form for an arbitrary memory size (here, $m = 2$) as follows:

$$\mathbf{P}_{\{m=2\}} = \begin{array}{l} ([K_n,E_n]\backslash[K_{n+1},E_{n+1}]) \\ {[0,E^l]} \\ {[0,E^h]} \\ {[1,E^l]} \\ {[1,E^h]} \\ {[2,E^l]} \\ {[2,E^h]} \end{array} \begin{array}{c} \begin{array}{cccccc} [0,E^l] & [0,E^h] & [1,E^l] & [1,E^h] & [2,E^l] & [2,E^h] \end{array} \\ \begin{pmatrix} (1-p_1) & 0 & 0 & p_1 & 0 & 0 \\ p_2 & 0 & 0 & (1-p_2) & 0 & 0 \\ (1-p_1)\frac{1}{m} & 0 & (1-p_1)(1-\frac{1}{m}) & p_1(\frac{1}{m}) & 0 & p_1(1-\frac{1}{m}) \\ p_2(\frac{1}{m}) & 0 & p_2(1-\frac{1}{m}) & (1-p_2)(\frac{1}{m}) & 0 & (1-p_2)(1-\frac{1}{m}) \\ 0 & 0 & (1-p_1) & 0 & 0 & p_1 \\ 0 & 0 & p_2 & 0 & 0 & 1-p_2 \end{pmatrix} \end{array}$$

We evaluate the stationary distribution of $[K_n, E_n]$ pair for a fluctuating environment driven by $p$ and $\rho_{EE}(1)$ as follows:

$$\boldsymbol{y} = \lim_{n\to\infty} \boldsymbol{y}_0 P^n \tag{S4}$$

where, $\boldsymbol{y} = \{y_{0,l}\,, y_{0,h}\,, y_{1,l}\,, y_{1,h}\,, ..., y_{k,e}\,, ..., y_{m,l}\,, y_{m,h}\}$ denote the joint stationary probabilities of the cell memory states and environment with $k \in \{0, 1, 2, ..., m\}$ and $e \in \{l, h\}$ for $E^l$ and $E^h$ environments, and $\boldsymbol{y}_0$ denote an arbitrary distribution of initial joint probabilities.

The stationary probability of cell memory state $K_n$ is then quantified as:

$$x_k = y_{k,l} + y_{k,h} \tag{S5}$$

where, $k \in 0, 1, 2, ..., m$

###### S3.1.1 Stationary distribution of $K_n$ in uncorrelated stochastic environment

In uncorrelated stochastic environments, the state transition probabilities between cell memory states reduces to:

$$K_{n+1} = \begin{cases} & \text{next state} \qquad \text{probability} \\ K_n, & \left(\frac{K_n}{m}\right)p + \left(1-\frac{K_n}{m}\right)\left(1-p\right) \\ K_n+1, & \left(1-\frac{K_n}{m}\right)p \\ K_n-1, & \left(\frac{K_n}{m}\right)\left(1-p\right) \end{cases} \tag{S6}$$

Using the above state transition probabilities, we next derive the stationary probability distribution $\boldsymbol{x}$ of $K_n$ as follows:
At stationarity,

$$x_k = \sum_{j=0}^{m} x_j p_{jk}, \tag{S7}$$

where, $p_{jk} = \mathbb{P}(K_{n+1} = k\,|K_n = j)$ and $p_{jk} \neq 0$ only when $|j - k| \leq 1$. We expand the eq. (S7) for each $K_n = k \in \{0, 1, ..., m-1, m\}$ while substituting the values of $p_{jk}$ from eq. (S3) in each term. This leads to $m+1$ equations in $m+1$ unknowns ($[x_0, x_1, ..., x_N]$) whose solution can be written in terms of $x_0$ as follows:

$$x_k = x_0 \binom{m}{k} \left(\frac{p}{1-p}\right)^k. \tag{S8}$$

Since

$$\sum_{k=0}^{m} x_k = 1,$$

we obtain

$$x_0 = \frac{1}{\sum_{j=0}^{m} \binom{m}{j} \left(\frac{p}{1-p}\right)^j} = (1 + \frac{p}{1-p})^{-m} = (1-p)^m,$$

and thus, from eq. (S8) we have,

$$x_k = \binom{m}{k} p^k (1-p)^{m-k}. \tag{S9}$$

Thus, $x \sim Binomial(m, p)$.

#### S3.2 Cell memory dynamics in periodic environment

For a cell experiencing a periodically varying environment, the distribution of $K_n$ becomes periodic with period $T$ as shown below:

Let $\boldsymbol{x}(n)$ denote the time-dependent row vector of probabilities that the cell has $k \in \{0, 1, 2, ..., m\}$ $S^h$ environmental experiences at time $n$. Now, due to the periodicity of the environment, asymptotically (for $n \to \infty$) we have $\boldsymbol{x}(n) = \boldsymbol{x}(n+T)$. The time evolution of $\boldsymbol{x}(n)$ can then be written as follows:

$$\boldsymbol{x}(rT + l) = \begin{cases} \boldsymbol{x}(rT)\ M_0^l & 1 \leq l \leq T_{E^l_{bias}} \\ \boldsymbol{x}(rT)\ M_0^{T_{E^l_{bias}}}\ M_1^{l - T_{E^l_{bias}}} & T_{E^l_{bias}} < l \leq T \end{cases} \tag{S10}$$

where $r$ is an arbitrary large integer, $M_0$ and $M_1$ are the State Transition Matrices (STMs) when every environmental encounter is $S^l$ and $S^h$, respectively. Elements of the $M_0$ and $M_1$ matrices are obtained as follows: For $M_0$, we substitute $p_1 = 0$, $p_2 = 1$, and $\mathbb{1}_{E_n = E^h} = 0$ in eq. (S6), obtaining

$$K_{n+1} = \begin{cases} \text{next state} & \text{probability} \\ K_n, & \left(1 - \dfrac{K_n}{m}\right) \\ K_n + 1, & 0 \\ K_n - 1, & \left(\dfrac{K_n}{m}\right) \end{cases} \tag{S11}$$

Similarly, for $M_1$ we substitute $p_1 = 1$, $p_2 = 0$ and $\mathbb{1}_{E_n = E^h} = 1$ in eq. (S3), which gives

$$K_{n+1} = \begin{cases} \text{next state} & \text{probability} \\ K_n, & \left(\dfrac{K_n}{m}\right) \\ K_n + 1, & \left(1 - \dfrac{K_n}{m}\right) \\ K_n - 1, & 0 \end{cases} \tag{S12}$$

To obtain $\boldsymbol{x}(rT)$ in eq. (S10)], put $l = T$ and denote the transpose of matrix (or vector) $A$ by $A'$. Then,

$$\boldsymbol{x}(rT+T) = \boldsymbol{x}(rT)\, M_0^{T_{E^l_{bias}}}\, M_1^{T_{E^h_{bias}}} = \boldsymbol{x}(rT)$$
$$\boldsymbol{x}(rT)\big[I - M_0^{T_{E^l_{bias}}}\, M_1^{T_{E^h_{bias}}}\big] = 0$$
$$\big[I - M_0^{T_{E^l_{bias}}}\, M_1^{T_{E^h_{bias}}}\big]'\, \boldsymbol{x}(rT)' = 0.$$

where, $T_{E^h_{bias}} = T - T_{E^l_{bias}}$. Thus, the $\boldsymbol{x}(rT)'$ corresponds to the null solutions of matrix $\big[I - M_0^{T_{E^l_{bias}}} M_1^{T_{E^h_{bias}}}\big]'$.

### S4 Identifying the optimal cell state given the mean environment

With the cell experiencing either $E^h$ or $E^l$ environment at any time instance '$n$', we consider that there exist two phenotypic states, $S^h$ and $S^l$, which are preferred in environments $E^h$ and $E^l$, respectively, and impart greater proliferation tendency to the cell. Here, we denote the proliferation tendency or 'cell fitness' by $r^{\pm}_{l,h}$, with subscripts denoting cell state $S$ and superscripts denoting whether state $S$ matches $(+)$ or mismatches $(-)$ the environment $E$. For example, the cell's proper match of $S^l$ to environment $E^l$ results in a fitness value $r^+_l$, and a similar fitness value $r^+_h$ occurs for the cell's match of $S^h$ to environment $E^h$. However, state-environmental mismatch (either $S^l$ with $E^h$ or $S^h$ with $E^l$) results in a reduced fitness ($r^-_l$ or $r^-_h$) of the cell. The state-environmental (mis)matching, as described above, by definition, requires inequalities:

$$r^-_l \le r^+_l, \quad \text{and} \quad r^-_h \le r^+_h. \tag{S13}$$

In this model, cells in the population switch between $S^h$ and $S^l$ phenotypic states by estimating the preferable stated based on their memory of previous environmental occurrences so as to maximize their average fitness in the upcoming environmental encounter. Further, as cell state switching requires resource diversion to orchestrate changes in multiple intracellular processes, we also incorporate a cost associated with each cell state switching event - $c_{lh}$ (cost for $S^l$-to-$S^h$ switch) and $c_{hl}$ (cost for $S^h$-to-$S^l$ switch).

Letting $E_n$ and $S_n$ to denote the state of the environment and cell at time $n$, the fitness $R_n$ accrued by the cell at time $n$ is given by:

$$R_n = \begin{cases} r^+_h\, \mathbb{1}_{E_n=E^h} + r^-_h(1-\mathbb{1}_{E_n=E^h}) - c_{lh}(1-\mathbb{1}_{S_{n-1}=S^h}), & S_n = S^h; \\ r^+_l(1-\mathbb{1}_{E_n=E^h}) + r^-_l\mathbb{1}_{E_n=E^h}(1-\mathbb{1}_{S_n=S^h}) - c_{hl}\mathbb{1}_{S_{n-1}=S^h}, & S_n = S^l. \end{cases} \tag{S14}$$

This may be represented more succinctly as:

$$\begin{aligned} R_n = {} & \mathbb{1}_{S_n=S^h}\left(r^+_h\,\mathbb{1}_{E_n=E^h} + r^-_h(1-\mathbb{1}_{E_n=E^h})\right) - c_{lh}\mathbb{1}_{S_n=S^h}(1-\mathbb{1}_{S_{n-1}=S^h}) \\ & + \left(r^+_l(1-\mathbb{1}_{E_n=E^h}) + r^-_l\mathbb{1}_{E_n=E^h}\right)(1-\mathbb{1}_{S_n=S^h}) - c_{hl}(1-\mathbb{1}_{S_n=S^h})\,\mathbb{1}_{S_{n-1}=S^h}. \end{aligned} \tag{S15}$$

And clearly we have that $\mathbb{E}\left[\mathbb{1}_{X_n=X^h}\right] = \mathbb{P}\left(X_n = X^h\right)$

For the given mean environment $\mathbb{E}\left[\mathbf{1}_{E_n=E^h}\right] = p$, the average fitness under no-cost adaptation ($c_{lh} = c_{hl} = 0$) while residing in cell state $S_n \in \{S^h, S^l\}$ is given by

$$\begin{aligned} \mathbb{E}\left[R_n \mid S_n = S^h\right] &= r^+_h\, p + r^-_h(1-p) \\ \mathbb{E}\left[R_n \mid S_n = S^l\right] &= r^+_l(1-p) + r^-_l\, p \end{aligned} \tag{S16}$$

The above expected fitness are equal whenever:

$$\mathbb{E}\left[R_n \mid S^h\right] = \mathbb{E}\left[R_n \mid S^l\right]$$

or, equivalently,

$$r_h^+ \; p + r_h^-(1-p) = r_l^+(1-p) + r_l^- \; p,$$

which identifies a unique indifference environment $p_I$ for which each cell state is equally beneficial:

$$p_I = \frac{r_l^+ - r_h^-}{(r_h^+ - r_h^-) + (r_l^+ + r_l^-)}. \tag{S17}$$

When adaptation costs are non-negligible ($c_{lh}$ or $c_{hl} \neq 0$), the above calculations must also account for the cell state at previous time $n-1$:

$$\begin{aligned}
\mathbb{E}\left[R_n \mid S_n = S^h, S_{n-1} = S^h\ \right] &= r_h^+ p + r_h^-(1-p) \\
\mathbb{E}\left[R_n \mid S_n = S^h, S_{n-1} = S^l\ \right] &= r_h^+ p + r_h^-(1-p) - c_{lh} \\
\\
\mathbb{E}\left[R_n \mid S_n = S^l, S_{n-1} = S^l\ \right] &= r_l^+(1-p) + r_l^- p \\
\mathbb{E}\left[R_n \mid S_n = S^l, S_{n-1} = S^h\ \right] &= r_l^+(1-p) + r_l^- p - c_{hl}
\end{aligned} \tag{S18}$$

The additional cost incurred during cell state transitions subsequently affects the indifference environment $p_I$ in eq. (S17) depending on the type of transition as follows:

For navigating a $S^l$-to-$S^h$ transition, we have

$$\mathbb{E}\left[R_n \mid S_n = S^h, S_{n-1} = S^l\ \right] = \mathbb{E}\left[R_n \mid S_n = S^l, S_{n-1} = S^l\ \right],$$

giving

$$p_{lh} = \frac{r_l^+ - r_h^- + c_{lh}}{(r_h^+ - r_h^-) + (r_l^+ + r_l^-)}. \tag{S19}$$

here, $p_{lh}$ represent the indifference environment for $S^l \to S^h$ state transition.

Similarly, for navigating a $S^h$-to-$S^l$ transition, we have

$$\mathbb{E}\left[R_n \mid S_n = S^l, S_{n-1} = S^h\ \right] = \mathbb{E}\left[R_n \mid S_n = S^h, S_{n-1} = S^h\ \right]$$

giving

$$p_{hl} = \frac{r_l^+ - r_h^- - c_{hl}}{(r_h^+ - r_h^-) + (r_l^+ + r_l^-)} \tag{S20}$$

in this case, $p_{hl}$ represent the indifference environment for $S^l \to S^h$ state transition.

Note that for non-negligible adaptation costs, $p_{hl} < p_I <_{lh}$.

### S5 Environment inference and Cellular decision making

The cellular decision to reside in an optimal cell state depends on an estimate, $\pi_n$, of the $E^h$ environment occurrence probability $p$. The cell makes this estimate based on its cellular memory. The estimate $\pi_n$ is determined by the relative frequency of $E^h$ environmental experiences in the cellular memory of size $m$, i.e.,

$$\pi_n = \frac{K_n}{m}$$

Assuming that the cell selects its state to maximize the average fitness in next time step, the subsequent cell state is determined based on comparing the current environment estimate $\pi_n$ to the indifference environment $p_I$ as follows:

$$S_n = \begin{cases} S^h, & \pi_{n-1} > p_I \implies \mathbb{E}\left[R_n|S_n = S^h\right] > \mathbb{E}\left[R_n|S_n = S^l\right]; \\ \\ S^l, & \pi_{n-1} \leq p_I \implies \mathbb{E}\left[R_n|S_n = S^h\right] \leq \mathbb{E}\left[R_n|S_n = S^l\right]. \end{cases}$$

We note that the above inequality between $\pi_n$ and $p_I$ implies a preference of phenotypic state based on the corresponding inequality of expected fitness in each cell state (derived from eq. (S16)), which this model assumes drives cellular adaptation behavior.

Under costly adaptations, the cellular decision-making is modified as follows:

$$S_n = \begin{cases} S^h, & \begin{array}{l} S_{n-1} = S^h,\ \pi_{n-1} > p_{hl} \implies \mathbb{E}\left[R_n|S_n = S^h\right] > \mathbb{E}\left[R_n|S_n = S^l\right] - c_{hl}; \\ \qquad \textit{or} \\ S_{n-1} = S^l,\ \pi_{n-1} > p_{lh} \implies \mathbb{E}\left[R_n|S_n = S^h\right] - c_{lh} > \mathbb{E}\left[R_n|S_n = S^l\right]; \end{array} \\ \\ S^l, & \begin{array}{l} S_{n-1} = S^h,\ \pi_{n-1} \leq p_{hl} \implies \mathbb{E}\left[R_n|S_n = S^h\right] \leq \mathbb{E}\left[R_n|S_n = S^l\right] - c_{hl}; \\ \qquad \textit{or} \\ S_{n-1} = S^l,\ \pi_{n-1} \leq p_{lh} \implies \mathbb{E}\left[R_n|S_n = S^h\right] - c_{lh} \leq \mathbb{E}\left[R_n|S_n = S^l\right]; \end{array} \end{cases} \tag{S21}$$

Again, the implications above is the cell's projection of the average fitness using the estimate $\pi_{n-1}$ of the environment and previous cell state $S_{n-1}$ (eq. (S18)).

### S6 Dynamics of cell state

#### S6.1 Dynamics of cell state under stochastic environment

To capture the dynamics of costly cell state, ( $\mathbb{P}\left(S_n = S^h\right)$) we first model the dynamics of cell memory states that map to cell states $S^l$ and $S^h$. To do this, we start by partitioning the cell memory states as follows:

$$\hat{k} = \begin{Bmatrix} b_1 & b_{2,l} & b_{2,h} & b_3 \end{Bmatrix}$$

where,

$$\begin{aligned} b_1 &= \{0, 1, ..., k_{hl} - 1\} \\ b_{2,l} &= \{k_{hl}, k_{hl} + 1, ..., k_{lh} - 1\} \\ b_{2,h} &= \{k_{hl}, k_{hl} + 1, ..., k_{lh} - 1\} \\ b_3 &= \{k_{lh}, k_{lh} + 1, ..., m\} \end{aligned}$$

Here, $k_{lh} = \lceil p_{lh}\, m \rceil$ and $k_{hl} = \lceil p_{hl}\, m \rceil$ are the minimal and maximal number of $E^h$ environmental encounters required to stay in $S^h$ and $S^l$ cell states, respectively. Although, the sets $b_{2,l}$ and $b_{2,h}$ having overlapping cell memory states, they map distinctly to cell state $S^l$ and $S^h$, respectively, due to the presence of hysteresis in cellular decision making with non-negligible adaptation cost. Thus, the joint set $\{b_1\ b_{2,l}\}$ contains cell memory states that map to cell state $S^l$, while the joint set $\{b_{2,h}\ b_3\}$ contains cell memory states that map to cell state $S^h$.

The dynamics of cell memory within and across the above defined partitioning of states can be modeled as the following Markov Chain:

$$\begin{bmatrix}\boldsymbol{y}_{b_1} & \boldsymbol{y}_{b_{2,l}} & \boldsymbol{y}_{b_{2,h}} & \boldsymbol{y}_{b_3}\end{bmatrix}_{n+1} = \begin{bmatrix}\boldsymbol{y}_{b_1} & \boldsymbol{y}_{b_{2,l}} & \boldsymbol{y}_{b_{2,h}} & \boldsymbol{y}_{b_3}\end{bmatrix}_{n} \hat{P} \tag{S22}$$

where elements of vectors $[\boldsymbol{y}_{b_1,n}]$, $[\boldsymbol{y}_{b_{2,l},n}]$, $[\boldsymbol{y}_{b_{2,h},n}]$, and $[\boldsymbol{y}_{b_3,n}]$ are the probabilities of cell memory being in states of subsets $b_1$, $b_{2,l}$, $b_{2,h}$, and $b_3$, resp., at time $n$. In other words,

$$\begin{aligned}
\boldsymbol{y}_{b_1,n} &= \begin{bmatrix}y_{0,l} & y_{0,h} & y_{1,l} & y_{2,h}\cdots & y_{k_{hl}-1,l} & y_{k_{hl}-1,h}\end{bmatrix}_n \\
\boldsymbol{y}_{b_{2,l},n} &= \begin{bmatrix}y_{k_{hl},l} & y_{k_{hl},h} & y_{k_{hl}+1,l} & y_{k_{hl}+1,h} & \cdots & y_{k_{lh}-1,l} & y_{k_{lh}-1,h}\end{bmatrix}_n \\
\boldsymbol{y}_{b_{2,h},n} &= \begin{bmatrix}y'_{k_{hl},l} & y'_{k_{hl},h} & y'_{k_{hl}+1,l} & y'_{k_{hl}+1,h} & \cdots & y'_{k_{lh}-1,l} & y'_{k_{lh}-1,h}\end{bmatrix}_n \\
\boldsymbol{y}_{b_3,n} &= \begin{bmatrix}y_{k_{lh},l} & y_{k_{lh},h} & y_{k_{lh}+1,l} & y_{k_{lh}+1,h} & \cdots & y_{m-1,l} & y_{m-1,h} & y_{m,l} & y_{m,h}\end{bmatrix}_n
\end{aligned}$$

Above, we have defined cell memory state probabilities ($y_{k,e}$ with $e \in \{l, h\}$) depending on whether they were achieved after experiencing $E^l$ or $E^h$ environment.

The STM $\hat{P}$ above is defined as:

$$\hat{P} = \begin{bmatrix} P_{b_1} & P_{b_1 b_{2,l}} & 0 & 0 \\ P_{b_{2,l} b_1} & P_{b_{2,l}} & 0 & P_{b_{2,l} b_3} \\ P_{b_{2,h} b_1} & 0 & P_{b_{2,h}} & P_{b_{2,h} b_3} \\ 0 & 0 & P_{b_3 b_{2,h}} & P_{b_3} \end{bmatrix}$$

where,

$$\begin{aligned}
P_{b_1} &= \{P_{ij};\ i < 2k_{hl}+1,\ j < 2k_{hl}+1\} = \ \mathbb{P}\left(S_n = S^l \mid S_{n-1} = S^l\right) \\
P_{b_{2,l}} &= \{P_{ij};\ 2k_{hl}+1 \le i < 2k_{lh}+1,\ 2k_{hl}+1 \le j < 2k_{lh}+1\} = \ \mathbb{P}\left(S_n = S^l \mid S_{n-1} = S^l\right) \\
P_{b_{2,h}} &= \{P_{ij};\ 2k_{hl}+1 \le i < 2k_{lh}+1,\ 2k_{hl}+1 \le j < 2k_{lh}+1\} = \ \mathbb{P}\left(S_n = S^h \mid S_{n-1} = S^h\right) \\
P_{b_3} &= \{P_{ij};\ 2k_{lh}+1 \le i,\ 2k_{lh}+1 \le j\} = \ \mathbb{P}\left(S_n = S^h \mid S_{n-1} = S^h\right) \\
P_{b_1 b_{2,l}} &= \{P_{ij};\ i < 2k_{hl}+1,\ 2k_{hl}+1 \le j < 2k_{lh}+1\} = \ \mathbb{P}\left(S_n = S^l \mid S_{n-1} = S^l\right) \\
P_{b_{2,l} b_1} &= \{P_{ij};\ 2k_{hl}+1 \le i < 2k_{lh}+1,\ j < 2k_{hl}+1\} = \ \mathbb{P}\left(S_n = S^l \mid S_{n-1} = S^l\right) \\
P_{b_3 b_{2,h}} &= \{P_{ij};\ 2k_{lh}+1 \le i,\ 2k_{hl}+1 \le j < 2k_{lh}+1\} = \ \mathbb{P}\left(S_n = S^h \mid S_{n-1} = S^h\right) \\
P_{b_{2,h} b_3} &= \{P_{ij};\ 2k_{hl}+1 \le i < 2k_{lh}+1,\ 2k_{lh}+1 \le j\} = \ \mathbb{P}\left(S_n = S^h \mid S_{n-1} = S^h\right) \\
P_{b_{2,l} b_3} &= \{P_{ij};\ 2k_{hl}+1 \le i < 2k_{lh}+1,\ 2k_{lh}+1 \le j\} = \ \mathbb{P}\left(S_n = S^h \mid S_{n-1} = S^l\right) \\
P_{b_{2,h} b_1} &= \{P_{ij};\ 2k_{hl}+1 \le i < 2k_{lh}+1,\ j < 2k_{hl}+1\} = \ \mathbb{P}\left(S_n = S^l \mid S_{n-1} = S^h\right)
\end{aligned}$$

where, $P$ is the STM for the joint update of cell memory state and environment (Eq. S3) and $P_{ij}$ denote its elements at position $\{i, j\}$, $i, j \in \{1, 2..., m, ..., 2m+2\}$.

Using the dynamics of cell memory states as defined in eq. (S22) we then quantify the dynamics of cell state $S^h$ (by mapping the sum of the memory state vector to the cell state as done before) while accounting for the fact that the cell memory state at time $n$ determines the cell state at time $n+1$, $S_{n+1}$:

$$\mathbb{P}\left(S_{n+1} = S^h\right) = \left|\left|\begin{bmatrix}\boldsymbol{y}_{b_{2,h}} & \boldsymbol{y}_{b_3}\end{bmatrix}_n\right|\right|_1 = 1 - \ \left|\left|\begin{bmatrix}\boldsymbol{y}_{b_1} & \boldsymbol{y}_{b_{2,l}}\end{bmatrix}_n\right|\right|_1$$

In the above, $||\cdot||_1$ denotes the usual $\ell^1$ norm.

#### S6.2 Dynamics of cell state under periodic environment

As we observed previously, the periodicity in the probability distribution of cell memory state $K_n$ in a periodic environment may lead to periodicity in the dynamics of the cell state (section S3.2). To capture the oscillating dynamics of cell state in a periodic environment, we construct STMs $\hat{M}_0$ and $\hat{M}_1$ from STMs $M_0$ and $M_1$, respectively, (section S3.2) following the same partitioning of cell memory states ($\{b_1\ b_{2,l}\ b_{2,h}\ b_3\}$) as for capturing cell state dynamics in stochastic environment (section S6.1):

$$\hat{M}_0 = \begin{bmatrix} M_{0,\,b_1} & M_{0,\,b_1 b_{2,l}} & 0 & 0 \\ M_{0,\,b_{2,l} b_1} & M_{0,\,b_{2,l}} & 0 & M_{0,\,b_{2,l} b_3} \\ M_{0,\,b_{2,h} b_1} & 0 & M_{0,\,b_{2,h}} & M_{0,\,b_{2,h} b_3} \\ 0 & 0 & M_{0,\,b_3 b_{2,h}} & M_{0,\,b_3} \end{bmatrix} \qquad \hat{M}_1 = \begin{bmatrix} M_{1,\,b_1} & M_{1,\,b_1 b_{2,l}} & 0 & 0 \\ M_{1,\,b_{2,l} b_1} & M_{1,\,b_{2,l}} & 0 & M_{1,\,b_{2,l} b_3} \\ M_{1,\,b_{2,h} b_1} & 0 & M_{1,\,b_{2,h}} & M_{1,\,b_{2,h} b_3} \\ 0 & 0 & M_{1,\,b_3 b_{2,h}} & M_{1,\,b_3} \end{bmatrix} \tag{S23}$$

The dynamics of the cell memory state is then given by:

$$\begin{bmatrix}\boldsymbol{x}_{b_1} & \boldsymbol{x}_{b_{2,l}} & \boldsymbol{x}_{b_{2,h}} & \boldsymbol{x}_{b_3}\end{bmatrix}_{n+1} = \begin{cases} \begin{bmatrix}\boldsymbol{x}_{b_1} & \boldsymbol{x}_{b_{2,l}} & \boldsymbol{x}_{b_{2,h}} & \boldsymbol{x}_{b_3}\end{bmatrix}_n \hat{M}_0, & n+1 = rT + l \ \ \&\ \ 1 \le l \le T_{E^l_{bias}} \\ \begin{bmatrix}\boldsymbol{x}_{b_1} & \boldsymbol{x}_{b_{2,l}} & \boldsymbol{x}_{b_{2,h}} & \boldsymbol{x}_{b_3}\end{bmatrix}_n \hat{M}_1, & n+1 = rT + l \ \ \&\ \ \left(T_{E^l_{bias}} < l \le T\right) \end{cases} \tag{S24}$$

where $l \in \{1, 2, 3, ..., T\}$, $r \in \{0, 1, 2, 3, ...\}$; the elements of vectors $[\boldsymbol{x}_{b_1,\,n}]$, $[\boldsymbol{x}_{b_{2,l},\,n}]$, $[\boldsymbol{x}_{b_{2,h},\,n}]$, and $[\boldsymbol{x}_{b_3,\,n}]$ are the probabilities of cell memory residing in states contained in the portioning subsets $b_1$, $b_{2,l}$, $b_{2,h}$, and $b_3$, respectively, at time $n$. In other words,

$$\begin{aligned} \boldsymbol{x}_{b_1,n} &= \begin{bmatrix} x_0 & x_1 & ... & x_{k_{hl}-1}\end{bmatrix}_n \\ \boldsymbol{x}_{b_{2,l},n} &= \begin{bmatrix} x_{k_{hl}} & x_{k_{hl}+1} & ... & x_{k_{lh}-1}\end{bmatrix}_n \\ \boldsymbol{x}_{b_{2,h},n} &= \begin{bmatrix} x'_{k_{hl}} & x'_{k_{hl}+1} & ... & x'_{k_{lh}-1}\end{bmatrix}_n \\ \boldsymbol{x}_{b_3,n} &= \begin{bmatrix} x_{k_{lh}} & x_{k_{lh}+1} & ... & x_{m-1} & x_m\end{bmatrix}_n \end{aligned}$$

Using the dynamics of cell memory states as defined in eq. (S24), we then quantify the dynamics of cell state $S^h$ (by mapping the memory state to cell state as done previously):

$$\mathbb{P}\left(S_{n+1} = S^h\right) = \left|\left|\begin{bmatrix}\boldsymbol{x}_{b_{2,h}} & \boldsymbol{x}_{b_3}\end{bmatrix}_n\right|\right|_1 = 1 - \left|\left|\begin{bmatrix}\boldsymbol{x}_{b_1} & \boldsymbol{x}_{b_{2,l}}\end{bmatrix}_n\right|\right|_1$$

### S7 Quantification of cell state transition probabilities

In order to calculate the central moments of cell fitness and analyze cell population dynamics we must quantify cell state transition probabilities:

$$t_{ll} = \mathbb{P}\left(S_n = S^l \mid S_{n-1} = S^l\right), \quad t_{lh} = \mathbb{P}\left(S_n = S^h \mid S_{n-1} = S^l\right) = 1 - t_{ll}$$

$$t_{hh} = \mathbb{P}\left(S_n = S^h \mid S_{n-1} = S^h\right), \quad t_{hl} = \mathbb{P}\left(S_n = S^l \mid S_{n-1} = S^h\right) = 1 - t_{hh}$$

#### S7.1 Stochastic Correlated Environments

To quantify $t_{hh}$, we first consider the probability vectors $\begin{bmatrix}\boldsymbol{y}_{\boldsymbol{b}_{2,h}} & \boldsymbol{y}_{\boldsymbol{b}_3}\end{bmatrix}_{n-2}$ that represent the probabilities of cell memory states/vectors associated with cell state $S^h$ a time $n-1$(section S6). We normalize the above probability vectors to obtain the condition for which the cell occupies state $S^h$ at time $n-1$:

$$\boldsymbol{w}_h = \frac{\begin{bmatrix}\boldsymbol{y}_{b_{2,h}} & \boldsymbol{y}_{b_3}\end{bmatrix}_{n-2}}{\left|\left|\begin{bmatrix}\boldsymbol{y}_{b_{2,h}} & \boldsymbol{y}_{b_3}\end{bmatrix}_{n-2}\right|\right|_1}$$

Next, with the above normalized probability vector $\boldsymbol{w_b}$, we may find the probabilities of staying in the $S^h$ state after a single time step:

$$t_{hh} = \mathbb{P}\left(S_n = S^h \mid S_{n-1} = S^h\right) = \left|\left| \boldsymbol{w_h} \begin{bmatrix} P_{b_{2,h}} & P_{b_{2,h}b_3} \\ P_{b_3 b_{2,h}} & P_{b_3} \end{bmatrix} \right|\right|_1$$

Note that above conditional probability is only evaluated when $P(S_{n-1} = S^h) = \left|\left| \begin{bmatrix} \boldsymbol{y}_{b_{2,h}} & \boldsymbol{y}_{b_3} \end{bmatrix}_{n-2} \right|\right|_1 > \epsilon = 10^{-9}$, otherwise we set $t_{hh} = 0$.

Similarly, to quantify $t_{ll}$, we first consider the probability vectors $\begin{bmatrix} \boldsymbol{y_{b_1}} & \boldsymbol{y_{b_{2,l}}} \end{bmatrix}_{n-2}$ that represent the probabilities of cell memory states/vectors associated with cell state $S^l$ at time $n-1$ (section S6). We normalize the above probability vectors to obtain the condition for which the cell occupies state $S^h$ at time $n-1$:

$$\boldsymbol{w}_l = \frac{\begin{bmatrix} \boldsymbol{y}_{b_1} & \boldsymbol{y}_{b_{2,l}} \end{bmatrix}_{n-2}}{\left|\left| \begin{bmatrix} \boldsymbol{y}_{b_1} & \boldsymbol{y}_{b_{2,l}} \end{bmatrix}_{n-2} \right|\right|_1}$$

Next, with the above normalized probability vector $\boldsymbol{w_l}$, we may find the probabilities of staying in the $S^l$ state after a single time step:

$$t_{ll} = \mathbb{P}\left(S_n = S^l \mid S_{n-1} = S^l\right) = \left|\left| \boldsymbol{w_l} \begin{bmatrix} P_{b_1} & P_{b_1 b_{2,l}} \\ P_{b_{2,l} b_1} & P_{b_{2,l}} \end{bmatrix} \right|\right|_1$$

Again note that above conditional probability is only evaluated when $P(S_{n-1} = S^l) = \left|\left| \begin{bmatrix} \boldsymbol{y}_{b_1} & \boldsymbol{y}_{b_{2,l}} \end{bmatrix}_{n-2} \right|\right|_1 > \epsilon = 10^{-9}$, otherwise we set $t_{ll} = 0$.

Using the above calculations, we also determine the following joint probabilities of inter-temporal cell states, with or without environment state, which are later required in the quantification of central moments of cell fitness:

$$\begin{aligned} \mathbb{P}\left(S_n = S^l, S_{n-1} = S^l\right) &= \mathbb{P}\left(S_n = S^l \mid S_{n-1} = S^l\right) \mathbb{P}\left(S_{n-1} = S^l\right) = t_{ll}\, \mathbb{P}\left(S_{n-1} = S^l\right) \\ \mathbb{P}\left(S_n = S^h, S_{n-1} = S^l\right) &= \mathbb{P}\left(S_n = S^h \mid S_{n-1} = S^l\right) \mathbb{P}\left(S_{n-1} = S^l\right) = (1 - t_{ll})\, \mathbb{P}\left(S_{n-1} = S^l\right) \\ \mathbb{P}\left(S_n = S^l, S_{n-1} = S^h\right) &= \mathbb{P}\left(S_n = S^l \mid S_{n-1} = S^h\right) \mathbb{P}\left(S_{n-1} = S^h\right) = (1 - t_{hh})\, \mathbb{P}\left(S_{n-1} = S^h\right) \\ \mathbb{P}\left(S_n = S^h, S_{n-1} = S^h\right) &= \mathbb{P}\left(S_n = S^h \mid S_{n-1} = S^h\right) \mathbb{P}\left(S_{n-1} = S^h\right) = t_{hh}\, \mathbb{P}\left(S_{n-1} = S^h\right) \end{aligned} \tag{S25}$$

$$\begin{aligned} \mathbb{P}\left(S_n = S^h, S_{n-1} = S^h, E_{n-1} = E^l\right) &= \mathbb{P}\left(S_{n-1} = S^h\right) \sum_{i=1}^{m-k_{hl}+1} \left( \boldsymbol{w_h} \begin{bmatrix} P_{b_{2,h}} & P_{b_{2,h}b_3} \\ P_{b_3 b_{2,h}} & P_{b_3} \end{bmatrix} \right)_{2i-1} \\ \mathbb{P}\left(S_n = S^h, S_{n-1} = S^h, E_{n-1} = E^h\right) &= \mathbb{P}\left(S_{n-1} = S^h\right) \sum_{i=1}^{m-k_{hl}+1} \left( \boldsymbol{w_h} \begin{bmatrix} P_{b_{2,h}} & P_{b_{2,h}b_3} \\ P_{b_3 b_{2,h}} & P_{b_3} \end{bmatrix} \right)_{2i} \end{aligned} \tag{S26}$$

with $i$ being the indexing variable which is used to selectively sample probabilities when the environment $E_{n-1}$ is $E^l$ (odd samples) and $E^h$ (even samples).

##### S7.2 Periodic Environments

The periodic switches between $E^l$ and $E^h$ environments lead to time-dependent changes in the transition probabilities and are quantified as follows:

$$t_{ll,\,n} = \begin{cases} \dfrac{\left[\boldsymbol{x}_{b_1} \;\; \boldsymbol{x}_{b_{2,l}}\right]_{n-2}}{\left|\left|\left[\boldsymbol{x}_{b_1} \;\; \boldsymbol{x}_{b_{2,l}}\right]_{n-2}\right|\right|_1} \begin{bmatrix} M_{1,\,b_1} & M_{1,\,b_1 b_{2,l}} \\ M_{1,\,b_{2,l} b_1} & M_{1,\,b_{2,l}} \end{bmatrix}, & n = rT + l \;\; \& \;\; (l = 1 \;\; or \;\; T_{E^l_{bias}} + 1 < l \leq T) \\ \\ \dfrac{\left[\boldsymbol{x}_{b_1} \;\; \boldsymbol{x}_{b_{2,l}}\right]_{n-2}}{\left|\left|\left[\boldsymbol{x}_{b_1} \;\; \boldsymbol{x}_{b_{2,l}}\right]_{n-2}\right|\right|_1} \begin{bmatrix} M_{0,\,b_1} & M_{0,\,b_1 b_{2,l}} \\ M_{0,\,b_{2,l} b_1} & M_{0,\,b_{2,l}} \end{bmatrix}, & n = rT + l \;\; \& \;\; 1 < l \leq T_{E^l_{bias}} + 1 \end{cases}$$

Similarly,

$$t_{hh,\,n} = \begin{cases} \dfrac{\left[\boldsymbol{x}_{b_{2,h}} \;\; \boldsymbol{x}_{b_3}\right]_{n-2}}{\left|\left|\left[\boldsymbol{x}_{b_{2,h}} \;\; \boldsymbol{x}_{b_3}\right]_{n-2}\right|\right|_1} \begin{bmatrix} M_{1,\,b_{2,h}} & M_{1,\,b_{2,h} b_3} \\ M_{1,\,b_3 b_{2,h}} & M_{1,\,b_3} \end{bmatrix}, & n = rT + l \;\; \& \;\; (l = 1 \;\; or \;\; T_{E^l_{bias}} + 1 < l \leq T) \\ \\ \dfrac{\left[\boldsymbol{x}_{b_{2,h}} \;\; \boldsymbol{x}_{b_3}\right]_{n-2}}{\left|\left|\left[\boldsymbol{x}_{b_{2,h}} \;\; \boldsymbol{x}_{b_3}\right]_{n-2}\right|\right|_1} \begin{bmatrix} M_{0,\,b_{2,h}} & M_{0,\,b_{2,h} b_3} \\ M_{0,\,b_3 b_{2,h}} & M_{0,\,b_3} \end{bmatrix}, & n = rT + l \;\; \& \;\; 1 < l \leq T_{E^l_{bias}} + 1 \end{cases}$$

and,

$$t_{lh,\,n} = 1 - t_{ll,\,n} \qquad t_{hl,\,n} = 1 - t_{hl,\,n}$$

where, $l \in \{1, 2, 3, ..., T\}$, $r \in \{0, 1, 2, 3, ...\}$; probability vectors $\left[\boldsymbol{x}_{b_1} \;\; \boldsymbol{x}_{b_{2,l}}\right]$ and $\left[\boldsymbol{x}_{b_{2,h}} \;\; \boldsymbol{x}_{b_3}\right]$ represent the probabilities of cell memory states associated with cell state $S^l$ and $S^h$, respectively, a time $n-1$ (section S6.2).

We note that above calculation of $t_{ll}$ holds only when $P(S_{n-1} = S^l) = \left|\left|\left[\boldsymbol{x}_{b_1} \;\; \boldsymbol{x}_{b_{2,l}}\right]_{n-2}\right|\right|_1 > \epsilon = 10^{-9}$, otherwise we set $t_{ll} = 0$. Similarly, the above calculation of $t_{hh}$ holds only when $P(S_{n-1} = S^h) = \left|\left|\left[\boldsymbol{x}_{b_{2,h}} \;\; \boldsymbol{x}_{b_3}\right]_{n-2}\right|\right|_1 > \epsilon = 10^{-9}$, otherwise we set $t_{hh} = 0$.

Further, the inter-temporal joint probabilities of cell state can be determined as described before (Eq. S25).

#### S8 Quantifying the central moments of cellular fitness

From Eq. S15 we can write down the cell fitness conditional on its state as follows,

$$\begin{aligned} R_{n,h} &= r_h^+ \, \mathbb{1}_{E_n=E^h} + r_h^- (1 - \mathbb{1}_{E_n=E^h}) - c_{lh}(1 - \mathbb{1}_{S_{n-1}=S^h}) \\ R_{n,l} &= r_l^+ (1 - \mathbb{1}_{E_n=E^h}) + r_l^- \, \mathbb{1}_{E_n=E^h}(1 - \mathbb{1}_{S_n=S^h}) - c_{hl} \mathbb{1}_{S_{n-1}=S^h} \end{aligned} \tag{S27}$$

Based on the following results:

$$\begin{aligned} \mathbb{E}\left[\mathbb{1}_{X_n=X^h}\right] = \mathbb{E}\left[(\mathbb{1}_{X_n=X^h})^k\right] &= \mathbb{P}\left(X_n = X^h\right) \\ \mathbb{E}\left[\mathbb{1}_{X_n=X^h} \, \mathbb{1}_{X_{n-1}=X^h}\right] &= \mathbb{P}\left(X_n = X^h, \, X_{n-1} = X^h\right) \end{aligned} \tag{S28}$$

The conditional expected fitness at time $n$ is then given by:

$$\overline{R}_{n,h} = r_h^+ p_n + r_h^-(1-p_n) - c_{lh}\left(\frac{\mathbb{P}\left(S_{n-1}=S^l,\, S_n=S^h\right)}{\mathbb{P}\left(S_n=S^h\right)}\right)$$
$$\overline{R}_{n,l} = r_l^+(1-p_n) + r_l^- p_n - c_{hl}\left(\frac{\mathbb{P}\left(S_{n-1}=S^h,\, S_n=S^l\right)}{\mathbb{P}\left(S_n=S^l\right)}\right)$$

where, the current and inter-temporal probabilities of cell states $\mathbb{P}\left(S_n=S^h\right)$, $\mathbb{P}\left(S_{n-1}=S^l,\, S_n=S^h\right)$, and $\mathbb{P}\left(S_{n-1}=S^h,\, S_n=S^h\right)$ are determined in the section S6 and section S7, and

for stochastic correlated environments

$$p_n = \begin{cases} \mathbb{P}\left(E_n=E^h|S_n=S^h\right) = \dfrac{\mathbb{P}\left(E_n=E^h, S_n=S^h\right)}{\mathbb{P}\left(S_n=S^h\right)}, & S_n=S^h \\ \\ \mathbb{P}\left(E_n=E^h|S_n=S^l\right) = \dfrac{\mathbb{P}\left(E_n=E^h, S_n=S^l\right)}{\mathbb{P}\left(S_n=S^l\right)}, & S_n=S^l \end{cases}$$

for periodic environments

$$p_n = \begin{cases} 0, & n = rT + l \ \ \& \ \ 1 \le l \le T_{E^l\,bias} \\ 1, & n = rT + l \ \ \& \ \ T_{E^l\,bias} < l \le T, \end{cases}$$

where, $l \in \{1,2,3,...,T\}$, $r \in \{0,1,2,3,...\}$;

**Average cell fitness**

$$\begin{aligned} \mathbb{E}\left[R_n\right] = \overline{R}_n &= \overline{R}_{n,l}\,\mathbb{P}\left(S_n=S^l\right) + \overline{R}_{n,h}\,\mathbb{P}\left(S_n=S^h\right) \\ &= r_l^+ + \mathbb{P}\left(E_n=E^h, S_n=S^l\right)(r_l^- - r_l^+) - c_{hl}\,\mathbb{P}\left(S_{n-1}=S^h,\, S_n=S^l\right) \\ &\quad + r_h^- + \mathbb{P}\left(E_n=E^h, S_n=S^h\right)(r_h^+ - r_h^-) - c_{lh}\,\mathbb{P}\left(S_{n-1}=S^l,\, S_n=S^h\right) \end{aligned}$$

for stochastic correlated environments

$$\begin{aligned} \mathbb{P}\left(E_n=E^h, S_n=S^h\right) =&\ \mathbb{P}\left(E_n=E^h, S_n=S^h \mid E_{n-1}=E^h\right)\,\mathbb{P}\left(E_{n-1}=E^h\right) + \\ &\ \mathbb{P}\left(E_n=E^h, S_n=S^h \mid E_{n-1}=E^l\right)\,\mathbb{P}\left(E_{n-1}=E^l\right) \\ =&\ \mathbb{P}\left(E_n=E^h \mid E_{n-1}=E^h\right)\,\mathbb{P}\left(S_n=S^h \mid E_{n-1}=E^h\right)\,\mathbb{P}\left(E_{n-1}=E^h\right) + \\ &\ \mathbb{P}\left(E_n=E^h \mid E_{n-1}=E^l\right)\,\mathbb{P}\left(S_n=S^h \mid E_{n-1}=E^l\right)\,\mathbb{P}\left(E_{n-1}=E^l\right) \end{aligned}$$

here,

$$\mathbb{P}\left(E_n=E^l\right) = \mathbb{P}\left(E_{n-1}=E^l\right) = \frac{p_2}{p_1+p_2},$$
$$\mathbb{P}\left(E_n=E^h\right) = \mathbb{P}\left(E_{n-1}=E^h\right) = \frac{p_1}{p_1+p_2},$$
$$\mathbb{P}\left(E_n=E^h \mid E_{n-1}=E^h\right) = 1-p_2,$$
$$\mathbb{P}\left(E_n=E^h \mid E_{n-1}=E^l\right) = p_1,$$

and,

$$\mathbb{P}\left(S_n=S^h \mid E_{n-1}=E^l\right) = \frac{\sum_{i=1}^{m-k_{hl}+1}[\boldsymbol{y}_{b_{2,h}}\ \ \boldsymbol{y}_{b_3}]_{2i-1,\,n-1}}{\sum_{i=1}^{m-k_{hl}+1}[\boldsymbol{y}_{b_{2,h}}\ \ \boldsymbol{y}_{b_3}]_{2i-1,\,n-1} + \sum_{i=1}^{k_{lh}-1}[\boldsymbol{y}_{b_1}\ \ \boldsymbol{y}_{b_{2,l}}]_{2i-1,\,n-1}}$$

$$\mathbb{P}\left(S_n=S^h \mid E_{n-1}=E^h\right) = \frac{\sum_{i=1}^{m-k_{hl}+1}[\boldsymbol{y}_{b_{2,h}}\ \ \boldsymbol{y}_{b_3}]_{2i,\,n-1}}{\sum_{i=1}^{m-k_{hl}+1}[\boldsymbol{y}_{b_{2,h}}\ \ \boldsymbol{y}_{b_3}]_{2i,\,n-1} + \sum_{i=1}^{k_{lh}-1}[\boldsymbol{y}_{b_1}\ \ \boldsymbol{y}_{b_{2,l}}]_{2i,\,n-1}}$$

where, the joint vectors $[\boldsymbol{y}_{b_1} \;\; \boldsymbol{y}_{b_{2,l}}]$ and $[\boldsymbol{y}_{b_{2,h}} \;\; \boldsymbol{y}_{b_3}]$ contain the probabilities of residing in the cell memory states $\{b_1 \; b_{2,l}\}$ and $\{b_{2,l} \; b_3\}$ associated with cell state $S^l$ and $S^h$, respectively, as defined before in section S6. The indices of the above joint vectors is represented by $i$. The summation of the odd indices of $[\boldsymbol{y}_{b_1} \;\; \boldsymbol{y}_{b_{2,l}}]$ represents the joint probability of experiencing $E^l$ at time $n-1$ and residing in cell state $S^l$ at time $n$, and the summation of its even indices represents the joint probability of experiencing $E^h$ at time $n-1$ and residing in cell state $S^l$ at time $n$. Similarly, the summation of the odd indices of $[\boldsymbol{y}_{b_{2,h}} \;\; \boldsymbol{y}_{b_3}]$ represents the joint probability of experiencing $E^l$ at time $n-1$ and residing in cell state $S^h$ at time $n$, and the summation of its even indices represents the joint probability of experiencing $E^h$ at time $n-1$ and residing in cell state $S^h$ at time $n$.

$$\begin{aligned}
\mathbb{P}\left(E_n = E^h, S_n = S^l\right) = \; & \mathbb{P}\left(E_n = E^h, S_n = S^l \mid E_{n-1} = E^h\right) \; \mathbb{P}\left(E_{n-1} = E^h\right) + \\
& \mathbb{P}\left(E_n = E^h, S_n = S^l \mid E_{n-1} = E^l\right) \; \mathbb{P}\left(E_{n-1} = E^l\right) \\
= \; & \mathbb{P}\left(E_n = E^h \mid E_{n-1} = E^h\right) \; \mathbb{P}\left(S_n = S^l \mid E_{n-1} = E^h\right) \; \mathbb{P}\left(E_{n-1} = E^h\right) + \\
& \mathbb{P}\left(E_n = E^h \mid E_{n-1} = E^l\right) \; \mathbb{P}\left(S_n = S^l \mid E_{n-1} = E^l\right) \; \mathbb{P}\left(E_{n-1} = E^l\right)
\end{aligned}$$

where, $\mathbb{P}\left(E_{n-1} = E^l\right)$, $\mathbb{P}\left(E_{n-1} = E^h\right)$, $\mathbb{P}\left(E_n = E^h \mid E_{n-1} = E^h\right)$, and $\mathbb{P}\left(E_n = E^h \mid E_{n-1} = E^l\right)$ are as defined before and

$$\begin{aligned}
\mathbb{P}\left(S_n = S^l \mid E_{n-1} = E^l\right) &= 1 - \; \mathbb{P}\left(S_n = S^h \mid E_{n-1} = E^l\right) \\
\mathbb{P}\left(S_n = S^l \mid E_{n-1} = E^h\right) &= 1 - \; \mathbb{P}\left(S_n = S^h \mid E_{n-1} = E^h\right)
\end{aligned}$$

for periodic environments

$$\begin{aligned}
\mathbb{P}\left(E_n = E^h, S_n = S^h\right) &= \; \mathbb{P}\left(E_n = E^h\right) \; \mathbb{P}\left(S_n = S^h\right) \\
\mathbb{P}\left(E_n = E^h, S_n = S^l\right) &= \; \mathbb{P}\left(E_n = E^h\right) \; \mathbb{P}\left(S_n = S^l\right)
\end{aligned}$$

**Variance of the fitness**

$$\mathbb{V}\mathrm{ar}\left(R_n\right) = \mathbb{E}\left[R_n^2\right] - \mathbb{E}\left[R_n\right]^2$$

with $\mathbb{E}\left[R_n\right]$ as defined above and

$$\begin{aligned}
\mathbb{E}\left[R_n^2\right] = & \; (r_l^+)^2 + a_1 \, p_n + a_2 \, P(S_n = S^h) + a_3 \; \mathbb{P}\left(E_n = E^h, S_n = S^h\right) + a_4 \; \mathbb{P}\left(S_n = S^h, S_{n-1} = S^h\right) \\
& + a_5 \; \mathbb{P}\left(E_n = E^h, S_{n-1} = S^h\right) + a_6 \; \mathbb{P}\left(S_n = S^h, S_{n-1} = S^h, E_n = E^h\right)
\end{aligned}$$

with

$$\begin{aligned}
a_1 &= (r_l^-)^2 - (r_l^+)^2 \\
a_2 &= (r_h^-)^2 + c_{hl}^2 + c_{lh}^2 - (r_l^+)^2 - 2r_h^- c_{lh} - 2r_l^+ c_{hl} \\
a_3 &= -(r_h^-)^2 - (r_l^-)^2 + (r_h^+)^2 + (r_l^+)^2 + 2r_h^- c_{lh} - 2r_h^+ c_{lh} \\
a_4 &= -c_{hl}^2 - c_{lh}^2 + 2r_h^- c_{lh} + 2r_l^+ c_{hl} \\
a_5 &= -2r_l^- c_{hl} + 2r_l^+ c_{hl} \\
a_6 &= -2r_h^- c_{lh} + 2r_l^- c_{hl} + 2r_h^+ c_{lh} - 2r_l^+ c_{hl}
\end{aligned}$$

for stochastic correlated environments

$$
\begin{aligned}
\mathbb{P}\left(E_n = E^h, S_{n-1} = S^h\right) = \ & \mathbb{P}\left(E_n = E^h, S_{n-1} = S^h \mid E_{n-2} = E^h\right) \ \mathbb{P}\left(E_{n-2} = E^h\right) \\
& + \mathbb{P}\left(E_n = E^h, S_{n-1} = S^h \mid E_{n-2} = E^l\right) \ \mathbb{P}\left(E_{n-2} = E^l\right) \\
= \ & \mathbb{P}\left(E_n = E^h \mid E_{n-2} = E^h\right) \ \mathbb{P}\left(S_{n-1} = S^h \mid E_{n-2} = E^h\right) \ \mathbb{P}\left(E_{n-2} = E^h\right) \\
& + \mathbb{P}\left(E_n = E^h \mid E_{n-2} = E^l\right) \ \mathbb{P}\left(S_{n-1} = S^h \mid E_{n-2} = E^l\right) \ \mathbb{P}\left(E_{n-2} = E^l\right) \\
= \ & \left((1-p_2)^2 + p_1 p_2\right) \mathbb{P}\left(S_{n-1} = S^h \mid E_{n-2} = E^h\right) \ \mathbb{P}\left(E_{n-2} = E^h\right) \\
& + \left((1-p_1)p_1 + p_1(1-p_2)\right) \mathbb{P}\left(S_{n-1} = S^h \mid E_{n-2} = E^l\right) \ \mathbb{P}\left(E_{n-2} = E^l\right)
\end{aligned}
$$

$$
\begin{aligned}
& \mathbb{P}\left(S_n = S^h, S_{n-1} = S^h, E_n = E^h\right) \\
& = \ \mathbb{P}\left(E_n = E^h \mid S_n = S^h, S_{n-1} = S^h\right) \ \mathbb{P}\left(S_n = S^h, S_{n-1} = S^h\right) \\
& = \Big( \ \mathbb{P}\left(E_n = E^h \mid S_n = S^h, S_{n-1} = S^h, E_{n-1} = E^h\right) \ \mathbb{P}\left(E_{n-1} = E^h \mid S_n = S^h, S_{n-1} = S^h\right) \\
& + \ \mathbb{P}\left(E_n = E^h \mid S_n = S^h, S_{n-1} = S^h, E_{n-1} = E^l\right) \ \mathbb{P}\left(E_{n-1} = E^l \mid S_n = S^h, S_{n-1} = S^h\right) \Big) \ \mathbb{P}\left(S_n = S^h, S_{n-1} = S^h\right) \\
& = (1-p_2) \, \mathbb{P}\left(E_{n-1} = E^h, S_n = S^h, S_{n-1} = S^h\right) + p_1 \, \mathbb{P}\left(E_{n-1} = E^l, S_n = S^h, S_{n-1} = S^h\right)
\end{aligned}
$$

where $\mathbb{P}\left(E_{n-1} = E^h, S_n = S^h, S_{n-1} = S^h\right)$ and $\mathbb{P}\left(E_{n-1} = E^l, S_n = S^h, S_{n-1} = S^h\right)$ are quantified as formulated before in Eq. S26.
for periodic environments

$$
\mathbb{P}\left(E_n = E^h, S_{n-1} = S^h\right) = \ \mathbb{P}\left(E_n = E^h\right) \ \mathbb{P}\left(S_{n-1} = S^h\right)
$$

$$
\mathbb{P}\left(S_n = S^h, S_{n-1} = S^h, E_n = E^h\right) = \ \mathbb{P}\left(S_n = S^h, S_{n-1} = S^h\right) \ \mathbb{P}\left(E_n = E^h\right)
$$

### S9 Quantification of correlation between environment and cell state

$$
Corr(E_n, S_n) = \frac{\mathbb{E}\left[\mathbb{1}_{E_n=E^h} \mathbb{1}_{S_n=S^h}\right] - \mathbb{E}\left[\mathbb{1}_{E_n=E^h}\right] \mathbb{E}\left[\mathbb{1}_{S_n=S^h}\right]}{\mathbb{V}\text{ar}\left(\mathbb{1}_{E_n=E^h}\right) \mathbb{V}\text{ar}\left(\mathbb{1}_{S_n=S^h}\right)}
$$

using the result from the Eq. S28 we have

$$
Corr(E_n, S_n) = \frac{\mathbb{P}\left(E_n = E^h, S_n = S^h\right) - \ \mathbb{P}\left(E_n = E^h\right) \ \mathbb{P}\left(S_n = S^h\right)}{\Big( \ \mathbb{P}\left(E_n = E^h\right)\left(1 - \ \mathbb{P}\left(E_n = E^h\right)\right)\Big)\Big( \ \mathbb{P}\left(S_n = S^h\right)\left(1 - \ \mathbb{P}\left(S_n = S^h\right)\right)\Big)}
$$

### S10 Population Dynamics

So far, we have described how cell memory updates at discrete time steps influence the cell state, and the match or mismatch of the cell state with the environment then alters the cell fitness. Now, if we consider a population of cells, with each cell making an independent decision to reside in either cell states based on its memory, then we could capture the dynamic change in the mean population size using a discrete time Markov chain model by adapting the method developed in [1] as follows:

$$\begin{bmatrix} N_l \\ N_h \\ N_d \end{bmatrix}_{n+1} = \left( \begin{bmatrix} \alpha_{ll} & \alpha_{hl} & 0 \\ \alpha_{lh} & \alpha_{hh} & 0 \\ \alpha_{ld} & \alpha_{hd} & 1 \end{bmatrix}_n \times \begin{bmatrix} 1+\overline{R}_l \mathbb{1}_{\overline{R}_l \geq 0} & 0 & 0 \\ 0 & 1+\overline{R}_h \mathbb{1}_{\overline{R}_h \geq 0} & 0 \\ 0 & 0 & 1 \end{bmatrix}_n \right) \begin{bmatrix} N_l \\ N_h \\ N_d \end{bmatrix}_n \tag{S29}$$

where, $N_i$'s with $i \in \{l, h, d\}$ are the mean number of cells in states $S^l$, $S^h$, and death, '$d$', cell compartment; $\alpha_{ij}$'s with $i, j \in \{l, h, d\}$ are the transition probabilities between states $S^l$, $S^h$ and death cell compartments; $\overline{R}_l$ and $\overline{R}_h$ are conditional expected fitness of residing in cell states $S^l$ and $S^h$, respectively, as described in the previous subsection and define the fraction of cells in each state that divide in a step time. The elements of the above state transition matrix must follow the below inequalities:

$$\sum_{j=l,h,d} \alpha_{ij} = 1, \quad i = l, h, d; \qquad |\overline{R}_i| \leq 1 \quad i \in \{l, h\} \tag{S30}$$

The above transition probabilities $\alpha_{ij}$ are defined as follows:

$$\begin{aligned}
\alpha_{ld} &= -\overline{R}_l \mathbb{1}_{\overline{R}_l < 0} \\
\alpha_{hd} &= -\overline{R}_h \mathbb{1}_{\overline{R}_h < 0} \\
\alpha_{ll} &= (1 + \overline{R}_l \mathbb{1}_{\overline{R}_l < 0})\, t_{ll} \\
\alpha_{lh} &= (1 + \overline{R}_l \mathbb{1}_{\overline{R}_l < 0})\, t_{lh} \\
\alpha_{hh} &= (1 + \overline{R}_h \mathbb{1}_{\overline{R}_h < 0})\, t_{hh} \\
\alpha_{hl} &= (1 + \overline{R}_h \mathbb{1}_{\overline{R}_h < 0})\, t_{hl}
\end{aligned}$$

We remark that the cellular fitness are interpreted as either fraction of cells that divide or probabilities to transition into the death compartment. Therefore, to satisfy the condition in Eq S30, the unit time step in the model needs to be smaller the fastest cell division and death timescales $1/r_l^+$, $1/r_h^+$, $1/r_l^-$ or $1/r_h^-$, and accordingly memory size has to be modulated.

For two (or more) cell subpopulations with distinct adaptation costs, the mean cell counts dynamics can be captured as follows:

$$\begin{bmatrix} N_{l,1} \\ N_{h,1} \\ N_{l,2} \\ N_{h,2} \\ N_d \end{bmatrix}_{n+1} = \left( \begin{bmatrix} \alpha_{ll,1} & \alpha_{hl,1} & 0 & 0 & 0 \\ \alpha_{lh,1} & \alpha_{hh,1} & 0 & 0 & 0 \\ 0 & 0 & \alpha_{ll,2} & \alpha_{hl,2} & 0 \\ 0 & 0 & \alpha_{hl,2} & \alpha_{hh,2} & 0 \\ \alpha_{ld,1} & \alpha_{hd,1} & \alpha_{ld,2} & \alpha_{hd,2} & 1 \end{bmatrix}_n \times \right.$$

$$\left. \begin{bmatrix} 1+\overline{R}_{l,1} \mathbb{1}_{\overline{R}_{l,1} \geq 0} & 0 & 0 & 0 & 0 \\ 0 & 1+\overline{R}_{h,1} \mathbb{1}_{\overline{R}_{h,1} \geq 0} & 0 & 0 & 0 \\ 0 & 0 & 1+\overline{R}_{l,2} \mathbb{1}_{\overline{R}_{l,2} \geq 0} & 0 & 0 \\ 0 & 0 & 0 & 1+\overline{R}_{h,2} \mathbb{1}_{\overline{R}_{h,2} \geq 0} & 0 \\ 0 & 0 & 0 & 0 & 1 \end{bmatrix}_n \right) \begin{bmatrix} N_{l,1} \\ N_{h,1} \\ N_{l,2} \\ N_{h,2} \\ N_d \end{bmatrix}_n$$

where $\alpha_{ij,k}$ and $\overline{R}_{i,k}$ with $k = \{1, 2\}$ represent the cell state transition probabilities and conditional state fitness, respectively, for the individual subpopulation $k$.

#### S11 Cell phenotypic residence times

To calculate the probability and statistical measures of time $T_h$ (resp. $T_l$) spent in cell state $S^h$ (resp. $S^l$) before switching to state $S^l$ (resp. $S^h$), our strategy is to partition the cell memory states, $k$, and State Transition Matrix, $P$, and probability distribution, $\boldsymbol{y}$, of the cell memory states (from section S3) as follows:

for sets:

$$k = \{0, 1, 2, ..., m-1, m\} = \begin{Bmatrix} b & b^c \end{Bmatrix}$$

STMs:

$$P = \begin{bmatrix} P_b & P_{bb^c} \\ P_{b^c b} & P_{b^c} \end{bmatrix}$$

and vectors:

$$\boldsymbol{y} = [y_{0,l} \;\; y_{0,h} \;\; y_{1,l} \;\; y_{1,h} \;\; ... \; y_{m-1,l} \;\; y_{m-1,h} \;\; y_{m,l} \;\; y_{m,h}] = \begin{bmatrix} \boldsymbol{y}_b & \boldsymbol{y}_{b^c} \end{bmatrix}$$

where,

$$\begin{aligned}
b &= \{0, 1, ..., k_I - 1\} \\
b^c &= \{k_I, k_I + 1, ..., m\} \\
P_b &= \{P_{ij};\ i < 2k_I + 1,\ j < 2k_I + 1\} = \mathbb{P}\left(S_n = S^l \mid S_{n-1} = S^l\right) \\
P_{bb^c} &= \{P_{ij};\ i < 2k_I + 1,\ j \geq 2k_I + 1\} = \mathbb{P}\left(S_n = S^h \mid S_{n-1} = S^l\right) \\
P_{b^c b} &= \{P_{ij};\ i \geq 2k_I + 1,\ j < 2k_I + 1\} = \mathbb{P}\left(S_n = S^l \mid S_{n-1} = S^h\right) \\
P_{b^c} &= \{P_{ij};\ i \geq 2k_I + 1,\ j \geq 2k_I + 1\} = \mathbb{P}\left(S_n = S^h \mid S_{n-1} = S^h\right) \\
\boldsymbol{y}_b &= \begin{bmatrix} y_{0,l} & y_{0,h} & y_{1,l} & y_{1,h} \cdots & y_{k_I-1,l} & y_{k_I-1,h} \end{bmatrix} \\
\boldsymbol{y}_{b^c} &= \begin{bmatrix} y_{k_I,l} & y_{k_I,h} & \cdots & y_{m-1,l} & y_{m-1,h} & y_{m,l} & y_{m,h} \end{bmatrix}
\end{aligned}$$

In the above, cell memory states present in subsets $b$ and $b^c$ map to cell states $S^l$ and $S^h$, respectively; $i$ and $j$ represent the indices of STM $P$; and $y_{k,e}$ denote the probability $\mathbb{P}\left(K_n = k, E_n = E^e\right)$.

A sojourn in cell state $S^l$ begins with cell memory states in $b$ with associated probabilities given by the vector $\boldsymbol{v}$:

$$\boldsymbol{v} = \begin{bmatrix} v_{0,l} & v_{0,h} & v_{1,l} & v_{1,h} & ... & v_{k_I-1,l} & v_{k_I-1,h} \end{bmatrix}$$

Now, as the transition from cell state $S^h$ to $S^l$ happens when $K_n$ decreases from $k_I$ to $k_I - 1$ on experiencing an $E^l$ environment, we have $v_{k_I-1,l} = 1$ with the remaining elements being zero.

Similarly, a sojourn in cell state $S^h$ begins with cell memory states in $b^c$ with associated probabilities given by the vector $\boldsymbol{w}$:

$$\boldsymbol{w} = \begin{bmatrix} w_{k_I,l} & w_{k_I,h} & ... & w_{m-1,l} & w_{m-1,h} & w_{m,l} & w_{m,h} \end{bmatrix}$$

Again, as the transition from cell state $S^l$ to $S^h$ happens when $K_n$ increments from $k_I - 1$ to $k_I$ on experiencing an $E^h$ environment, we have $w_{k_I,h} = 1$ with the remaining elements being zero.

The probability distribution of sojourn times $\tau_l$ and $\tau_h$ in cell states $S^l$ and $S^h$ can then be calculated as follows:

$$\begin{aligned}
\mathbb{P}\left(k_i \in b^c \mid k_m \in b : m \in \{0, 1, ..., i-1\}\right) &= \mathbb{P}\left(T_l = i\right) = \boldsymbol{v} P_b^{i-1} (I - P_b)\mathbf{1};\ i = 1, 2, 3, ... \\
\mathbb{P}\left(k_i \in b \mid k_m \in b^c : m \in \{0, 1, ..., i-1\}\right) &= \mathbb{P}\left(T_h = i\right) = \boldsymbol{w} P_{b^c}^{i-1} (I - P_{b^c})\mathbf{1};\ i = 1, 2, 3, ...
\end{aligned} \qquad \text{(S31)}$$

Here, $\mathbf{1}$ is the column vector of ones matching the length of the vector to its left (for example, matching length of vector $\boldsymbol{v}$ for $T_l$ calculation and length of vector $\boldsymbol{w}$ for $T_h$ calculation).

Given the probability distributions, we have statistical moments of the residence (sojourn) times in terms of factorial moments (chapter 5 of [5]):

$$\mathbb{FM}_j\left(T_l\right) = j!\,\boldsymbol{v}\,P_b^{j-1}\left(I - P_b\right)^{-j}\mathbf{1}$$
$$\mathbb{FM}_j\left(T_h\right) = j!\,\boldsymbol{w}\,P_{b^c}^{j-1}\left(I - P_{b^c}\right)^{-j}\mathbf{1}$$

for the $j^{th}$ order factorial moment. The above can be used to obtain the ordinary moments:

$$\begin{aligned} \mathbb{E}\left[T_l^n\right] &= \sum_{j=0}^{n}\binom{n}{j}\mathbb{FM}_j\left(T_l\right) \\ \mathbb{E}\left[T_h^n\right] &= \sum_{j=0}^{n}\binom{n}{j}\mathbb{FM}_j\left(T_h\right) \end{aligned} \tag{S32}$$

Under costly cellular adaptation, the differences between $p_{lh}$ and $p_{hl}$ results in overlap of the cell memory states to cell state $S^l$ and $S^l$. Despite this, residence times may be calculated in the same way as in the no-cost case with minimal modification to the critical cutoff $k_I$:

$$\tilde{k} = \begin{Bmatrix} b_1 & b_2 \end{Bmatrix} \qquad \tilde{P} = \begin{bmatrix} P_{b_1} & P_{b_1 b_2} \\ P_{b_2 b_1} & P_{b_2} \end{bmatrix} \qquad \tilde{\boldsymbol{y}} = \begin{bmatrix} \boldsymbol{y}_{b_1} & \boldsymbol{y}_{b_2} \end{bmatrix}$$

where,

$$\begin{aligned} b_1 &= \{0, 1, ..., k_{lh} - 1\} \\ b_2 &= \{k_{hl}, k_{hl} + 1, ..., m\} \\ P_{b_1} &= \{P_{ij};\ i < 2k_{lh} + 1,\ j < 2k_{lh} + 1\} = \mathbb{P}\left(S_n = S^l \mid S_{n-1} = S^l\right) \\ P_{b_1 b_2} &= \{P_{ij};\ i < 2k_{lh} + 1,\ j \geq 2k_{hl} + 1\} = \mathbb{P}\left(S_n = S^h \mid S_{n-1} = S^l\right) \\ P_{b_2 b_1} &= \{P_{ij};\ i \geq 2k_{hl} + 1,\ j < 2k_{lh} + 1\} = \mathbb{P}\left(S_n = S^l \mid S_{n-1} = S^h\right) \\ P_{b_2} &= \{P_{ij};\ i \geq 2k_{hl} + 1,\ j \geq 2k_{hl} + 1\} = \mathbb{P}\left(S_n = S^h \mid S_{n-1} = S^h\right) \\ \boldsymbol{y}_{b_1} &= \begin{bmatrix} y_{0,l} & y_{0,h} & y_{1,l} & y_{1,h} \cdots & y_{k_{lh}-1,l} & y_{k_{lh}-1,h} \end{bmatrix} \\ \boldsymbol{y}_{b_2} &= \begin{bmatrix} y_{k_h l,l} & y_{k_h l,h} & \cdots & y_{m-1,l} & y_{m-1,h} & y_{m,l} & y_{m,h} \end{bmatrix} \end{aligned}$$

Given these updated partitions, the probabilities, given by vector $\tilde{\boldsymbol{v}}$, of beginning a sojourn in cell state $S^l$ with cell memory states in $b_1$ now become:

$$\tilde{\boldsymbol{v}} = \begin{bmatrix} \tilde{v}_{0,l} & \tilde{v}_{0,h} \ldots \tilde{v}_{k_{hl}-1,l} & \tilde{v}_{k_{hl}-1,h} & \tilde{v}_{k_{hl},l} & \tilde{v}_{k_{hl},h} & \tilde{v}_{k_{hl}+1,l} & \tilde{v}_{k_{hl}+1,h} \ldots \tilde{v}_{k_{lh}-1,l} & \tilde{v}_{k_{lh}-1,h} \end{bmatrix}$$

Now, as the transition from cell state $S^h$ to $S^l$ happens when $K_n$ decrements from $k_{hl}$ to $k_{hl} - 1$ on experiencing an $E^l$ environment, we have $\tilde{v}_{k_{hl}-1,l} = 1$ with the remaining elements being zero.

Similarly, the probabilities, given by vector $\tilde{\boldsymbol{w}}$, of beginning a sojourn in cell state $S^h$ with cell memory states in $b_2$ now become:

$$\tilde{\boldsymbol{w}} = \begin{bmatrix} \tilde{w}_{k_{hl},l} & \tilde{w}_{k_{hl},h} \ldots \tilde{w}_{k_{lh}-1,l} & \tilde{w}_{k_{lh}-1,h} & \tilde{w}_{k_{lh},l} & \tilde{w}_{k_{lh},h} & \tilde{w}_{k_{lh}+1,l} & \tilde{w}_{k_{lh}+1,h} \ldots \tilde{w}_{m,l} & \tilde{w}_{m,h} \end{bmatrix}$$

Again, as the transition from cell state $S^l$ to $S^h$ happens when $K_n$ increments from $k_{lh} - 1$ to $k_{lh}$ on experiencing an $E^h$ environment, we have $\tilde{w}_{k_{lh},h} = 1$ with the remaining elements being zero.

With these updated partition matrices and probabilities for starting the next sojourn, we use the above formulae (eq. (S31) and eq. (S32)) to calculate the sojourn time probability distribution and its statistical measures.

### S12 Transient cellular responses

So far we have characterized the long-term response of cellular memory states for each model through their stationary distributions. We next determine the dynamics of the memory models on shorter times (time steps of the order of memory size $m$) by analyzing the time it takes for cells to adapt to environment shifts.

#### S12.1 Single abrupt shift in the environment

The mathematical descriptions in the previous sections describe how a cell adapts its state in a stationary environment, i.e., an environment whose occurrence probability $p$ is constant. Next, we considered scenarios where there is a discrete jump in $p$. Such scenarios represent, for example, abrupt changes in nutrient availability or a switch from normal growth conditions to the presence of cytotoxic/cytostatic drugs in the environment. We begin first by analyzing the time it takes for cells to restructure their memory such that the optimal phenotype to the current environment is attained.

Let $n \to n+1$ denote the time around which the environment switches from $p_1 \to p_2$ while preserving its autocorrelation $\rho_{EE}(1)$. To determine the mean adaptation time to optimal cell state on environment switch ($p_1 \to p_2$), we quantified the mean residence time in the suboptimal cell state. We used the formulation developed in section S11 for mean residence times while modifying the probability vectors $\boldsymbol{v}$ and $\boldsymbol{w}$ of starting a sojourn in states of subsets $b$ and $b^c$, respectively, as follows:

For $p_1 < p_I < p_2$ as $S^l$ is suboptimal after the environment switch, we calculate $\mathbb{E}[T_l]$ from eq. (S32) using:

$$\boldsymbol{v} = \frac{\boldsymbol{y}_b}{||\boldsymbol{y}_b||_1}$$

and, for $p_2 < p_I < p_1$ as $S^h$ is suboptimal state after the environment switch, we calculate $\mathbb{E}[T_h]$ from eq. (S32) using:

$$\boldsymbol{w} = \frac{\boldsymbol{y}_{b^c}}{||\boldsymbol{y}_{b^c}||_1}$$

where, $\boldsymbol{y}_b$ and $\boldsymbol{y}_{b^c}$ are stationary distributions of partitioned cell memory states $b$ and $b^c$ arising from cell memory under environment $p_1$ (section S3.1 and section S11).

The above calculation of mean adaptation times also apply to the costly adaptation case as well, provided that: 1) the partitioning of cell memory states/vectors $b$ and $b^c$ are replaced with $b_1$ and $b_2$, and; 2) making concomitant changes in the partitioning of STM P and quantification of vectors $\boldsymbol{v}$ and $\boldsymbol{w}$ (section S11).

#### S12.2 Periodic shifts in the environment

The probabilistic erasure of past environmental experiences in the cellular memory retains the stochastic update of the cell memory state, $K_n$, even under periodic (and therefore also deterministic) environments (section S3.2). Thus, owing to the stochastic update dynamics of cell memory, we quantified mean residence time in the suboptimal cell state to determine the mean adaptation time to the optimal cell state upon an environmental switch. When the environment undergoes transition $E^l \to E^h$, the mean residence time in $S^l$, $\mathbb{E}[T_l]$, characterizes the mean adaptation time to the $S^h$ state (preferred in the $E^h$ environment). Similarly, when the environment undergoes transition $E^h \to E^l$, the mean residence time in $S^h$, $\mathbb{E}[T_h]$, characterizes the mean adaptation time to the $S^h$ state (preferred in the $E^l$ environment).

To determine probability distribution of residence time $T_l$, and thereby adaptation times, to cell state $S^h$ during a periodic transition $E^l \to E^h$, we partition the cell memory states, $k$, and State Transition Matrix, $M_0$ and $M_1$, and probability distribution, $\boldsymbol{x}$, of the cell memory states (from section S3.2) as follows:

cell memory states:

$$k = \{0, 1, 2, ..., m-1, m\} = \begin{Bmatrix} b & b^c \end{Bmatrix}$$

and the probability vector:

$$\boldsymbol{x} = [x_0 \;\; x_1 \;\; ... \; x_{m-1} \;\; x_m] = \begin{bmatrix} \boldsymbol{x}_b & \boldsymbol{x}_{b^c} \end{bmatrix}$$

are partitioned as follows:

$$\begin{aligned}
b &= \{0, 1, ..., k_I - 1\} \\
b^c &= \{k_I, k_I + 1, ..., m\} \\
M_{e,b} &= \{M_{e,ij};\; i < k_{lh},\; j < k_{lh}\} = \; \mathbb{P}\left(S_n = S^l \mid S_{n-1} = S^l\right) \\
M_{e,bb^c} &= \{M_{e,ij};\; i < k_{lh},\; j \geq k_{lh}\} = \; \mathbb{P}\left(S_n = S^h \mid S_{n-1} = S^l\right) \\
M_{e,b^cb} &= \{M_{e,ij};\; i \geq k_{hl},\; j < k_{hl}\} = \; \mathbb{P}\left(S_n = S^l \mid S_{n-1} = S^h\right) \\
M_{e,b^c} &= \{M_{e,ij};\; i \geq k_{hl},\; j \geq k_{hl}\} = \; \mathbb{P}\left(S_n = S^h \mid S_{n-1} = S^h\right) \\
\boldsymbol{x}_b &= \begin{bmatrix} x_0 & x_1 & ... & x_{k_{lh}-1} \end{bmatrix} \\
\boldsymbol{x}_{b^c} &= \begin{bmatrix} x_{k_{hl}} & ... & x_{m-1} & x_m \end{bmatrix}
\end{aligned}$$

with the transition between above partitioned states are tracked by following STMs:

$$\begin{aligned}
M_{e,b} &= \{M_{e,ij};\; i < k_{lh},\; j < k_{lh}\} = \; \mathbb{P}\left(S_n = S^l \mid S_{n-1} = S^l\right) \\
M_{e,bb^c} &= \{M_{e,ij};\; i < k_{lh},\; j \geq k_{lh}\} = \; \mathbb{P}\left(S_n = S^h \mid S_{n-1} = S^l\right) \\
M_{e,b^cb} &= \{M_{e,ij};\; i \geq k_{hl},\; j < k_{hl}\} = \; \mathbb{P}\left(S_n = S^l \mid S_{n-1} = S^h\right) \\
M_{e,b^c} &= \{M_{e,ij};\; i \geq k_{hl},\; j \geq k_{hl}\} = \; \mathbb{P}\left(S_n = S^h \mid S_{n-1} = S^h\right)
\end{aligned} \tag{S33}$$

where, $e \in \{0, 1\}$.

Next, to consider that the cell resides in cell state $S^l$ and $S^h$ by the end of $E^l$ and $E^h$ environment cycles, we normalize the probabilities associated with their cell memory states $\boldsymbol{x}_b$ and $\boldsymbol{x}_{b^c}$, respectively:

$$\boldsymbol{v} = \frac{\boldsymbol{x}_b(rT + T_{E^l_{bias}})}{||\boldsymbol{x}_b(rT + T_{E^l_{bias}})||_1}$$

$$\boldsymbol{w} = \frac{\boldsymbol{x}_{b^c}(rT + T)}{||\boldsymbol{x}_{b^c}(rT + T)||_1}$$

where, $\boldsymbol{x}_b(T_{E^l_{bias}})$ and $\boldsymbol{x}_b(T)$ refer to evaluation of $\boldsymbol{x}(rT+l)$ in eq. (S10) at $l = T_{E^l_{bias}}$ and $l = T$, respectively.

the quantification of probabilities of residence times in $S^h$ state when environment cycle begins with $E^l$ states are as follows:

$$P(T_h = rT + l) = \begin{cases} \boldsymbol{w}\, M_{0,b}^{(l-1)}\, M_{0,bb^c}, & r = 0 \,\&\, l \leq T_{E^l_{bias}} \\ \boldsymbol{w}\, M_{0,b}^{T_{E^l_{bias}}}\, M_{1,b}^{(l-T_{E^l_{bias}}-1)}\, M_{1,bb^c}, & r = 0 \,\&\, l > T_{E^l_{bias}} \\ \boldsymbol{w}\, M_{0,b}^{T_{E^l_{bias}}}\, M_{1,b}^{T_{E^h_{bias}}}\, (M_{0,b}^{T_{E^l_{bias}}}\, M_{1,b}^{T_{E^h_{bias}}})^{(r-1)}\, M_{0,b}^{l-1}\, M_{0,bb^c}, & r > 0 \,\&\, l \leq T_{E^l_{bias}} \\ \boldsymbol{w}\, M_{0,b}^{T_{E^l_{bias}}}\, M_{1,b}^{T_{E^h_{bias}}}\, (M_{0,b}^{T_{E^l_{bias}}}\, M_{1,b}^{T_{E^h_{bias}}})^{(r-1)}\, M_{0,b}^{T_{E^l_{bias}}}\, M_{1,b}^{(l-T_{E^l_{bias}}-1)}\, M_{1,bb^c}, & r > 0 \,\&\, l > T_{E^l_{bias}} \end{cases}$$

where $l \in \{1, 2, 3, ..., T\}$, $r \in \{0, 1, 2, 3, ...\}$.

Similarly, the quantification of probabilities of residence times in $S^l$ state when environment cycle begins with $E^h$ states are as follows:

$$P(T_l = rT + l) = \begin{cases} \boldsymbol{v}\, M_{1,b}^{(l-1)}\, M_{1,bb^c}, & r = 0 \;\&\; l \leq T_{E^h_{bias}} \\ \boldsymbol{v}\, M_{1,b}^{T_{E^h_{bias}}}\, M_{0,b}^{(l-T_{E^h_{bias}}-1)}\, M_{0,bb^c}, & r = 0 \;\&\; l > T_{E^h_{bias}} \\ \boldsymbol{v}\, M_{1,b}^{T_{E^h_{bias}}}\, M_{0,b}^{T_{E^l_{bias}}}\, (M_{1,b}^{T_{E^h_{bias}}}\, M_{0,b}^{T_{E^l_{bias}}})^{(r-1)}\, M_{1,b}^{l-1}\, M_{1,bb^c}, & r > 0 \;\&\; l \leq T_{E^h_{bias}} \\ \boldsymbol{v}\, M_{1,b}^{T_{E^h_{bias}}}\, M_{0,b}^{T_{E^l_{bias}}}\, (M_{1,b}^{T_{E^h_{bias}}}\, M_{0,b}^{T_{E^l_{bias}}})^{(r-1)}\, M_{1,b}^{T_{E^h_{bias}}}\, M_{0,b}^{(l-T_{E^h_{bias}}-1)}\, M_{0,bb^c}, & r > 0 \;\&\; l > T_{E^h_{bias}} \end{cases}$$

where, $l \in \{1, 2, 3, ..., T\}$ and $r \in \{0, 1, 2, 3, ...\}$

The above calculation of mean adaptation time follow under zero adaptation cost adaptation as well, with appropriate changes in the partitioning of cell memory states and STMs for $k_{hl} = k_{lh} = k_I$.

### References


[1] Vimalathithan Devaraj and Biplab Bose. Morphological state transition dynamics in egf-induced epithelial to mesenchymal transition. *Journal of clinical medicine*, 8(7):911, 2019.

[2] Guillaume Lambert and Edo Kussell. Memory and fitness optimization of bacteria under fluctuating environments. *PLoS genetics*, 10(9):e1004556, 2014.

[3] Jingxin Li, Pavithran T Ravindran, Aoife O'Farrell, Gianna T Busch, Ryan H Boe, Zijian Niu, Sean Woo, Margaret C Dunagin, Naveen Jain, Yogesh Goyal, et al. Ap-1 mediates cellular adaptation and memory formation during therapy resistance. *bioRxiv*, 2024.

[4] Angela Oliveira Pisco, Amy Brock, Joseph Zhou, Andreas Moor, Mitra Mojtahedi, Dean Jackson, and Sui Huang. Non-darwinian dynamics in therapy-induced cancer drug resistance. *Nature communications*, 4(1):2467, 2013.

[5] Gerardo Rubino and Bruno Sericola. *Markov chains and dependability theory.* Cambridge University Press, 2014.

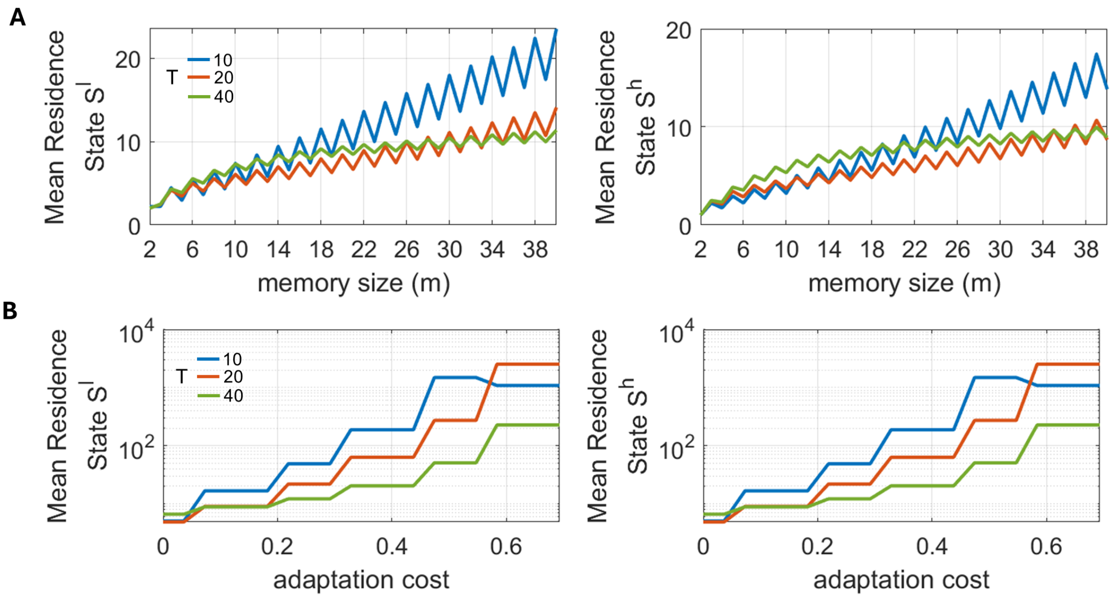


Figure S1: **Effects of memory size and adaptation cost on the mean residence times in cell states $S^l$ and $S^h$.** By conditioning the cell to reside in the $S^l$ state by the end of the $E^l$ environment cycle, the mean residence time in $S^l$ state determines how long it takes for the cell to switch its state during the current $E^h$ environment cycle and subsequent cycles of the oscillatory environments. Similarly, by conditioning the cell to reside in $S^h$ state by the end of the $E^h$ environment cycle, the mean residence time in $S^h$ state determines how long it takes for the cell to switch its state during the current $E^l$ environment cycle and subsequent cycles of the oscillatory environments. Refer to SI section S12.2 for the analytical formulation of the performed calculation. In panel A, adaptation cost $c_{lh} = c_{hl} = 0$; and in panel B, memory size, $m = 11$. The fitness values for (mis-)matching environment and cell state are $r_h^- = 0.3$, $r_h^+ = 0.8$, $r_l^- = 0.1$, and $r_l^+ = 1$.

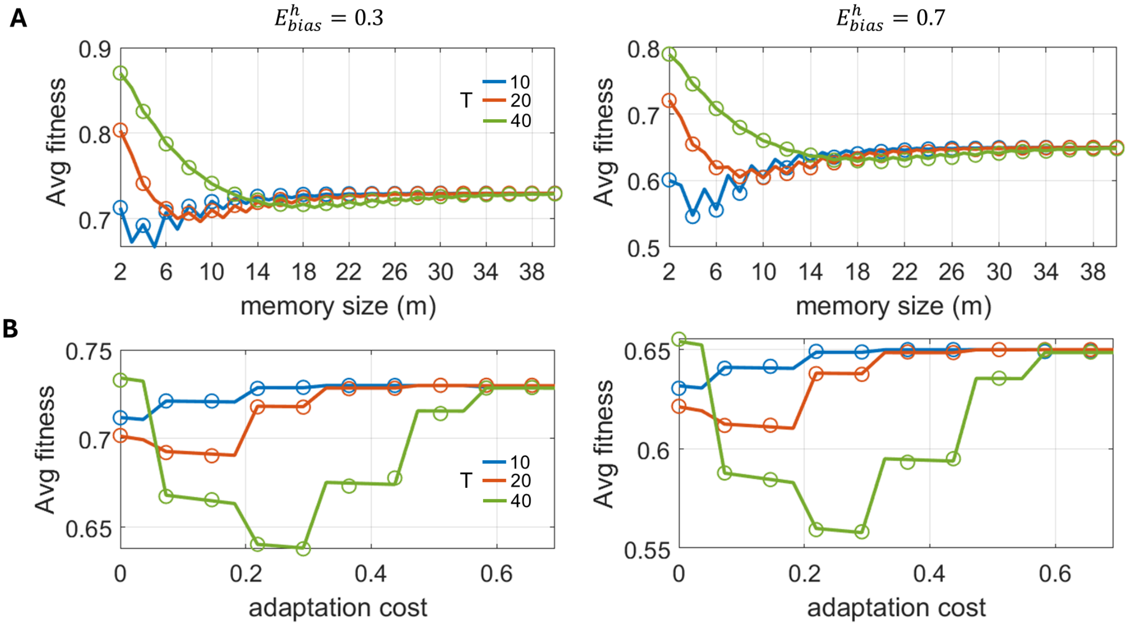


Figure S2: **Effects of memory size and adaptation cost on fitness for oscillating environments.** Here, $E^h_{bias}$ denotes the fraction of one environmental oscillation in the $E^h$ state, with results given for (left) $E^h_{bias} = 0.3$ generating environments that spend more time in the $E^l$ state, and (right) $E^h_{bias} = 0.7$ generating environments that spend more time in the $E^h$ state. Numerical validation of the analytical quantification is shown by circles, '$o$' (SI section S8). In panel A, adaptation cost $c_{lh} = c_{hl} = 0$; and in panel B, memory size, $m = 11$. The fitness values for (mis-)matching environment and cell state are $r_h^- = 0.3$, $r_h^+ = 0.8$, $r_l^- = 0.1$, and $r_l^+ = 1$.

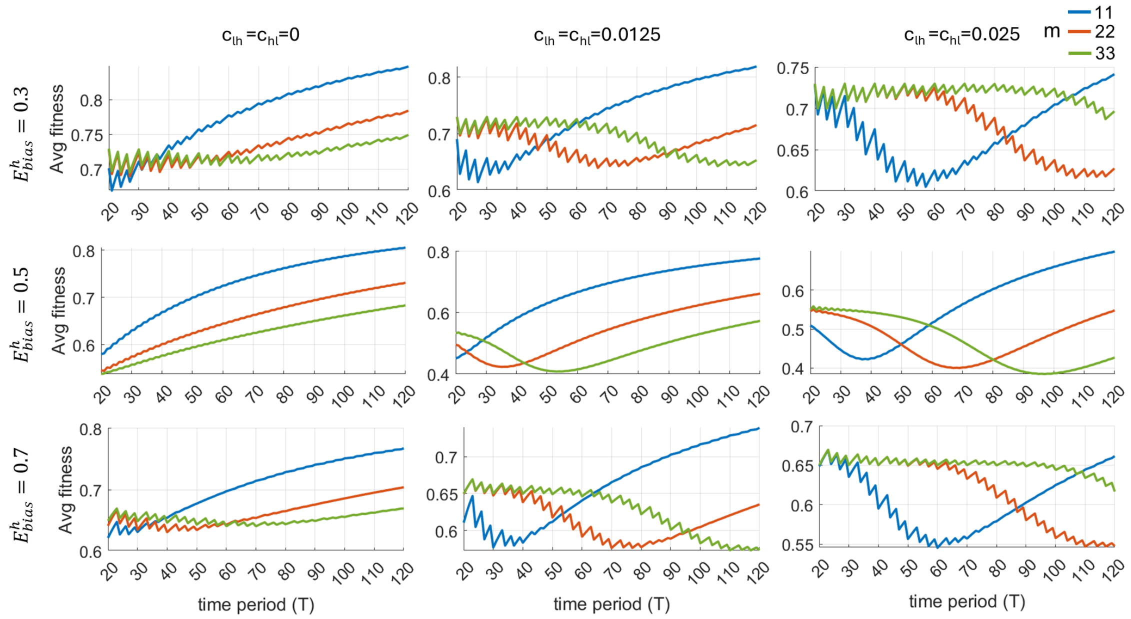


Figure S3: **Average fitness as a function of environmental oscillation period for variable environmental bias and adaptation costs.** The results are obtained from analytical quantification (SI section S8). The fitness values for (mis-)matching environment and cell state are $r_h^- = 0.3$, $r_h^+ = 0.8$, $r_l^- = 0.1$, and $r_l^+ = 1$.

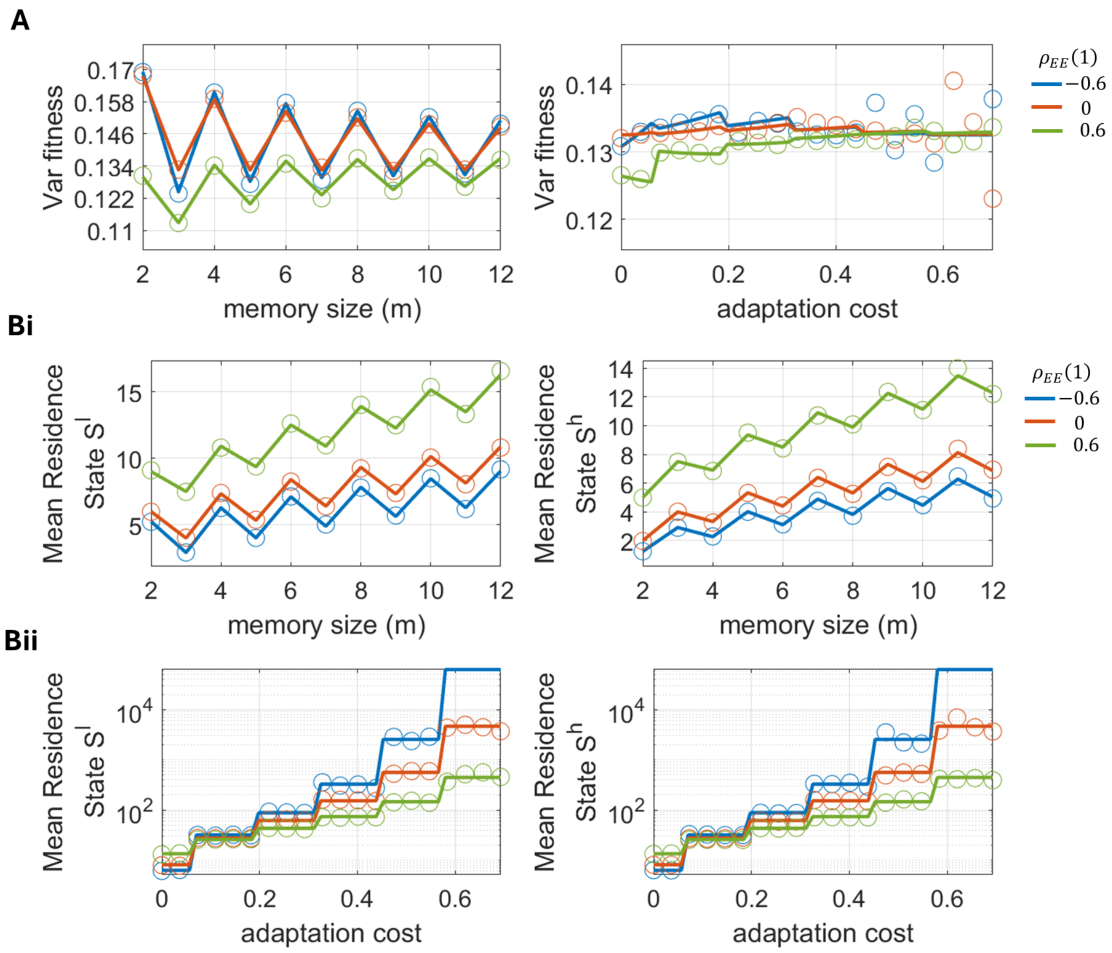


Figure S4: **Fitness variance and residence times in cell states as a function of memory size and adaptation cost in unbiased stochastic correlated environments.** (A) Variance in fitness is calculated as a function of memory size and adaptation cost. (B) Mean residence times in each state are given as a function of (i) memory size and (ii) adaptation cost. Numerical validation of the analytical quantification is shown by circles, '$o$' (SI section S8 and S11). While varying memory size, adaptation cost $c_{lh} = c_{hl} = 0$; and while varying adaptation cost, memory size $m = 11$. In all cases, $p = \mathbb{P}\left(S^n = S^h\right) = 0.5$ and the fitness values for (mis-)matching environment and cell state are $r_h^- = 0.3$, $r_h^+ = 0.8$, $r_l^- = 0.1$, and $r_l^+ = 1$.

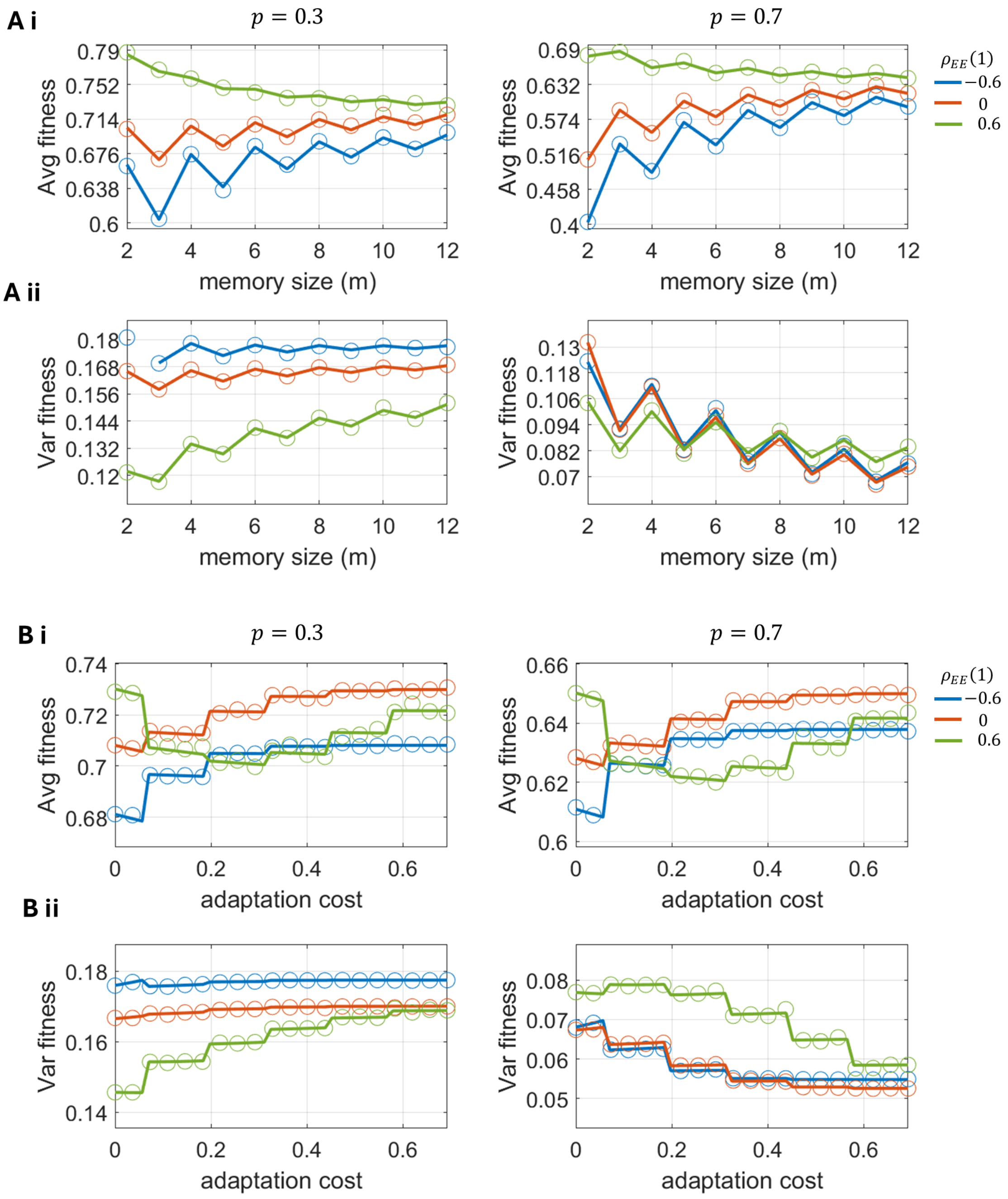


Figure S5: **Fitness mean and variance as a function of memory size and adaptation cost in biased stochastic correlated environments.**(i) Average fitness and (ii) variance in fitness values are given as a function of memory size (A) and adaptation cost (B) for stochastic environments biasing toward the $E^l$ state (left column) and $E^h$ state (right column). Here, $p = \mathbb{P}\left(S^n = S^h\right)$. Numerical validation of the analytical quantification is shown by circles, '$o$' (SI section S8). In panel A, adaptation cost was taken to be $c_{lh} = c_{hl} = 0$; in panel B, memory size is taken to be $m = 11$. In all cases $r_h^- = 0.3$, $r_h^+ = 0.8$, $r_l^- = 0.1$, and $r_l^+ = 1$.

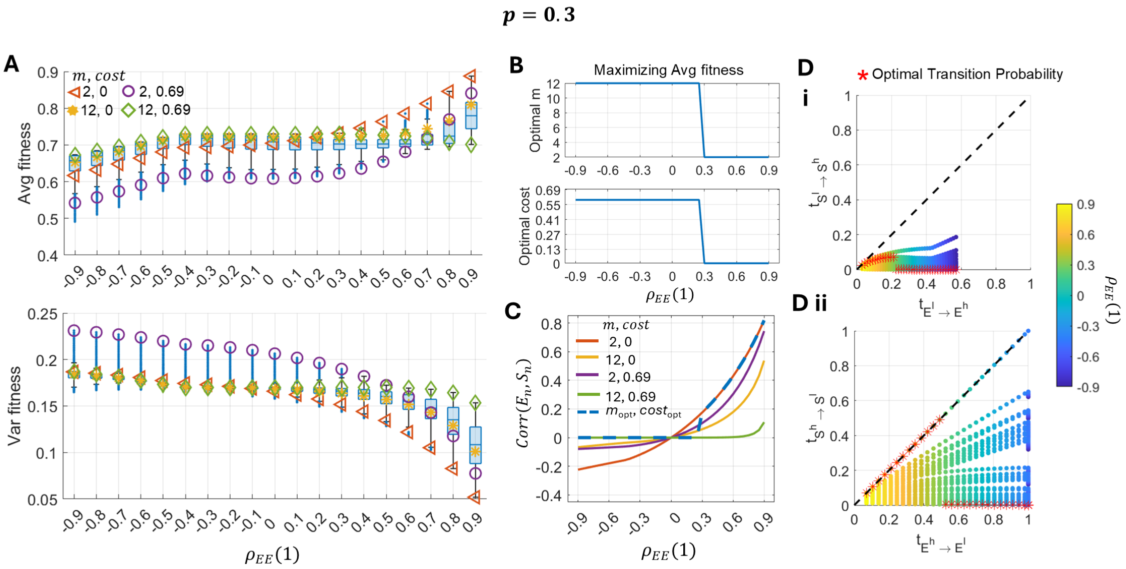


Figure S6: **Cellular adaptation to correlated stochastic environmental fluctuations biased for $E^l$ environment state. A)** Changes in the mean and variance of fitness with environmental correlation. The distributions for each correlation value are obtained by all possible combinations of $m \in [2, 12]$ and 100 values of adaptation cost linearly sampled from the range $c_{lh} = c_{hl} \in [0, 0.69]$. Average fitness trajectories from the extreme combination values $(m, c) \in \{\{2, 0\}, \{2, 0.69\}, \{12, 0\}, \{2, 0.69\}\}$ are highlighted with specific markers. **B)** Changes in the optimal memory size and cost that maximize average fitness for increasing environmental correlation. **C)** Cross-correlation between the current environment $(E_n)$ and cell state $(S_n)$ with increasing autocorrelation in the environment, $\rho_{EE}(1)$. The dashed blue curve highlights the correlation obtained from optimal $(m, c)$ obtained from panel B. **D)** Changes in the cell state transition probabilities (i) $t_{S^l \to S^h}$ and (ii) $t_{S^h \to S^l}$ with environmental transition probabilities, $t_{E^l \to E^h}$ and $t_{E^h \to E^l}$. $t_{E^l \to E^h} = p_1$ and $t_{E^h \to E^l} = p_2$ are obtained based on the fixed environmental correlation value (Eq. S1). Scatter points are combinations of transition probabilities obtained from all possible combinations of $m \in [2, 12]$ and 50 values of adaptation cost linearly sampled from the range $c_{lh} = c_{hl} \in [0, 0.69]$ with environmental correlation, $p_{EE}(1) \in [-0.9, 0.9]$ having a step size of 0.1. Optimal combinations of environment and cell state transition probabilities for increasing environmental correlation are denoted by astericks, '$*$'. In the above quantification, $r_h^- = 0.3$, $r_h^+ = 0.8$, $r_l^- = 0.1$ and $r_l^+ = 1$; and $\mathbb{P}(E_n = E^h) = p = 0.3$.; adaptation cost $(c_{lh} = c_{hl} = 0)$ and memory capacity $m = 11$, unless stated otherwise. The results are obtained from analytical quantification (SI section S8, S7, and S9).

$\boldsymbol{p = 0.7}$

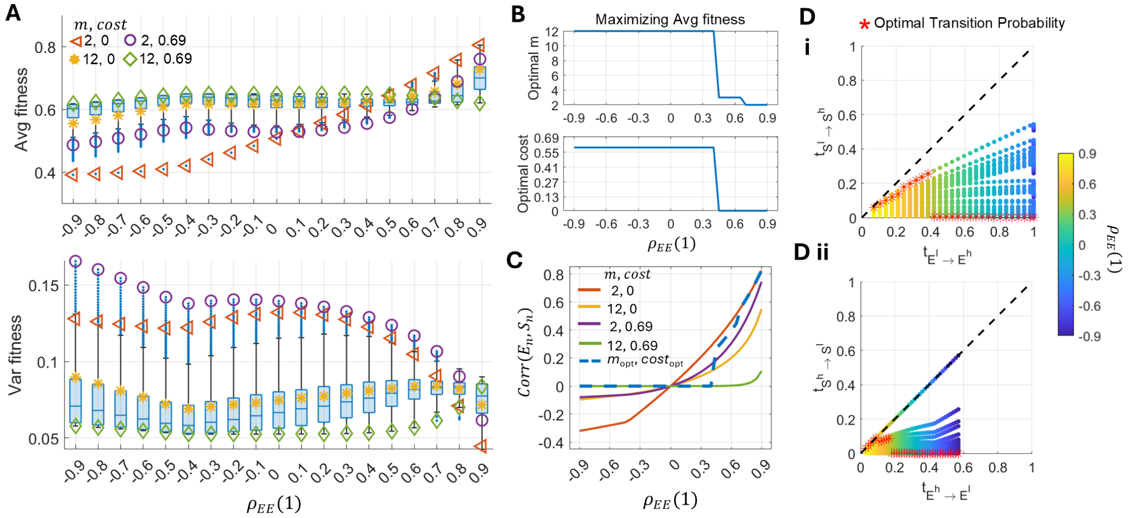

Figure S7: **Cellular adaptation to correlated stochastic environmental fluctuations biased for $E^h$ environment state. A)** Changes in the mean and variance of fitness with environmental correlation. The distributions for each correlation value are obtained by all possible combinations of $m \in [2, 12]$ and 100 values of adaptation cost linearly sampled from the range $c_{lh} = c_{hl} \in [0, 0.69]$. Average fitness trajectories from the extreme combination values $(m, c) \in \{\{2, 0\}, \{2, 0.69\}, \{12, 0\}, \{2, 0.69\}\}$ are highlighted with specific markers. **B)** Changes in the optimal memory size and cost that maximizes average fitness for increasing environmental correlation. **C)** Cross-correlation between the current environment $(E_n)$ and cell state $(S_n)$ with increasing autocorrelation in the environment, $\rho_{EE}(1)$. The dashed blue curve highlights the correlation obtained from optimal $(m, c)$ obtained from panel B. **D)** Changes in the cell state transition probabilities (i) $t_{S^l \to S^h}$ and (ii) $t_{S^h \to S^l}$ with environmental transition probabilities, $t_{E^l \to E^h}$ and $t_{E^h \to E^l}$. $t_{E^l \to E^h} = p_1$ and $t_{E^h \to E^l} = p_2$ are obtained based on the fixed environmental correlation value (Eq. S1). Scatter points are combinations of transition probabilities obtained from all possible combinations of $m \in [2, 12]$ and 50 values of adaptation cost linearly sampled from the range $c_{lh} = c_{hl} \in [0, 0.69]$ with environmental correlation, $p_{EE}(1) \in [-0.9, 0.9]$ having a step size of 0.1. Optimal combinations of environment and cell state transition probabilities for increasing environmental correlation are denoted by astericks, '$*$'. In the above quantification, $r_h^- = 0.3$, $r_h^+ = 0.8$, $r_l^- = 0.1$ and $r_l^+ = 1$; and $\mathbb{P}\left(E_n = E^h\right) = p = 0.7$.; adaptation cost $(c_{lh} = c_{hl} = 0)$ and memory capacity $m = 11$, unless stated otherwise. The results are obtained from analytical quantification (SI section S8, S7, and S9)

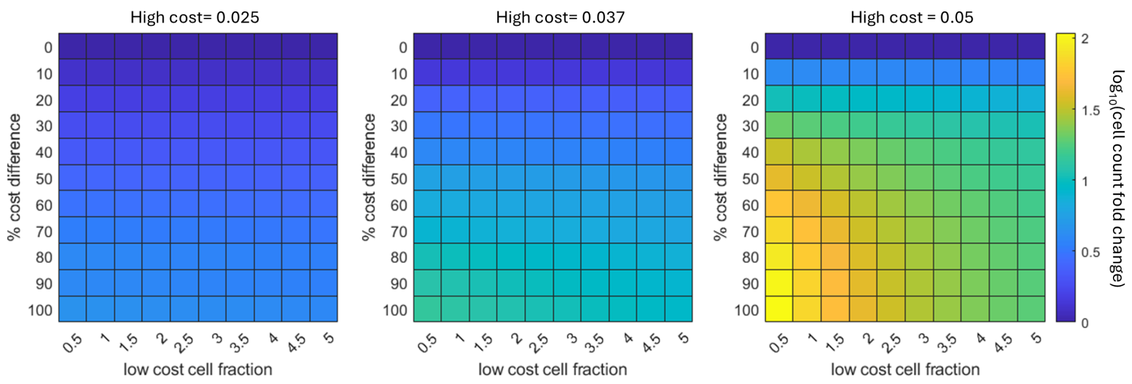


Figure S8: **Influence of phenotypic composition of heterogeneous population and adaptation cost differences between high- and low-cost subpopulations on population survival in cytotoxic environments.** The color bar shows the fold change in the total population size of the enriched low cost population compared to the heterogeneous population after seven days of continuous drug exposure. The analyses is done for fixing the high cost levels to 0.025, 0.037, and 0.05 and varying the low cost value as the % cost difference with respect to fixed high cost values. In the above, $r_l^+ = \frac{1}{20}h^{-1}$, $r_l^- = -\frac{1}{20}h^{-1}$, $r_h^+ = \frac{1}{100}h^{-1}$, and $r_h^- = 0\,h^{-1}$; and m = 40. The initial population size was $10^4$ cells, and the population grew in an $E^l$ environment ($p_1 = 0$) for 24h before switching environments at time 0.

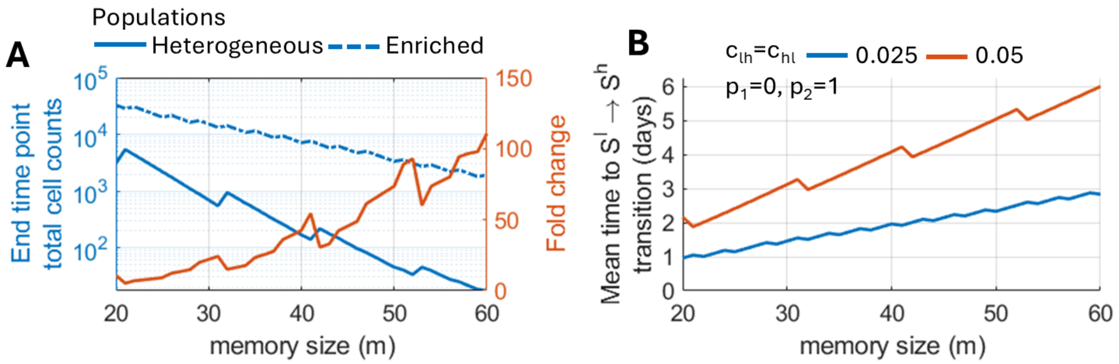


Figure S9: **Influence of memory size on population survival in cytotoxic environments. A)** Influence of memory size on the total population size achieved by day 14 on drug exposure for a (blue solid curve) heterogeneous population containing cells with low (0.04%) and high (99.96%) adaptation costs or (blue dashed curve) an population enriched with low cost cells (100%) The fold change in the total population size of the enriched population compared to the heterogeneous population is shown with a solid red line. **B)** Influence of memory size on mean adaptation time to the preferred cell state following a change in the environments for the low (0.025) and high (0.05) adaptation cost considered above. In the above, $r_l^+ = \frac{1}{20}h^{-1}$, $r_l^- = -\frac{1}{20}h^{-1}$, $r_h^+ = \frac{1}{100}h^{-1}$, and $r_h^- = 0\,h^{-1}$. The initial population size was $10^4$ cells, and the population grew in an $E^l$ environment ($p_1 = 0$) for 24h before switching environments at time 0.

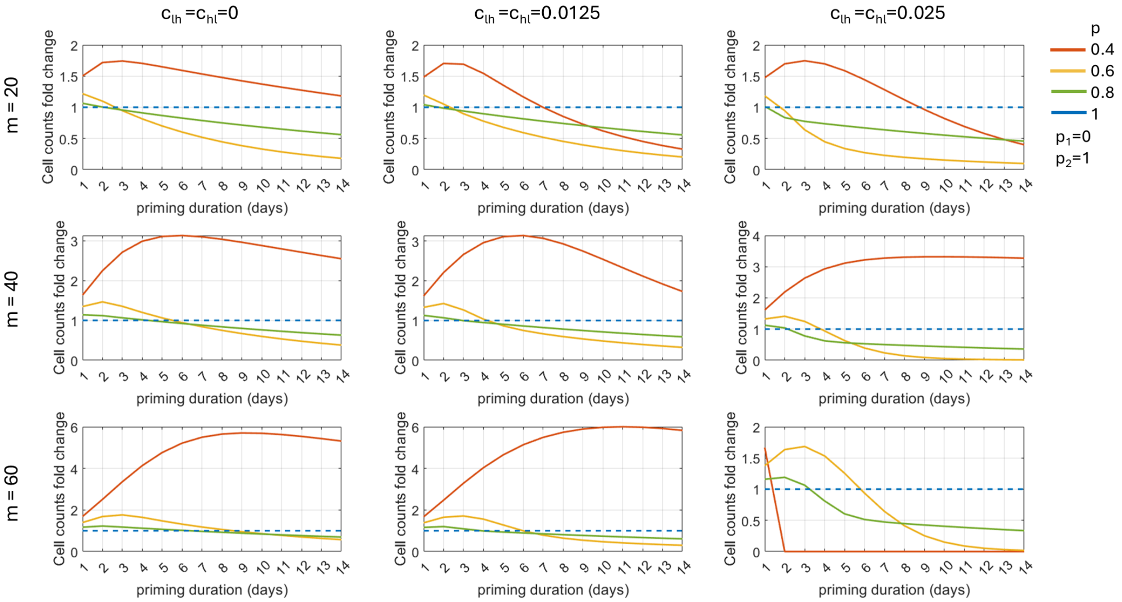


Figure S10: **Influence of memory size and adaptation cost on population-level response to lethal environments after priming with sub-lethal environmental exposures.** The y-axis denote the log10 fold change in mean cell counts at the end time point (day 14) for primed populations relative to treatment-naïve populations for increasing priming duration and variable priming environment ($p$).
